# The BPI-like TULIP domain proteins of *Drosophila melanogaster*: a novel class of candidate odorant transporters

**DOI:** 10.64898/2026.06.25.734463

**Authors:** Stéphane Dupas, Isabelle Chauvel, François Bousquet, Jérôme Cortot, Nicolas Kelle, Maud Bourgeois, Valentin Boichot, Aline Bonnotte, Laure Avoscan, Pierre-Yves Musso, Stéphane Fraichard, Loïc Briand, Fabrice Neiers, Jean-Philippe Charles

**Author notes:** Université Bourgogne Europe, Institut Agro Dijon, INRAE, UMR Agroécologie, Dijon, France.

## Abstract

TULIP (TUbular LIPid binding) domain proteins (TDPs) are found in all living organisms including bacteria. They have various documented functions, some of which clearly related to their intra- or extracellular lipid transfer activities. Extracellular, BPI-related TDPs of insects (B-TDPs, also known as Takeout-related proteins), are often found in chemosensory organs, but little is known regarding their exact location or how they could contribute to olfaction or gustation. We have surveyed and updated the full set of *Drosophila* B-TDPs and found that roughly 50% are overexpressed in chemosensory organs. Focusing on three genes clustered on the third chromosome, we provide evidence that at least one of the encoded proteins is secreted in the lymph cavity housing the dendrites of olfactory neurons. Biochemical data give support for a putative function of B-TDPs as odorant transporters, but loss-of-function analyses also hint to a potential role as a barrier against plant-emitted terpenoids.

## 1. Introduction

Lipid trafficking in eukaryotic cells relies not only on the vesicular pathway, but also on the activity of at least 27 families or superfamilies of Lipid Transfer Proteins (LTPs). Although structurally diverse, LTPs have in common a hydrophilic outer surface, and an inner hydrophobic pocket shielding lipid ligands from the surrounding aqueous environment (Chiapparino et al., 2016; Egea, 2021). The TULIP (TUbular LIPid binding) domain superfamily of LTPs was uncovered about 15 years ago by remote homology searches with profile HMM (Hidden Markov Models, (Kopec et al., 2011, 2010)). The TULIP domain is characterized by the presence of a long, more-or-less kinked α-helix wrapped by a twisted, anti-parallel ß meander, and forms an elongated hydrophobic pocket (Suppl Figure 1). TULIP domain proteins (TDPs) were found in all organisms surveyed so far, including monoderm and diderm bacteria, and despite an overall low degree of sequence identity (even among paralogs), the structure of the TULIP domain was remarkably conserved throughout evolution ((Kawano et al., 2018; Kopec et al., 2011, 2010; Kornmann et al., 2009; Levine, 2019; Wong and Levine, 2017), (Suppl Figure 1)).

The first ever described member of the TDP superfamily is the mammalian BPI protein (Bactericidal Permeability Increasing, (Beamer et al., 1997)), founder of the eponymous group of TDPs (BPI-like TDPs or B-TDPs), which are essentially secreted proteins composed of either a single TULIP domain, or two TULIP domains folded in a head-to-head orientation (also called pseudodimers, (Alva and Lupas, 2016; Wong and Levine, 2017), Suppl Figure 1). B-TDPs were shown to transport various lipid ligands, including the Lipid A moiety of LPS (LipoPolySaccharide, (Cordeiro et al., 2013; Gautron et al., 2011)), JH (Juvenile Hormone, a sesquiterpenoid hormone of insects, (Goodman, 1990; Prestwich et al., 1996)), or to shuttle cholesteryl-esters and phospholipids between lipoproteins (Masson et al., 2009; Weiss, 2003). In addition to these functions in defense against bacteria and lipid metabolism, clearly linked to their ability to bind lipids, there is also substantial evidence that B-TDPs have bacteriostatic effects as surfactants in the secretions of the upper airways (Bartlett et al., 2011).

Members of the SMP group (Synaptotagmin-like, Mitochondrial and lipid binding proteins) of TDPs, in contrast, possess additional motifs affixed to the TULIP domain (typically, at least a transmembrane domain), are intracellular and concentrated at membrane contact sites between various organelles, where they likely participate in lipid transfer between membranes (Kopec et al., 2010; Lee and Hong, 2006; Toulmay and Prinz, 2011). For example, SMP proteins of the yeast ERMES complex (ER–Mitochondria Encounter Structure) tether the endoplasmic reticulum and the mitochondrial outer membranes and were shown to facilitate phospholipid exchange between these organelles (Kawano et al., 2018; Kornmann et al., 2009).

Most studies in Insects have so far focused on B-TDPs, also called Takeout-like after th*e Drosophila* Takeout protein (CG11853 (To), (Sarov-Blat et al., 2000; So et al., 2000)). Phenotypic analyses of the To^[1]^ allele have demonstrated the involvement of the Takeout protein in various functions, often in relation with photoperiod, such as feeding and courtship behavior, locomotor activity, and longevity (Dauwalder et al., 2002; Meunier et al., 2007). The *Drosophila* B-TDP Daywake (CG2650) was shown to regulate circadian activity and to work as an "anti-siesta" factor (Villegas et al., 2024). In other insects, the best-known B-TDPs are Lepidopteran Juvenile Hormone binding proteins (JHBPs). JHBPs are high affinity hemolymph transporters for juvenile hormones (JHs), a family of linear sesquiterpenoids inhibiting metamorphosis and regulating sexual maturation in insects (Goodman, 1990; Prestwich et al., 1996).

In the last decades, a wealth of studies have reported the presence of high levels of B-TDP mRNAs or proteins in insect chemosensory organs (Bohbot and Vogt, 2005; Corcoran et al., 2015; Dauwalder et al., 2002; Fujikawa et al., 2006; Gu et al., 2011; Hamiaux et al., 2009; Justice et al., 2003; Levine, 2019; Rund et al., 2013; Saito et al., 2006; Sarov-Blat et al., 2000; Yoshizawa et al., 2011) suggesting that B-TDPs may be involved in chemical senses. Chemosensation in insects is accomplished by hair-like porous structures called sensillae. These cuticular projection are filled with an aqueous sensillar lymph secreted by accessory cells, bathing the dendrites of one or a few sensory neurons, which bear at their surface transmembrane receptors triggering (and sometimes inhibiting) neuron spiking upon binding specific ligands (Joseph and Carlson, 2015; Robertson, 2019; Schmidt and Benton, 2020).

In the course of million years of insect and plants coevolution, a rich and complex network of chemical communication has developed, allowing plants to discourage or kill unwanted herbivores with repellent or toxic molecules, attract natural predators and parasitoids of phytophagous species, or attract pollinators (Labandeira, 2013; Unsicker et al., 2009). Conversely, insects rely on the volatile organic compounds (VOCs) released by plants as cues to distinguish host from non-host plants, whether they need them for feeding, laying eggs or meeting mates for reproduction (Bruce, 2015; Bruce and Pickett, 2011; Xu and Turlings, 2018). Although chemically very diverse, the complex volatile blend secreted by plants, sometimes in copious amounts referred to as essential oils, is mainly composed of lipophilic compounds, and dominated by terpenoids (also called isoprenoids), a large family of molecules composed of one or more C5 isoprene units (Bakkali et al., 2008; Baldwin, 2010; De Groot and Schmidt, 2016; Dudareva et al., 2006; Maffei et al., 2011).

An important and still debated question is the nature of the mechanism allowing lipophilic odorants such as terpenoids to diffuse through the aqueous sensillar lymph and reach their cognate receptors in the membrane of olfactory dendrites. An early hypothesis is that these molecules may diffuse through the lymph, adsorbed at the surface of nanometric "pore tubules" attached at one end to sensillar pore openings, and establishing several contacts on the other side with dendritic membranes of olfactory neurons (Steinbrecht, 1997; Steinbrecht and Mueller, 1971). The characterization of a male antennal protein able to bind the female pheromone of the silk moth *Antherea polyphemus* (Vogt and Riddiford, 1981) led to an alternative hypothesis, suggesting that lipophilic odorants may be transported through the sensillar lymph by specialized transporter proteins. Three candidate protein families, named Odorant Binding Proteins (OBPs), Chemosensory Proteins (CSPs) and Nieman-Pick proteins, type C2 (NPC2) have since been identified. They are present in high concentrations in the sensillar lymph, and possess a binding pocket fit for the transport of small hydrophobic ligands (Pelosi et al., 2014, 2018a; Rihani et al., 2021; Schmidt and Benton, 2020).

In this paper, we provide an updated survey of *Drosophila melanogaster* B-TDP genes and report their widespread expression in chemosensory organs of the adult head. We examined in more detail the genes clustered at the 82E locus. We show that both CG14661 and CG2016 are expressed in the trichogen accessory cells of the maxillary palp and bring evidence that CG14661 is secreted in the sensillar lymph. We hypothesized, based on existing cues for ligand binding by insect B-TDPs, that their elongated hydrophobic pocket may allow the solubilization of linear terpenoid odorants in the sensillar lymph. Flies harboring deletions at the 82E locus, however, were not impaired in an olfactory assay towards a panel of ubiquitous terpenoid plant volatiles, but surprisingly, showed signs of hyperesthesia. Together with evidence gathered in other insects, our data show that B-TDPs qualify as a novel family of putative odorant transporters, although their actual function in chemosensory organs remains uncertain.

## 2. Results

### *Drosophila* BPI-like TULIP domain proteins genes

TULIP domain proteins are essentially subdivided into SMP-TDPs (Synaptotagmin-like, Mitochondrial and lipid binding proteins), and B-TDPs (BPI-like; Bactericidal Permeability Increasing-like) (Wong and Levine, 2017)). Based on Interpro entry IPR031468 (Synaptotagmin-like mitochondrial-lipid-binding domain), the *Drosophila* genome contains at least three SMP-TDP domain protein genes (CG6643/Esyt2, CG10362/Pdzd8 and CG43783) (Blum et al., 2025; Gramates et al., 2022), that were not investigated in this paper. We focused instead on B-TDPs, motivated by their documented expression in chemosensory organs of several insects. The *Drosophila* genome contains 30 B-TDP genes, associated with Interpro entries IPR038606 (Takeout superfamily) and IPR010562 (Haemolymph juvenile hormone binding) (Blum et al., 2025). They are organized in seven distinct clusters containing two to five genes, and seven isolated genes (Figure 1A). The intergenic distance within clusters ranges from -86 pb (CG10264 and CG10407 are overlapping) to 5549 bp (between CG14259 and CG17189), with an average of *ca.* 1.4 kb (Suppl Table 2). The five B-TDPs genes clustered at 34D, together with a sixth, unrelated gene (CG33307) are located within the first large intron (54857 bp) of a large mRNA of the Centaurin-gamma 1A gene (CENG1A-RB), and are thus "nested" genes (Gibson et al., 2005).

**Figure 1:**
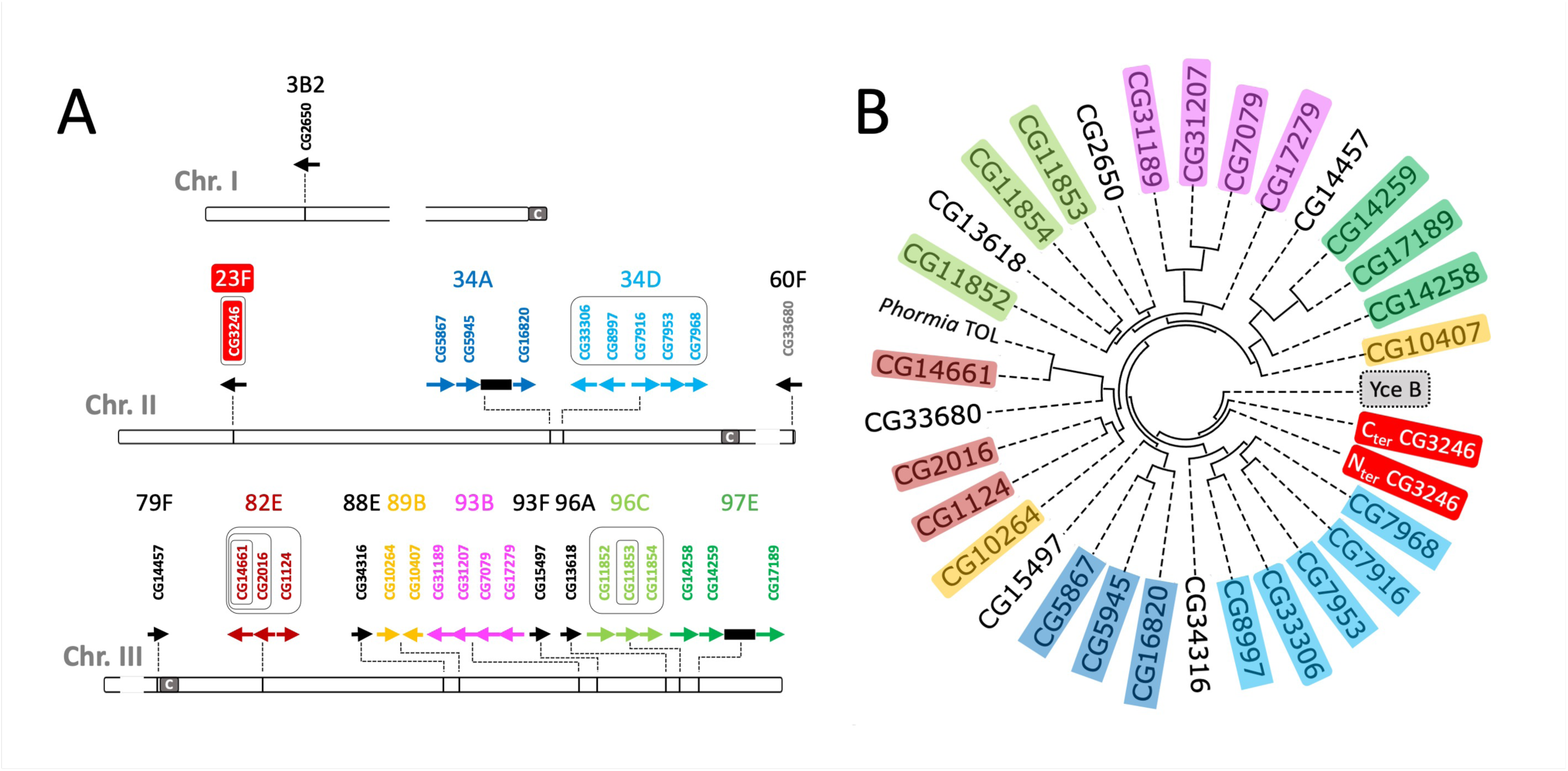
Drosophila melanogaster B-TDP genes. **A:** Twenty-three *D. melanogaster* B-TDP genes are organized in 7 distinct clusters (colored arrows and text), while the remaining seven genes are isolated (black arrows, black or white text). The gene written with white text on a red background (CG3246) encodes a protein with two TULIP domains (see text and Suppl Figure 1). CG33680 (grey text) is a pseudogene (see text and Suppl Figure 6). Each box with a thin black outline corresponds to a distinct fly line for which the boxed gene(s) or cluster was deleted by Crispr-Cas9. Arrows show the direction of transcription. The approximate cytology is indicated above each gene/cluster. The centromere (C) of each chromosome is shown as a grey rectangle. The two black rectangles correspond to non-TDP genes within TDP genes clusters. **B:** Structure based phylogenetic tree of *D. melanogaster* B-TDP genes. The tree was calculated using Alphafold structures ((Jumper et, al. 2021), Suppl Figure 1), with the open source tool Foldtree https://github.com/DessimozLab/fold_tree based on Foldseek’s “local structural alphabet” approach (Moi et al., 2023; van Kempen et al., 2024). Note that physically clustered genes are also generally closer paralogs. The two TULIP domains of CG3246 were analyzed separately, and cluster away from single TULIP domain proteins, with the bacterial representative TDP YceB used as an outgroop (Wong and Levine, 2017). The *Phormia regina* TOL protein was previously shown to be orthologous to the CG14661 protein (Fujikawa et al., 2006).

*Drosophila* B-TDPs are all predicted to be potentially secreted proteins, except CG33680 (Suppl Figure 5). The absence of a signal peptide in CG33680 is only apparent, as this gene presents a point mutation in the ATG start codon, and a 17 bp deletion in its coding sequence (Suppl Figure 6). In addition, modENCODE Temporal Expression Profile data suggest extremely low levels or no expression in all tissues examined (Gramates et al., 2022), consistently with our RT-qPCR analyses (Figure 3). CG33680 therefore qualifies as a pseudogene.

As predicted by Alphafold, all proteins share the typical TULIP fold, with a single protein (CG3246) having two TULIP domains folded in a head-to-head orientation (Suppl Figure 1, (Jumper et al., 2021)). Pairwise identities vary from about 30% for proteins belonging to the same clusters to as low as about 10% in proteins from distinct clusters (based on sequence alignments performed with Clustal Omega (Suppl Figure 7, (Madeira et al., 2024)), a fairly low percentage known as the "twilight zone", where it is difficult to infer phylogenies from protein sequences (Himmel et al., 2023; Rost, 1997). We drew a phylogenetic tree using the Treefold tool, a structure-based phylogeny reconstruction approach relying on the fact that 3D structures are more conserved than primary amino-acids sequences (Illergård et al., 2009; Moi et al., 2023; van Kempen et al., 2024). As shown in Figure 1B, the phylogenetic tree closely reflects the cytology, with most cluster members grouping in the same clade. Remarkably, both the N and C terminal TULIP domains of CG3236 cluster away from single domain TDPs, together with the remotely related bacterial TULIP domain protein YceB (Figure 1B, (Wong and Levine, 2017)).

A phylogenetic analysis of *Drosophila melanogaster* and *Bombyx mori* (silkworm) full sets of B-TDPs, and *bona fide* JHBPs from three other Lepidopteran species (*Manduca sexta*, *Heliothis virescens* and *Galleria mellonella* (Fig 2 and Suppl Table 3)) shows that five out of the seven *Drosophila* clusters have well supported orthologs in *Bombyx*. One of *Bombyx* B-TDPs, referred to as *Bm* JHBP in Figure 2, clusters with JHBPs of other species, but has no *Drosophila* ortholog. JHBPs therefore constitute a Lepidopteran-specific branch of B-TDP’s, sharing a conserved disulfide bridge that is not found in other B-TDPs (Figure 2B).

**Figure 2:**
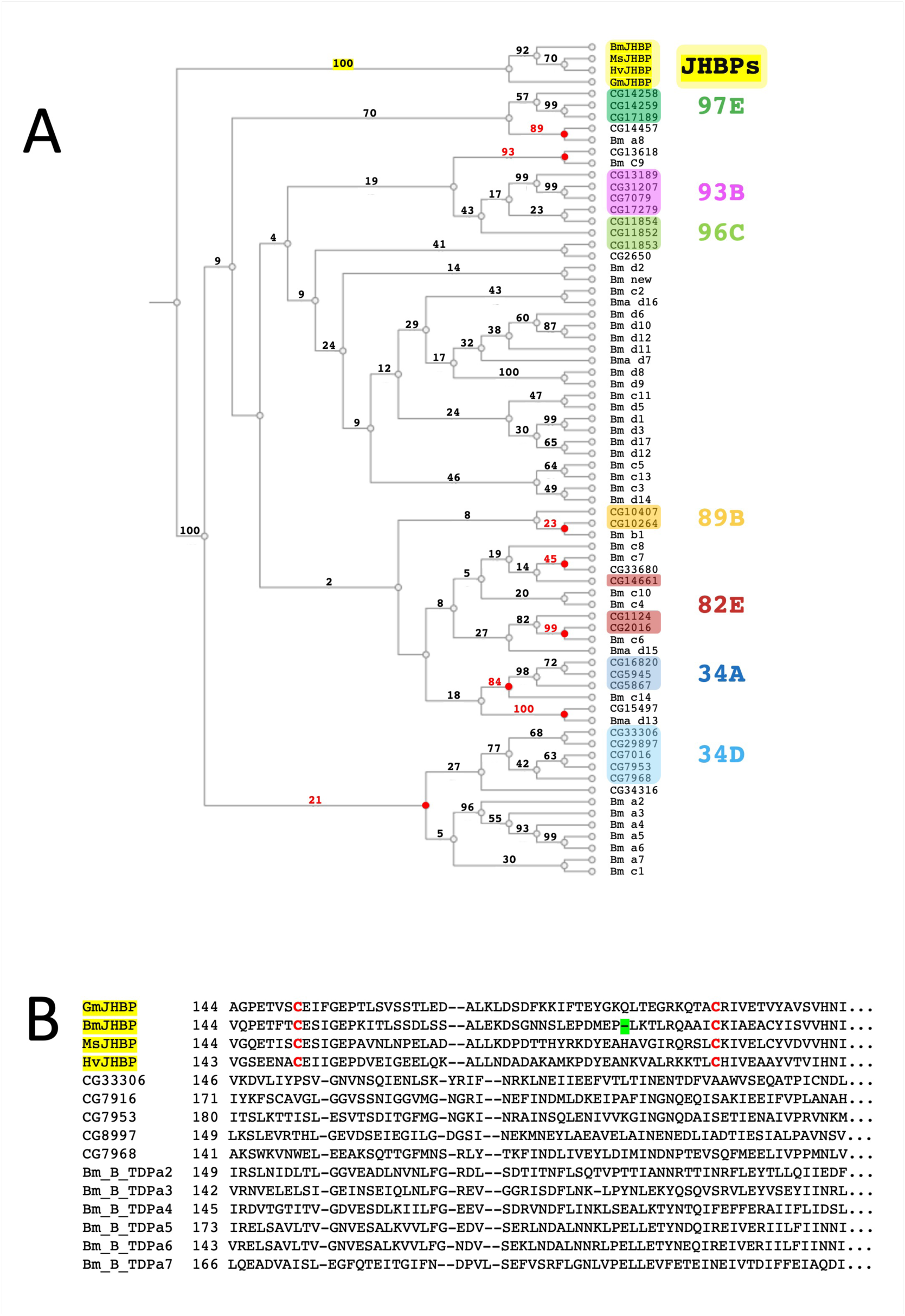
JHBPs are a diverged, Lepidopteran-specific clade of B_TDPs. A: Phylogenetic tree of single domain B_TDPs of Drosophila (CG #), Bombyx (Bm, Bma, (Li et al., 2016)), and four *bona fide* Lepidopteran Juvenile Hormone Binding Proteins (JHBPs) from *Bombyx mori* (BmJHBP, (BmJHBPd18(Li et al., 2016)), *Manduca sexta* (MsJHBP), *Heliothis virescens* (HvJHBP), and *Galleria melonella* (Gm JHBP). All proteins have a standard secretory Sec/SPI signal peptide, as predicted by Signal5P or Signal6P, which was removed before alignment (except Bm b1: no predicted signal peptide). Sequence alignment was performed with MAFFT version 7 (Katoh et al., 2019), using default parameters. Some *Bombyx mori* (Bm) proteins in the UniProt database appeared incomplete, perhaps owing to incorrect gene models. In those cases, the corresponding wild silkworm protein (*Bombyx mandarina,* Bma) was used for the alignement. Bm "new" corresponds to BMSK0013318 in SilkDB 3.0 (https://silkdb.bioinfotoolkits.net/main/species-info/-1) and was not reported earlier (Li et al., 2016). Two pseudodimeric *Bombyx* proteins with two TDP domains (BmJHBPd4 and BmJHBPa1 (Li et al., 2016) were not included in the alignment. The sequences of *Bombyx* B-TDPs used are detailed in Suppl Table 3. Phylogeny was inferred using the Neighbour-Joining method (Saitou and Nei, 1987), with 1000 bootstraps re-sampling. Numbers on the branches are the percentage of bootstraps supporting each corresponding node. Nodes highlighted in red indicate likely *Drosophila* and *Bombyx* orthologous proteins, supported by bootstrap values > 20%. B: Evidence for a conserved disulfide bridge in the C-terminal half of JHBPs. The alignment is an excerpt from the alignment used to infer the tree in (A). The two cysteine residues in the disulfide bridge linking the long C-terminal alpha helix (BmJHBP ⍺3 (Suzuki et al., 2011), GmJHBP ⍺4 (Kolodziejczyk et al., 2008)) to the ß meander (see Suppl Figure 1) are highlighted in red. The position of these cystein residues is almost perfectly conserved (with a single gap (green) in the BmJHBP protein relative to other JHBPs).

### Expression of *Drosophila* B-TDPs genes in chemosensory organs

We next sought to examine the expression of B-TDP genes in *Drosophila* main head chemosensory organs. RNAs extracted from olfactory organs (antennae and maxillary palps) and gustatory (labellae of the proboscis) organs of male or female heads were analyzed by RT-qPCR and compared with RNAs extracted from thoraces and abdomens from which wings and legs (also carrying chemosensory organs), were removed. As shown in Figure 3, roughly half of *Drosophila* B-TDPs genes are expressed at higher levels in chemosensory organs, notably CG2650 (Daywake) on the first chromosome, and those clustered at 34A, 82E, 93B, 96C and 97E (Figure 3A,B-D,E). This trend was more pronounced for female flies (Figure 3A,B). Expression ratios of genes belonging to the same cluster are in general remarkably similar, suggesting the existence of a mechanism of concerted regulation. In contrast, genes of the 34D cluster were significantly overexpressed in body samples. CG17279, in the 93B cluster, appeared highly specifically expressed in the olfactory system (arrowhead in Figure 3A-C,D-F), and was further investigated, but neither *in situ* hybridization, nor Western-blotting or immunocytochemistry with the M2 antibody (in fly lines with a Crispr-Cas9 edited CG17279 3XFLAG epitope terminus flagged) allowed us to corroborate this finding. One possible reason for this discrepancy is that the absolute expression level of this gene is very low or restricted to a low number of cells that we failed to detect. The expression ratios in male *vs* female comparisons were similar in chemosensory organs (Figure 3G,H), but all genes were expressed at higher levels in male bodies (Figure 3I).

**Figure 3:**
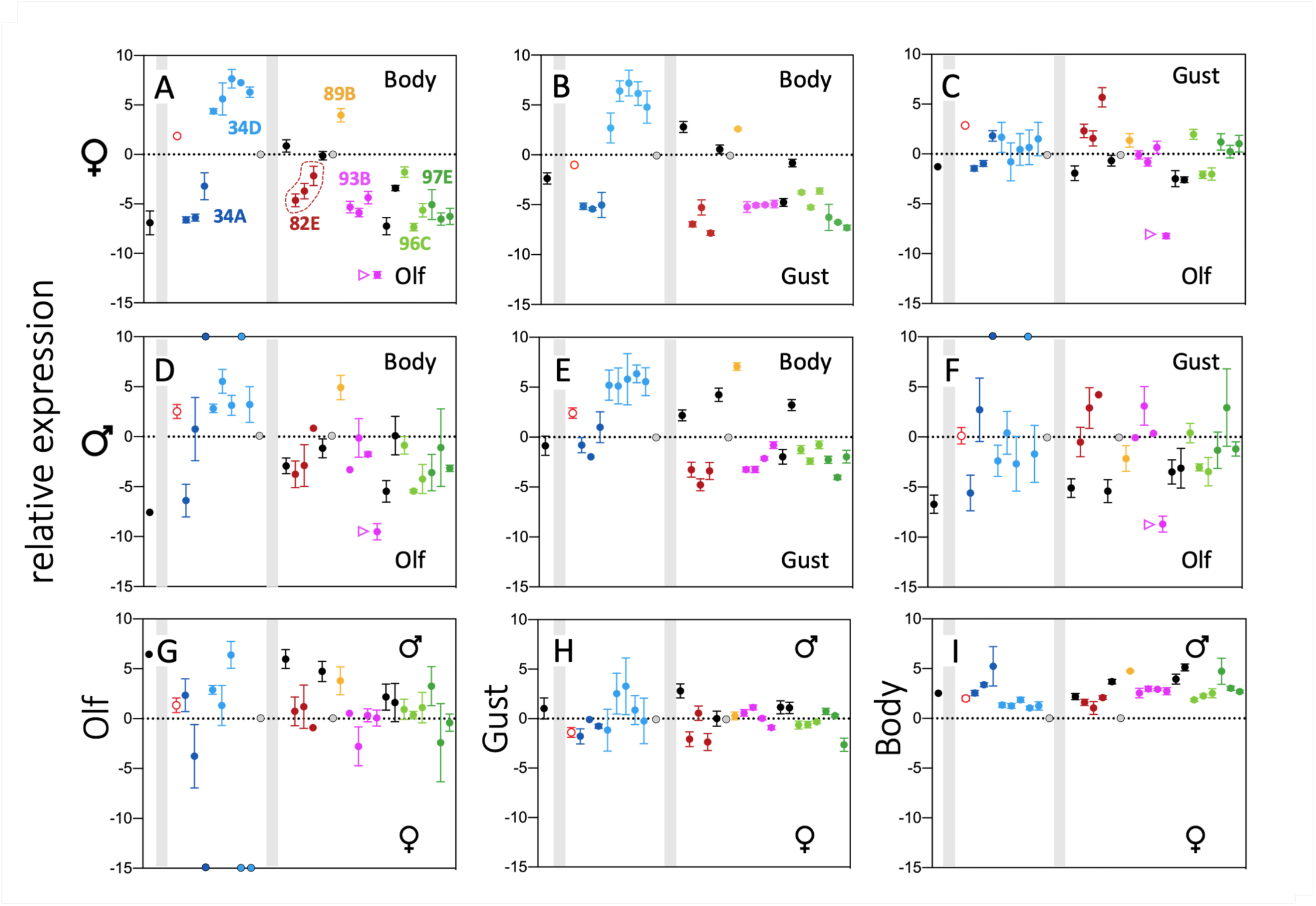
Expression of *Drosophila melanogaster* TDP genes in chemosensory organs. The relative expression of TDP genes in olfactory organs (Olf: antennae and maxillary palps), gustatory organs (Gust: Labellum), and in the body (thorax and abdomen, excluding the head) was analysed by RT-qPCR (the primers used are listed in Suppl Table 1). Genes are organized according to cytology, as shown in Figure 1, using the same color code. The seven B-TDP gene clusters are reminded in (**A**). Gene names are not recapitulated for clarity. The gene indicated with a magenta arrow (CG17279) was apparently highly specifically expressed in olfactory organs, but it could not be confirmed by other methods (see text). The three genes of the 82E cluster (dashed line) were selected for a detailed functional analysis. The CG33680 and CG10264 are represented as grey circles with a null ratio value, because their expression levels were not different from background in all samples (see also text). For some samples (CG7953 and CG7968 in the 34D cluster, and CG16820 in the 34A cluster), some ratio values are shown on either the top or the bottom axis, because the ΔCt value in male antennae was null or at background level. In each panel, a ΔΔCt value around zero correspond to a similar expression in the two types of samples being compared. Sample names at the top and bottom of each panel indicate the side of the panel corresponding to a higher expression in one sample type relative to the other. The vertical grey bars separate chromosomes I-III (Chromosome I on the left, with a single TDP gene (CG2650), chromosome II in the middle, chromosome III on the right side). Top row (A-C): Comparison of female samples. Middle row (D-F): comparison of male samples. Bottom row (G-I): comparison of male *vs* female samples. Relative expression levels are expressed as ΔΔCt (Differences in threshold cycles, (Livak and Schmittgen, 2001)) ± S.E.M (n= 2-6).

We set out to study the three genes clustered at 82E (CG14661, CG2016, CG1184), which are significantly overexpressed both in olfactory and gustatory organs (Figure 3 A,B-D,E). Importantly, CG14661 is the orthologue of the fly *Phormia regina* TOL protein, which was shown to be secreted into the sensillar lymph where, we hypothesized, it may be involved in transporting lipophilic odorant or sapid molecules such as volatile terpenoids (Figure 1B, Suppl Figure 1 (Fujikawa et al., 2006)).

### Localization of Expression of 82E cluster TULIPs in chemosensory sensillae

To document the expression of the three genes clustered at the 82E locus, we used Minos-derived gene (transcriptional) trap lines (referred to below as Mi-14661, Mi-2016 and Mi-1184, see also Materials and Methods, (Li-Kroeger et al., 2018; Venken et al., 2011)), and by immunodetection of the CG14661 protein carrying the 3xFLAG epitope on its C-terminal end in a fly line engineered by Crispr-Cas9 and homologous recombination (referred below as 14661 3xFL, see also Material and Methods, and Suppl Figure 2, (Gratz et al., 2014)). The patterns appearing qualitatively similar in male and female flies, sex-specific singularities were not looked up, but cannot be ruled out. A schematic summary of the expression of the three B-TDP genes in the head sensory organs is shown in Figure 4 M.

**Figure 4:**
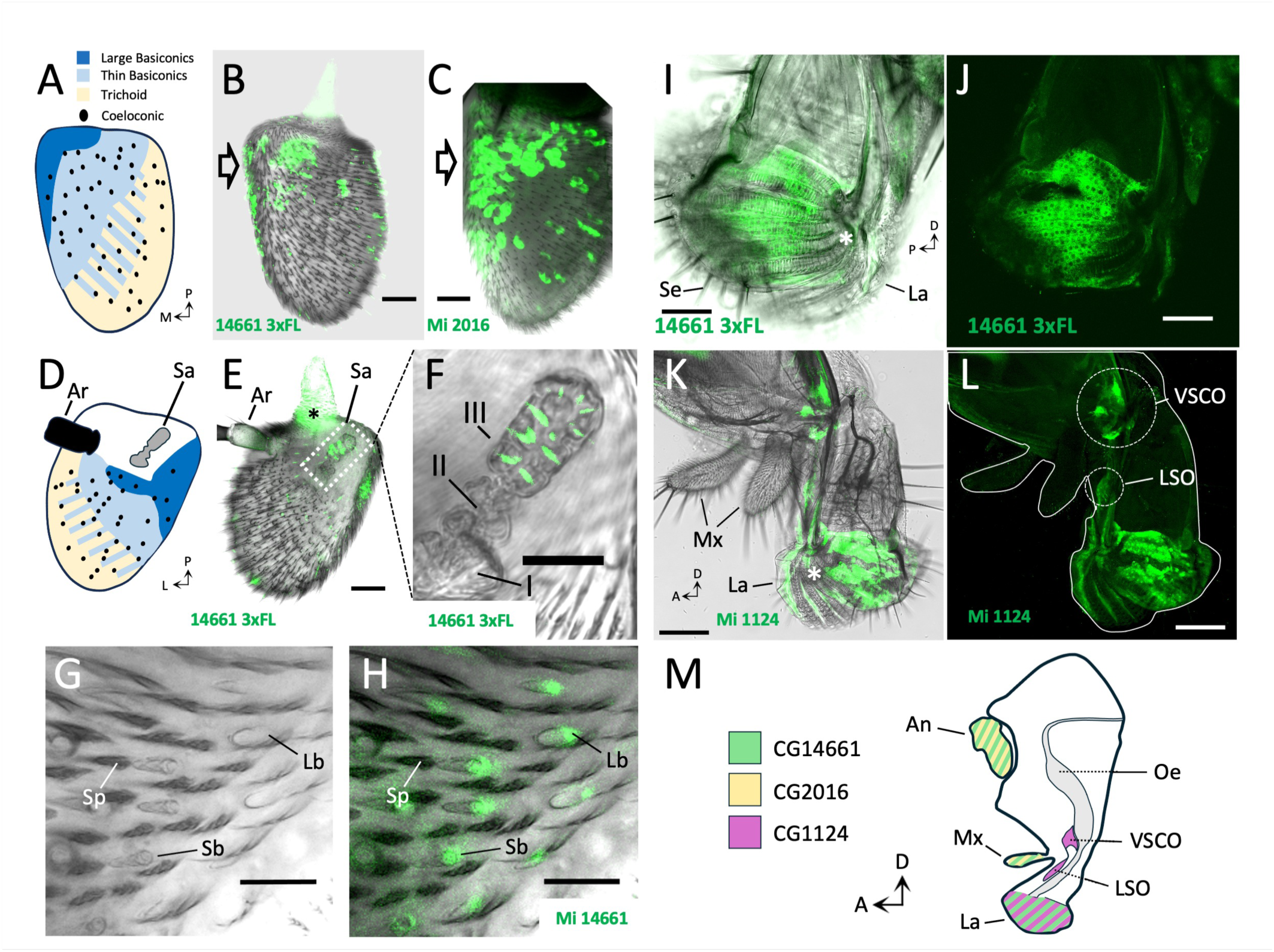
Expression of the 82E cluster genes in the antennae and proboscis of *D. melanogaster* antennae. The C-terminally flagged CG14661 (14661 3xFLAG) was detected with an anti-FLAG primary antibody (see Material and Methods). Gene expression was also documented in fly lines harboring either a Minos gene trap (Mi14661, Mi 2016, Mi1124; MiMICs, (Venken et al., 2011), (Li-Kroeger et al., 2018), see Material and Methods). (**A**) and (**D**), respectively showing repartition of sensillae on the anterior and posterior antennal surfaces, were redrawn from (Martin et al., 2013). **A-C**: Both CG14661 (**B**) and CG2016 (**C**) are expressed in the third antennal segment (funiculus), predominantly in a proximo-medial area of the anterior surface (large open arrows). **D-F**: CG14661 is also expressed in sensillae of the sacculus (Sa), a pit organ opening on the posterior surface of the third antennal segment (**F**: magnification of boxed area in (**E**), chambers of the sacculus are numbered I-III as in (Shanbhag et al., 1995)). **G**,**H**: In the medio-proximal area of the antenna (arrrows in (**B)** and (**C**)), cells expressing CG14661 reside underneath large (LB) and small (SB) basiconic sensillae, but not under noninervated cuticular spicules (Sp). In the proboscis (**I-L)**, CG14661 is expressed in cells lining pseudotracheae ((**I**) white asterisk), in the distal part of the proboscis (labellum, (La)), but is not associated to the gustatory sensillae (Se). **K**,**L**: CG1124 is expressed in cells lining pseudotracheae (white asterisk), and in two distinct areas next to the alimentary canal, coinciding with the labral sense organ (LSO) and the ventral cibarial sense organ (VSCO). **M**: A schematic summary of the expression of the three BTP genes clustered at 82E in the *Drosophila* head (outline redrawn from (Kendroud et al., 2018)). An: antenna, La: labellum Mx: maxillary palp. The expression in the maxillary palp (Mx) is reported in Figure 5. (**H**, **I**, **K**) are overlays of corresponding fluorescent and bright field images. The black asterisk in (**E**) indicates a specific signal in the hinge of the funiculus that could not be identified. Thin black arrows indicate orientation (A: anterior, D: dorsal, L: lateral, M: medial, P: proximal (**A,D**) or posterior (**I**)). Scale bars: **B**,**C**,**F** = 20µM; **E** = 15 µM; **G**,**H** = 10 µM; **I,J** = 50 µM; **K,L** = 100 µM.

#### Antennae

On the anterior surface of the third antennal segment, 14661 3xFL and Mi-2016 are detected primarily in a proximo-medial area populated by basiconic sensillae (Figure 4A; (Shanbhag, 1999; Stocker, 1994)), although with distinct patterns (Figure 4B,C). Mi-14661 expression (a gene trap tagging the cells expressing CG14661, rather than the localization of the protein itself) is seen underneath large and small basiconic sensillae (Figure 4G,H), but we could not ascertain that it is strictly restricted to this type of sensillae (and not present in some trichoid or coeloconic sensillae). The MI-2016 signal is more diffuse and did not allow us to correlate it with a specific sensillar type. It is however similar to the CG14661 pattern, and therefore likely essentially restricted to basiconic sensillae. The expression pattern of CG14661 on the posterior side of the antennae was less conspicuous, yet also mostly localized in a proximal area populated with basiconic sensillae (Figure 4D-F). In addition, the 14661 3xFL protein was detected in the sacculus, a multichambered pit organ located on the posterior face of the antenna, where it co-localizes, in the most proximal chamber III, in elongated structures likely corresponding to the shafts of grooved sensillae (Figure 4F, (Shanbhag et al., 1995)). The expression of CG2016 on the posterior side of the antenna was not recorded. CG1184 appears to be not expressed in the antenna.

#### Proboscis

A significant expression of CG14661 and CG1184, but not CG2016, was also observed in the proboscis, although these patterns were not associated with the large hair like gustatory sensillae of the labium (Figure 4I-L, Suppl Figure 8A,B). Rather, both genes were expressed in large, epidermal-looking cells lying underneath the cuticle of the pseudotracheae (food canals, white asterisks in Figure 4 I,K). This region also bears tiny taste sensillae (the taste pegs (Shanbhag et al., 2001)), in some of which CG14661 may also be expressed (Suppl Fig 8C-E). CG1184 is also expressed in two distinct areas along the alimentary canal, whose locations correspond to the labral sensory organ (LSO) and the ventral cibarial sense organ (VCSO), both of which are known to house gustatory sensillae (Figure 4I,J, (Nayak and Naresh Singh, 1983)).

#### Maxillary palp

Both CG14661 and CG2016 are expressed in sensillae of the dorsal surface of the maxillary palp, which are all of the basiconic type (Naresh Singh and Nayak, 1985), with highly similar patterns (Figure 5A,B), while the Mi-1124 gene trap was not detectable. Like all B-TDPs, CG14661 has an N-terminal typical Sec/SP1 secretory signal (Suppl Figure 5). Unsurprisingly, the 14661 3xFL protein could be detected within the shaft of basiconic sensillae (Figure 5C), suggesting that it is secreted in the sensillar lymph. Co-staining with the neuronal marker elav shows that CG14661 expression pattern does not match with neurons (Suppl Figure 8F-I). To determine in which type of sensillar accessory cell the CG144661 gene is expressed (thecogen, trichogen or tormogen; see Figure 5I; (Shanbhag et al., 2000)), we first compared the location of the 14661 3xFL protein with the Mi 14661 gene trap signal, which reveals the outline of the CG14661 expressing cells (Figure 5D). The bulk of flagged CG14661 protein is obviously extracellular and concentrated, apically, in the sensillar lymph (Figure 5D, I and Suppl movies 1-3). The cells expressing CG14661 have a complex shape, with a large internal lacuna that corresponds to the location of an ensheathed thecogen accessory cell and the dendrites of olfactory neurons (see below, and schematic representation in Figure 5I). In the cytoplasm, large, labelled granules are also observed (Figure 5E). We next compared the location of the 14661 3xFL protein and the MI 2016 signal (Figure 5 F, Suppl Movies 4-6)). These lateral views show that CG2016 and CG14661 are expressed in the same cells, and that the large lacuna (top row) narrows down apically (bottom row) to a narrow channel (*ca.* 1µm in diameter, see also inset in Figure 5G). We next localized 14661 3xFL in a fly line expressing a NompA:GFP fusion protein under the regulatory sequences of the NompA gene. The NompA protein is expressed in thecogen cells and contributes to an attachment site for dendrites of sensory neurons at the bottom of the sensillar shaft (Chung et al., 2001). As shown in Figure 5G, NompA:GFP is detected at the base of sensillar shafts, within the sensillar lymph, where the 14661 3xFL protein is concentrated. This observation confirms that the apical channel observed within CG14661 expressing cells (Inset in Figure 5G, filled white arrowheads in bottom rows of Figure 5D and 5F, Figure 5I), corresponds to the passageway of the sensory dendrites and their ensheathing thecogen cell through the trichogen cell (Figure 5I). Finally, we imaged the 14661 3xFL protein in a fly line expressing GFP under the control of the Ase5 enhancer, specifically in the tormogen cell (Barolo et al., 2000). As shown in Figure 4H, the 14661 3xFL signal is surrounded by the tormogen cell, confirming that CG14661 is expressed in trichogen cells (see also Suppl movies 7-9, and Figure 5I).

**Figure 5:**
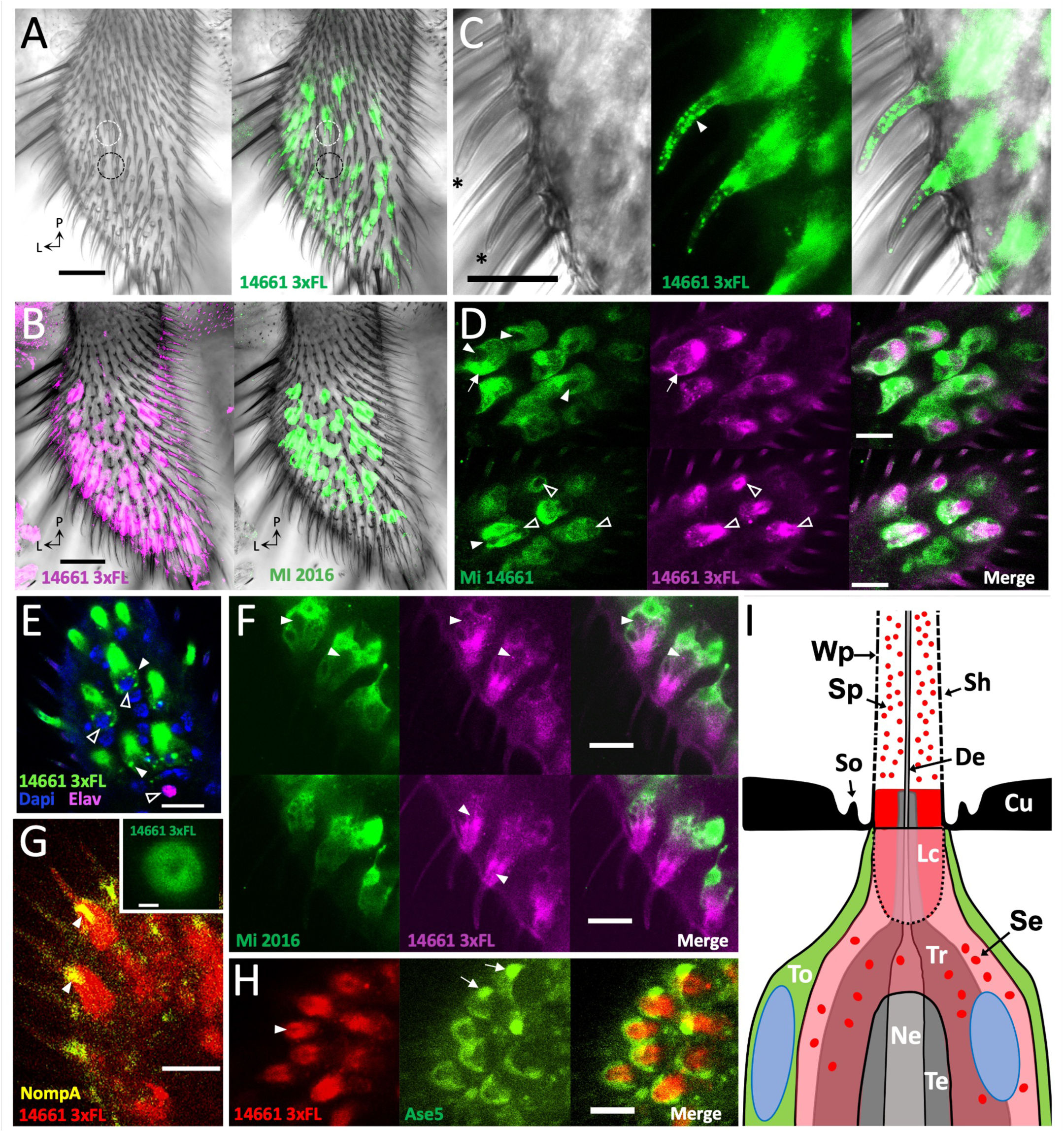
CG14661 and CG2016 are expressed in the trichogen cell of olfactory sensillae basiconicae of the maxillary palp. The expression of CG14661 and CG2016 was documented as described in the caption of Figure 4. A fly line driving the expression of GFP under the Ase5 regulatory element was used to image the tormogen cell ((**G**), (Barolo et al., 2000); Bloomington # 58449). The NompA protein, specifically located at the base of the sensillae in thecogen cells, was detected with an anti-GFP antibody in a drosophila line expressing a NompA::GFP fusion protein ((**H**), (Chung et al., 2001); Bloomington # 42694). The neuron nuclei were visualized with antibody against the neuronal marker elav. **A**: The CG14661 3xFLAG protein is primarily observed in a lateral distal area of the dorsal surface of the maxillary palp. It is found underneath a subset of basiconic olfactory bristles (translucent looking, white circle), but is not associated with uninervated trichomes (darker looking, black circle). **B**: Expression of CG14661 (left, 14661 3xFL)), and CG2016 (right, Mi-2016) in the same maxillary palp. Note the high similarity of the two patterns. **C**: Close-up on two basiconic sensillae (asteriks) showing a strong signal both below the cuticle surface as well as in speckles within the sensillar shaft (white arrowhead). **D**: Comparison of the localization of the 14661 3xFL protein and that of the CG14661 gene trap Mi-14661. Confocal slices of the same cells were taken *ca.* 8 µM (top row), and 5 µM (bottom row) beneath the cuticle surface. Note the large open cavity in CG144661 expressing cells in the deeper slices (top row, filled arrowheads) and the concentration of the 14661 3XFL protein around in the area beneath the sensillar shaft. A few micrometers higher, towards the cuticle surface (bottom row), the open cavity narrows into a *ca.* 1 µM diameter channel (filled, white arrowhead). At least part of the 3XFLAG protein is secreted into a cup-shaped extracellular space, called the receptor lymph cavity, just underneath the sensillar shaft (open, white arrowheads; see also Suppl movies (1-3), and schematic representation in (I).) The white arrows point to nuclei, as seen by Dapi staining (not shown). Note that the EGFP reporter protein is concentrated in the nuclei (EGFP is known to diffuse freely in nuclei (Wei et al., 2003)). **E**: CG14661 3xFLAG is concentrated in conspicuous secretion vesicles (filled, white arrowheads) underneath the sensillar shaft, near the nucleus (open, white arrowheads, stained with Dapi). The presence of a neuron nucleus (elav positive) in a deeper plane indicates that most CG14661 expressing cells are not neurons. **F**: Comparison of the localization of the 14661 3xFL protein and that of the CG2016 gene trap Mi 2016. The two confocal slices (top and bottom rows) of the same cells are 1,5 µM apart. Note that CG14661 and CG2016 are expressed in the same cells (Merge). The pear-shaped cavity corresponding to the location of thecogen cell (Top row, filled, white arrowheads), narrows apically to the channel (Bottom row, filled, white arrowheads), through which the thecogen cell, ensheathing the dendrites of the olfactory neurons, reach the lumen of the shaft (see also Suppl movies (4-6), and schematic representation in (I). **G**: Immunolocalization of CG14661 3xFLAG and a NompA:GFP fusion protein. The NompA protein is secreted by the thecogen cell and specifically adressed to a tethering structure for dendrites at the base of sensillar shafts (filled, white arrowheads, (Chung et al., 2001)). Note that NompA is located at the base of the shaft, in a central position corresponding to the channel observed within the CG14661 3xFLAG signal (inset), which indicates that CG14661 expressing cells ensheath the thecogen cells. **H**: Comparison of the localization of CG14661 3xFLAG and tormogen cells (as revealed with the tormogen cell-specific gene trap GFP reporter Ase5). The CG14661 expressing cells are partially ensheathed by the tormogen cells (Merge, see also Suppl movies (**7-9**) and (**I**)). The filled white arrowhead points to the channel as in (**E**). Arrows indicate concentration of GFP reporter in nuclei, as seen by Dapi staining (not shown). Thin black arrows in (**A-C**) indicate orientation (L: lateral, P: proximal). Scale bars: **A, C** = 25µM; **B,D,G** = 10 µm, inset in **G** = 1 µm. I: Schematic representation of a basiconic sensilla. The data above suggest that the CG146613xFLAG protein is concentrated in secretory vesicles (Se) of the trichogen cell (Tr), secreted in the receptor lymph cavity (Lc) at the base of the sensillar shaft (Sh), and adressed to speckles (Sp) higher up in the shaft. The thecogen cell (Te) ensheathing dendrite(s) (De) of olfactory neuron(s) (Ne) are indirectly localized as a channel through the lymph cavity. The trichogen cell is itself surrounded by the tormogen cell (To). Cu: Cuticle; So (Socket ridge), W (Wall pore). Nuclei of the tormogen and trichogen cells are sketched as blue ovals. The extracellular lymph cavity is outlined with dots.

### Functional analysis

The native CG14661 protein, like its flagged version, is probably secreted in the sensillar lymph and present throughout the sensillar shaft, suggesting that it may transport lipophilic odorant molecules to odorant receptors present in the dendritic membranes of olfactory neurons. We therefore performed preliminary competition fluorescence assays (Pelosi et al., 2018b), using recombinant CG14661 to test its affinity for a panel of potential ligands (Suppl Figures 3A and Figure 6). These candidate molecules were chosen based on a few criteria. Terpenoids appeared plausible ligands because of their documented abundance in plants volatile fractions and essential oils, and their well-known importance in plant-insect interactions (Baldwin, 2010; Boncan et al., 2020; Bruce, 2015; Rosenkranz et al., 2021)). Among them, linear (acyclic) terpenoids seemed more likely candidates than cyclic (bulkier) terpenoids, since CG14661 and other B-TDPs harbor a rather narrow and elongated pocket and because there is substantial experimental data either showing or hinting to B-TDPs binding linear terpenoids (Goodman, 1990; Hamiaux et al., 2009; Prestwich et al., 1996; Sugahara et al., 2020; Wybrandt and Andersen, 2001). Finally, we chose linear terpenoids among the most commonly encountered in plant volatiles and essential oils, because it is well established that plant odor specificity does generally result not from plant specific volatiles but rather from the emission of ratio-specific blends of widespread molecules (Bruce et al., 2005; Bruce and Pickett, 2011). As a pre-requisite for competition assays, several lipophilic fluorescent probes were tested for ability to bind to recombinant CG14661-6xHis (Fig 6A). Sypro® Orange, a widely used lipophilic fluorescent protein dye (Kroeger et al., 2017; Steinberg et al., 1996; Wu et al., 2023) was strongly fluorescent and exhibited a significant shift in emission wavelength ("blue shift") when incubated with CG14661-6xHis, indicating that the probe was bound in an hydrophobic environment, presumably the typical TULIP pocket of CG14661 (Suppl Fig 1, (Pelosi et al., 2018c)). When used as competitors at a five-fold molar excess, seven out of the nine linear mono-and sesquiterpenes tested were more effective in displacing Sypro® Orange from CG14661-6xHis than from BSA, which like other serum albumins present several hydrophobic binding sites for various lipophilic ligand including fatty acids and terpenoids such as retinoids ((Figure 6B, green asterisks ; (Bhattacharya et al., 2000; Charbonneau and Tajmir-Riahi, 2010; Curry et al., 1998)). Non-terpenoid small volatiles (benzaldehyde, trans-2 hexenal) also were able to displace Sypro® Orange, although in a lesser extent, perhaps merely because of their hydrophobic nature and small size. The affinity of CG14661-6xHis for Sypro® Orange (Kd; dissociation constant) is approximately 1.7 µM, as determined by means of a ligand saturation assay (Figure 6C (top)). A five-fold molar excess of competitors was at most able to displace about 50% of bound Sypro® Orange (farnesol, Figure 6B), which translates into an approximate Kd of ∼3.8 µM for this sesquiterpenoid (See Material and Methods section, (Pelosi et al., 2018b)). This range of concentrations discouraged further efforts to determine the Kds of any of these terpenoids for CG14661-6xHis by a ligand saturation assay, because these lipophilic molecules typically arrange into micelles at low concentrations, impeding affinity estimations (Pelosi et al., 2018b; Turina et al., 2006). Finally, the presence of a hydrophobic patch at the surface of CG14661 (Figure 6C (bottom)) calls for caution as this region might serve as a binding site for either Sypro® Orange or competing ligands. Notwithstanding these considerations, these preliminary data suggest that CG14661 has a significant affinity and specificity for linear terpenoids.

**Figure 6:**
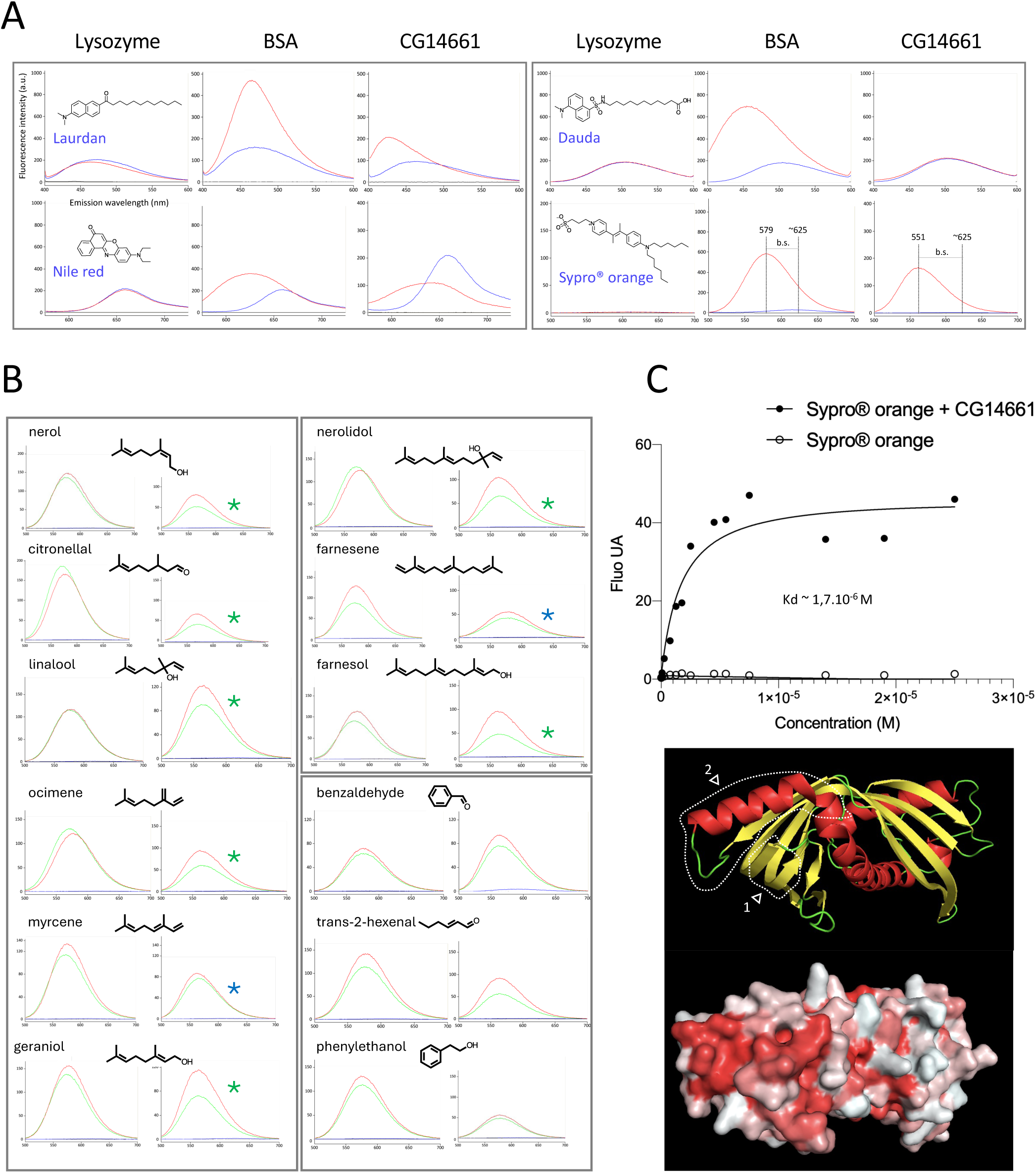
Fluorescent Competitive Binding Assays. **A:** Fluorescence emission spectra of four lipophilic fluorescent probes (1 µM) in the absence (blue line), or in the presence (red line) of 4 µM protein. Lysozyme is used as a negative control, as it does not have hydrophobic binding grooves (as BSA does), nor a hydrophobic pocket (like CG14661-6xHis). Note the great increase of fluorescence intensity of Sypro® orange in the presence of either BSA or CG14661-6xHis, and the shift of maximal emission towards shorter wavelengths (« blue shift », b.s.) that is typically seen when fluorescent lipophilic probes interact with hydrophobic domains of proteins (Pelosi et al., 2018b). Excitation wavelengths were 385 nm for Laurdan, 552 nm for Nile red, 335 nm for Dauda and 470 nm for Sypro® orange (Kroeger et al., 2017; Steinberg, 2009). **B:** A panel of monoterpenes (left), sesquiterpenes (upper right) and small volatile molecules (lower right) were tested for their ability to compete with the fluorescent lipidic probe Sypro® orange for binding to either BSA (left columns) or recombinant CG14661 (right columns). The emission spectra of the fluorescent probe in the presence of protein are shown as a red line, and the emission spectra in the presence of a 5x molar excess of competitor is shown as a green line. Note that except for myrcene and farnesene (blue asterisks), terpenoids proved better competitors for probe binding to CG14661 than to BSA, suggesting that CG14661 has a significant affinity for these compounds. **C: Top:** An estimate of the Kd of CG14661-6xHis for Sypro® orange was obtained in a saturation experiment using 4 µM protein and increasing concentrations of fluorescent probe. Note that a fivefold molar excess of ligands only partially competed with Sypro® orange (**B**), indicating that their Kds for CG14661 must be greater than 2 µM (see also text and Material and Methods). **Bottom:** Position of conserved Takeout motifs 1 and 2 (dashed lines), as described by (Hamiaux et al., 2009; Saurabh et al., 2018; So et al., 2000) based on Alphafold structure prediction (Jumper et al., 2021). These regions delineate a hydrophobic groove (rendered with red color in bottom image using the PyMol tool color_h (Eisenberg et al., 1984), that is thought to participate in a docking event, or/and to ligand exit from the internal hydrophobic cavity (Hamiaux et al., 2009)). This region is a potential alternate binding site for Sypro® orange and/or terpenoid ligands.

Reckoning that B-TDPs expressed in olfactory organs may be required for the perception of plant volatile terpenoids, we generated three Crispr-Cas9 deletions at the 82E locus, either removing CG14661 (Δ14661), CG14661 and CG2016 (Δ14661-Δ2016), or the three genes of the locus (Δ82E) (Figure 1, Suppl Figure 2). The olfactory performances of mutant and wild type flies were compared with a modified DART repulsion assay (Direct Airborn Repellent Test, (Kwon et al., 2010), Suppl Figure 4). The behavior of flies (males and females) towards a panel of linear mono- and sesquiterpenoids selected among the most commonly found in plant volatile blends and essential oils (Baldwin, 2010; Maffei et al., 2011), and three non-terpenic compounds, was recorded over a 40 minutes assay. The steepness of the responses (Slope, S) and their amplitude (Δ), near the source of odor (short range) or over the total size of the device (long range), were computed and are summarized in Figure 7 (original DART data are presented in Suppl Figure 9). Our premise was that if olfactory B-TDPs are required for the transport of terpenoid odorants, mutant flies should be anosmic, at least in some extent. However, as shown in Figure 7, mutant flies did not behave differently from wild type flies for some compounds (myrcene, farnesol), and surprisingly, showed even stronger responses (either steeper, or with a greater amplitude), for others. This increased sensitivity was not the consequence of an enhanced locomotory activity in the mutants, judging on the basis of recordings of climbing assays performed routinely after each DART experiment (Suppl Movie 10, (Ganetzky and Flanagan, 1978; Gargano et al., 2005)). Preliminary experiments using natural complex blends (essential oils) showed the same trend (Suppl Figure 10). The apparent increased sensitivity to farnesal and nerolidol in Δ14661-Δ2016 double mutants relatively to Δ14661 mutants may indicate a functional redundancy of the CG14661 and CG2016 proteins (Figure 7). Surprisingly, the enhanced response to linalool and geraniol of Δ14661 mutants is not seen in double (Δ14661-Δ2016) or triple (Δ82E) mutants, and, even more strikingly, triple Δ82E mutants behaved as wild type flies for most compounds tested (Figure 7). A plausible explanation for this contradiction lies in the well documented insecticidal and neurotoxic effects of terpenoids (Bakkali et al., 2008; Finetti et al., 2020; Isman, 2006). We suspect that above a certain threshold of exposure to volatile terpenoids, flies could experience reduced or even impaired locomotion, thereby artifactually mimicking a lack of olfactory perception. Indeed, Δ82E triple mutant flies appeared more sensitive than control CS flies and simple or double mutants, with a significantly greater number of fully anesthetized or dead flies after exposure to linalool or citronellal during the 40 min of the DART assay (Suppl Figure 11). Altogether, the data show that flies deficient for B-TDP genes at 82E are not impaired for the perception of terpenoids. Rather, they display more robust avoidance responses than control flies.

**Figure 7:**
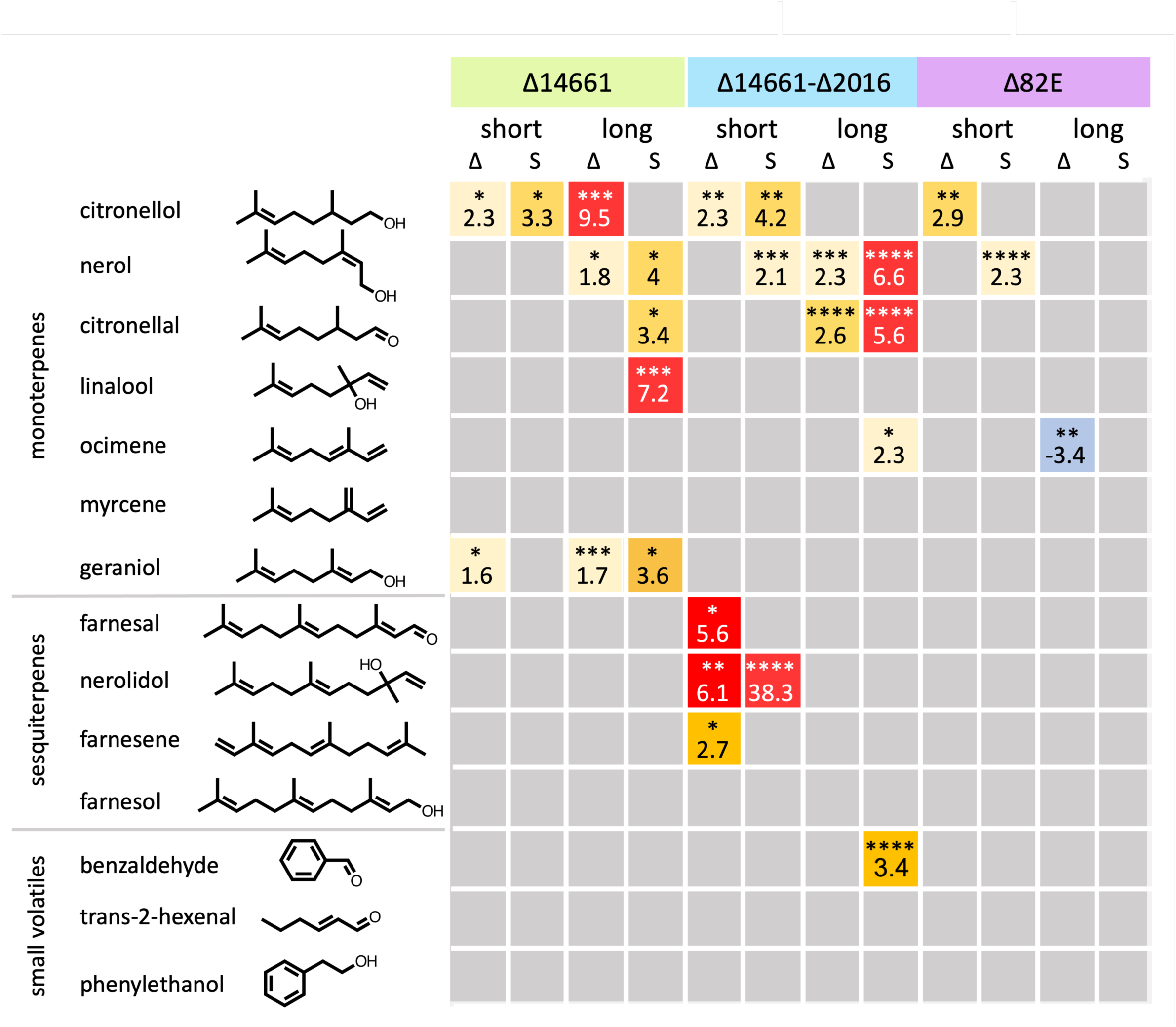
Functional (DART) assay of terpenoid perception in flies lacking B-TDP genes clustered at 82E. Repulsion response of flies lacking either CG14661 (Δ14661), both CG14661 and CG2016 (Δ2016) or all three TDM genes clustered at 82E (Δ82E), towards selected mono- and sesquiterpenes, or non-terpenoid volatiles was assessed with a modified <u>D</u>irect <u>A</u>irborne <u>R</u>epellent <u>T</u>est (DART, (Kwon et al., 2010), Suppl Figure 4). For each mutant line, the maximal distance from the odor source reached by the flies at the end of the experiment (Delta: Δ), and the steepness of the response at the beginning of the experiment (Slope: S) were determined. The number of flies was determined in different areas, to distinguish short-range from long-range effects (See Suppl Figure 4). Grey squares indicate non-significant differences relative to Canton S (CS) wild type control flies. Yellow, orange and red squares denote increased repulsion. Numbers indicate fold change relative to control flies. Note that most differences show an increased repulsion of mutants towards terpenoid volatiles when compared with the control line CS wild type flies. The formulas of odorants do not reflect isomery. All compounds were pure, except benzaldehyde (5%), trans-2-hexenal (2,5%) and ocimene (30%) (see Suppl Table 4 for details). Each experiment was replicated at least 11 times. Significant differences (one way ANOVA, followed by Dunnett’s multiple comparison test using the mean for CS wild type flies as the reference mean) are shown as * (P<0.05), ** (P<0.01), *** (P<0.0005) or **** (P<0.0001).

## 3. Discussion

### The *Drosophila melanogaster* full complement of B-TDPs

The TULIP superfamily of LTPs is recognized as very ancient, since it is present in a wide range of animal, fungal, algal, plant cells, as well as in prokaryotic cells (Alva and Lupas, 2016; Levine, 2019; Wong and Levine, 2017). Eukaryotic TDPs can be subdivided into intracellular SMP-like, and secreted BPI-like proteins (Alva and Lupas, 2016; Wong and Levine, 2017). Owing to an overall low level of identity at the amino-acid level, but a remarkable conservation of the TULIP domain ((Levine, 2019), Suppl Figures 1 and 7), findings of novel homologs have depended on the availability of sensitive informatic tools and 3D structures. Previous studies have therefore reported the presence of 21, 23, then 29 (Dauwalder et al., 2002; Li et al., 2016; Vanaphan et al., 2012), and more recently, only 16 B-TDP genes in *Drosophila* (Shin et al., 2022). Based on the latest release of the InterPro Ressource (InterPro 109.0, (Blum et al., 2025)), the full complement of *Drosophila* B-TDPs encompasses 30 members, grouped under the two fully overlapping protein domain signatures IPR038606 (Takeout superfamily) and IPR010562 (Haemolymph juvenile hormone binding) (Figure 1). The organization in seven distinct clusters (in addition to seven isolated genes), is typical of gene families, and thought to result from repeated gene duplications by unequal crossing-overs (Silver, 2001). Two genes, CG33680 and CG10264, showed no detectable expression at the RNA level (Figure 3). CG33680 is a pseudogene (Suppl Figure 6), but we found no obvious molecular defect in CG10264. ModENCODE Temporal Expression Profile data indicate that CG10264 is significantly expressed during larval and pupal stages (Gramates et al., 2022). The CG3246 gene is remarkable, as it is composed of two TULIP domains folded in a head-to-head orientation, and thus the only *Drosophila* representative of pseudodimeric relatives found in most other organisms (Suppl Figure 1, (Alva and Lupas, 2016; Levine, 2019; Wong and Levine, 2017)). Based on sequence alignments and on a structure-based phylogenetic analysis, its N-terminal and C-Terminal domains appear remotely related to those of the single TULIP domain proteins (Figure 1B). It is unclear if CG3246 is orthologous to the pseudodimeric B-TDPs of other organisms (such as BPI, CETP, LBP, PLTP, and L-PLUNCs / BPIFB in vertebrates, Suppl Fig 1, (Bingle et al., 2011; Krasity et al., 2011)), or if it results from an independent domain duplication event. It appears to have an essential function, however, since homozygous Crispr-Cas9 deletions of CG3246 were lethal in young first instar larvae (Figure 1A, Suppl Figure 2). In contrast, flies harboring homozygous Crispr-Cas9 deletions of either the 82E, 96C or 34D cluster, or combinations of homozygous cluster deletions removing up to eight genes (82E and 96C, 34D and 96C or 82E and 34D) were viable and displayed no obvious overt phenotype (Figure 1A, Suppl Figure 2).

### JHBPs are B-TDPs … but not all B-TDPs are JHBPs

The "Takeout" domain signature (Interpro domain IPR038606 (Blum et al., 2025)) refers to the first reported *Drosophila* member, *Takeout* (CG11853, (So et al., 2000)). The "JHBP" domain signature (Juvenile Hormone Binding Protein, IPR010562), is misleading, although it is related to the first and key finding that BPI-related proteins of vertebrates have homologues in insects (Kolodziejczyk et al., 2008). However, no *Drosophila* B-TDPs was ever shown to specifically bind Juvenile Hormone (JH).

In addition, phylogenetic analyses and structural considerations (absence of Takeout-motifs (Hamiaux et al., 2009; Saurabh et al., 2018)), Figure 6C) clearly show that JHBPs constitute a Lepidopteran-specific clade of B-TDPs, characterized by the presence of a conserved disulfide bridge and hydrogen bonds dividing the large internal hydrophobic cavity into two smaller ligand binding pockets (Suppl Figure 1, Figure 2B, (Hamiaux et al., 2013, 2009; Kolodziejczyk et al., 2008; Saito et al., 2006; Suzuki et al., 2011)). As previously shown by others, we confirm that no member of the present, updated set of *Drosophila* B-TDPs is orthologous to Lepidopteran JHBPs, while the existence of several orthologous groups of B-TDPs in *Drosophila* and the moth *Bombyx mori* (as a Lepidopteran representative) is readily evidenced (Figure 2A, (Hagai et al., 2007; Hamiaux et al., 2009; Li et al., 2016; Saito et al., 2006; Suzuki et al., 2011; Vanaphan et al., 2012)). The nature of *Drosophila* B-TDPs ligands thus remains uncertain.

### *Drosophila* B-TDPs are prevalent in chemosensory organs

The present study shows that about half of the *Drosophila* B-TDPs genes are expressed at higher levels in chemosensory organs in comparison with thorax and abdomen (Figure 3), confirming previous reports in Diptera and other insects that this protein superfamily may be involved in smell and taste (Bohbot and Vogt, 2005; Corcoran et al., 2015; Dauwalder et al., 2002; Fujikawa et al., 2006; Gu et al., 2011; Hamiaux et al., 2009; Justice et al., 2003; Levine, 2019; Rund et al., 2013; Saito et al., 2006; Sarov-Blat et al., 2000; Yoshizawa et al., 2011). The data shown in Figure 3 are ratios, and not absolute quantifications, hence some of these genes may also be significantly expressed in other organs or tissues, where they could contribute like their vertebrate counterparts to immunity against gram minus bacteria, lipoprotein metabolism and lipid trafficking, or to other yet unknown function (Alva and Lupas, 2016; C. D. Bingle et al., 2011; Krasity et al., 2011; Wong and Levine, 2017). Insect chemosensory transporters such as OBPs and CSPs were also shown to have additional functions, apparently unrelated to chemosensation (Pelosi et al., 2014; Rihani et al., 2021). In body samples (thorax and abdomen), all genes show a higher expression in males, thus confirming the previous observation that the expression of *Drosophila* B-TDPs is male-biased (Figure 3, (Vanaphan et al., 2012)).

The expression patterns of CG14661 and CG1124, in turn, are not associated with gustatory sensillae, but coincide with cells lining the pseudotracheae of the labellum (Figure 4). This pattern is puzzling, and reminds the localization of OBP19d, with is concentrated in the subcuticular space of the pseudotracheal region (Park et al., 2000; S. R. Shanbhag et al., 2001b). We hypothesized that these proteins may be secreted in the saliva to serve some function in feeding. However, we found no specific signal in "drink-blot" experiments (Liu et al., 2014), where the presence of CG14661-3xFLAG was examined with the M2 anti-Flag antibody (and a standard western blot procedure), on a piece of nitrocellulose wetted with a sucrose solution, on which starved transgenic CG14661 3xFL or wild type flies were allowed to feed for several hours (data not shown). The expression of CG1124 in the vicinity of the labral sense organ (LSO) and the ventral chemosensory sense organ (VCSO) suggests a gustatory function, although we were not able to ascertain its association with gustatory sensillae.

The expression of CG14661 and CG2016 in the sensillae of the third antennal segment and of the maxillary palp (Figure 5), the main peripheral olfactory organs of *Drosophila* adults, hints therefore to a putative function in odorant transport.

### B-TDPs qualify as a novel class of putative odorant transporters

Odorant transporters are expected to reside in the sensillar lymph, where they can encounter, solubilize, and shuttle lipophilic ligands to the dendrites of olfactory dendrites. We show here using confocal microscopy that CG1466 is likely secreted in the sensillar lymph, where it diffuses through the sensillar shaft, as was first demonstrated for its *Phormia regina* orthologue TOL in gustatory sensillae (Figure 5, Suppl Figure 1, (Fujikawa et al., 2006)). It can be reasonably hypothesized that other B-TDPs (such as CG2016, for example (Figure 5)), participate to the protein blend of the sensillar lymph, together with OBPs, CSPs and NPC2 proteins (Angeli et al., 1999; Hekmat-Scafe et al., 1997; Nagnan-Le Meillour, 2000; Park et al., 2000; Schmidt and Benton, 2020; S. R. Shanbhag et al., 2001b, 2001a; Zhu et al., 2018). Both CG14661 and CG2016 are only found in the trichogen cell, in contrast with previous evidence obtained for a few OBPs, where two or three types of accessory cells are involved (Park et al., 2000; S. R. Shanbhag et al., 2001b; Steinbrecht et al., 1992). A neuron expressing CG14661 was observed in a single among many preparations (Suppl Movie 7), inviting to scrutiny of expression data in future studies. Since it has been shown that OBPs can capture volatile ligands at an air-liquid interface and bring them into solution (Tsuchihara et al., 2005), the localization of CG14661 suggested that it could diffuse throughout the lymph cavity and be present within the *ca.* 30 nm pores of the sensillar shaft’s cuticle (Stocker, 1994), where it may capture ligands, directly at the lymph-atmosphere interface. However, despite trying several fixation and detection protocols, we failed to detect the CG14661-3xFLAG protein on the external surface of basiconic sensillae in scanning electron microscopy using anti-FLAG antibodies and secondary antibodies coupled to 5nm gold particles.

B-TDPs fulfill the four criteria expected for transporters of semiochemicals, as put forward by Pelosi and colleagues (Pelosi et al., 2014). First, B-TDPs exist as multiple gene families in all genomes surveyed so far, an anticipated condition to match the great variety of compounds of plant volatile blends (Malnic et al., 1999; Pelosi et al., 2014; Visser, 1986). The full complement of insect B-TDPs appears to be in the same range as OBPs (52 genes in Drosophila (Hekmat-Scafe et al., 2002)) and reported to include at least 8 genes in the honeybee (*Apis mellifera*, (Hagai et al., 2007)) and up to 43 genes in the diamondback moth (*Plutella xylostella*, (Li et al., 2016)). Next, B-TDPs fulfill the solubility criterion, and are relatively small (most insect B-TDPs are composed of a single, relatively small TULIP domain (*ca.* 25 kDa)), although somewhat larger than OBPs, CSPs and NPC2 proteins (*ca.* 14-20 kDa). Importantly, like other odorants transporters (Angeli et al., 1999; Hekmat-Scafe et al., 2002; Pelosi et al., 2014)), the structure of B-TDPs forms a compact pocket for lipophilic ligands ((Beamer et al., 1997; Hamiaux et al., 2013; Kolodziejczyk et al., 2008). Finally, an additional predicted characteristic of chemosensory transporters is that being in contact with the external environment, they should be chemically very stable. Small stable proteins often being powerful allergens, this consideration was used as a criterium to identify the Niemann-Pick disease related proteins as a new family of transporters for semiochemicals (Pelosi et al., 2014). Remarkably, this last criterion is fulfilled for at least one B-TDP, the Der P 7 protein, known as an important dust mite allergen (Alva and Lupas, 2016; Mueller et al., 2010).

### Volatile linear terpenoids as potential ligands for B-TDPs

Plant volatiles are essentially lipophilic and emitted as volatile blends dominated by terpenoids (Baldwin, 2010), which establishes *de facto* this large family of chemicals as a source of candidate ligands for chemosensory transporters. In vertebrates, most ligands studied so far are those carried by B-TDPs with two TULIP domains, also called pseudo-dimers (Alva and Lupas, 2016; Wong and Levine, 2017), Suppl Figure 1), and are rather bulky molecules, such as cholesteryl esters, phospholipids, triglycerides and lipopolysaccharide (LPS) (Qiu et al., 2007; Weiss, 2003; Wong et al., 2019). Juvenile hormones are the only high affinity (Kd*∼*25 nM), biologically relevant ligands ever characterized for B-TDPs in insects (Dupas et al., 2020; Goodman, 1990; Prestwich et al., 1996). These linear sesquiterpenoids fit in one half of the elongated cavity of the single TULIP domain of JHBPs ((Kolodziejczyk et al., 2008; Suzuki et al., 2011), Suppl Figure 1). JH binding B-TDPs (commonly referred to as JHBPs) are found only in Lepidoptera (Hamiaux et al., 2013; Saito et al., 2006). There is experimental evidence however, that B-TDPs may transport terpenoids other than JHs. A yellow B-TDP (YPT), responsible for the characteristic color of sexually mature gregarious males of the desert locust, *Schistocerca gregaria*, was purified and shown to bind a carotenoid (belonging to tetraterpenoids, (Wybrandt and Andersen, 2001)). Spectrophotometric data further showed that the YPT protein, and ALTO, another *Schistocerca gregaria* B-TDP, can bind lutein (also a tetraterpenoid, (Sugahara et al., 2020)). Ntsp1, an abundant B-TDP synthesized by cells lining the frontal gland of the termite *Nasutitermes takasagoensis*, is thought to act as a transporter for terpenoids, a major component of the defensive secretions in this species (Hojo et al., 2005). In addition, the hydrophobic cavity of Epiphyas posttvitana EpTO1 B-TDP, when expressed in *E Coli,* was crystallized with Coenzyme Q8, a surrogate bacterial ligand and a tetraterpenoid as well, that fully occupies the internal hydrophobic cavity (Hamiaux et al., 2009). Considering the relatively narrow, tubular shape of the hydrophobic pocket of B-TDP’s (Alva and Lupas, 2016; Hamiaux et al., 2013, 2009), linear compounds are more likely ligands than cyclic (and bulky) terpenoids. Since insects are guided in their chemical environment by host-specific blends of ubiquitous volatiles, rather than host-specific molecules (Bruce et al., 2005; Bruce and Pickett, 2011), we selected a panel of candidate ubiquitous linear volatile terpenoids for binding and behavioral assays (Figures 6 and 7). Our preliminary competition fluorescence assays establish the commonly used protein stain Sypro® Orange as a promising fluorescent tool for future studies of B-TDPs (Figure 6A). *E coli* expressed CG14661 significantly bound some linear terpenoids, although with apparent moderate affinities (Kd ∼ 3,8 µM for farnesol (Suppl Figures 3A and Figure 6B,C)). The apparent Kds measured when performing this type of assays, however, is very dependent of the fluorescent probe, so it therefore possible that the affinity of the linear terpenoids tested is higher than suggested by competition data of Figure 6B (Tan et al., 2020). Additionally, the single domain B-TDPs harbor a relatively large hydrophobic pocket (Hamiaux et al., 2013, 2009; Kolodziejczyk et al., 2008; Suzuki et al., 2011), able to accommodate a large linear ligand such as ubiquinone-8 (C49, tetraterpenoid (Hamiaux et al., 2009)), or two myristic acids (C14 (Hamiaux et al., 2013). It is thus conceivable that simultaneous binding of Sypro® Orange and terpenoid ligands occurs in distinct or overlapping areas of the pocket, thereby potentially reducing displacement of the fluorescent probe. Therefore, if B-TDP’s likely can bind a variety of linear terpenoids with low or moderate affinities, the possibility of the existence, among the many thousands of natural terpenoids, of high affinity, biologically relevant ligands cannot not be ruled out. It is worth noting that the propensity of B-TDPs for binding lipid ligands calls for caution. Indeed, several surrogate ligands have been encountered in structural studies ((Hamiaux et al., 2013, 2009)), and low affinity binding of radio-labelled juvenile hormone (JH) by jp29, a B-TDP of the moth *Manduca sexta*, turned out to be misleading clue to consider it as a nuclear receptor for JH ((Charles et al., 1996; Palli et al., 1994, 1990), Suppl Figure 1). Therefore, unequivocal demonstration and quantitative assessment of terpenoids binding by B-TDPs will require methods such as isothermal calorimetry (which is less limited by the low critical micellar concentration of these compounds (notably monoterpenes (Turina et al., 2006)), and structural studies are also warranted to confirm that ligand binding truly occurs in the inner pocket of CG14661, and not on a hydrophobic groove at its surface (Figure 6C).

### Hypothetical functions of B-TDPs as terpenoid transporters

Altogether, our preliminary binding data are consistent with the premise that B-TDPs may shuttle a variety of low affinity terpenoid odorants in the sensillar lymph, yet olfactory performance of flies deficient for B-TPBs of the 82E cluster towards a panel of ubiquitous terpenoids was not impaired (Figure 7). In the hypothesis that B-TDPs are required to transport odorants through the sensillar lymph to specific receptors located in the dendrites of olfactory neurons, unaltered perception in mutant flies could be explained by redundancy and compensation by other B-TBPs. Further studies will therefore require building an exhaustive expression map of B-TDPs in olfactory organs, such as the OBP-to-sensillum map established by Larter and co-workers (Larter et al., 2016). Interestingly, mutant flies appeared more sensitive than wild type flies (Figure 7). We were not able to critically test this hyperesthesia with rescue experiments, using heat shock driven GAL4 and an UAS-CG14661 construct and various heat-shock protocols, although overexpression of GAL4 and CG14661 was verified by RT-qPCR (data no shown). Abnormal processing and/or secretion of over-expressed CG14661, or alteration of behavior due to overexpression of Gal4 (Rezával et al., 2007) may explain failure of rescue of the hypersensitization observed.

These results are nevertheless intriguingly reminiscent of those recently obtained for Odorant Binding Proteins (OBPs). A widely accepted model for more than 30 years for the mode of action of OBPs is that they act as transporter for hydrophobic odorant molecules through the aqueous sensillar lymph (Brito et al., 2016; Larter et al., 2016; Sun et al., 2018), yet recent careful genetic analyses in *Drosophila* showed that olfactory neuron activity in flies deficient for the most abundant OBPs of identified olfactory sensillae is not abolished nor diminished when compared to wild type flies (Larter et al., 2016; Sun et al., 2018; Xiao et al., 2019). Furthermore, responses to odor pulses in mutant flies were found to be stronger for many odorants, suggesting that ligand binding by OBPs may support other functions than odorant delivery to the odorant receptors through the sensillar lymph (Larter et al., 2016; Sun et al., 2018; Xiao et al., 2019). Indeed, an old and persistent controversy in the field relates to the actual function(s) of transporters of lipophilic compounds in the sensillar lymph. The first OBP transporter ever discovered was hypothesized to have a function in ligand clearing after stimulation (Vogt and Riddiford, 1981). At the time, the prevalent hypothesis was that lipophilic odorants may reach their cognate receptors through a mono-dimensional adsorption pathway at the surface of nanometric pore tubules attached to the pore openings of basiconic sensillae on one side, and establishing several contacts with dendritic membranes on the other (Larter et al., 2016; Steinbrecht, 1997). Clearing of odorants is essential for insects to be immediately aware of fast fluctuations in the concentration of the stimulus, as when leaving a plume of pheromone, or when flying in a complex background noise in the odorant landscape, scanning for the ephemeral blend of molecules matching the host plant (Bruce and Pickett, 2011). A function in odorant clearing was indeed demonstrated for the OBPs OBP83b and OSF (OBP83a). In response to stimulation by a few odorants, including the sesquiterpenoid farnesol, the deactivation kinetics of olfactory neuron spiking was much slower in null mutants for both genes, indicating that these proteins normally work in dampening neuronal excitation (Scheuermann and Smith, 2019).

Co-evolution of plants and insects is thought to have started with herbivory in the Devonian period, 400 million years ago (Labandeira, 2013), bringing about the development of a plant chemical arsenal targeting herbivorous insects (Baldwin, 2010; Unsicker et al., 2009). The repellent and toxic (and in some cases neurotoxic) effects of plant terpenoids is well documented ((Bleeker et al., 2012; Finetti et al., 2020; Regnault-Roger et al., 2012; Stahl et al., 2018; Unsicker et al., 2009; Visser, 1986), and is the basis for the widespread use of essential oils as insect repellents or insecticides of (Bakkali et al., 2008; Isman, 2006; Lee, 2018; Tavares et al., 2018). Although the molecular and cellular underlying mechanisms for repellent or insecticidal effects of terpenoids are poorly known and may be very diverse, the chemosensory system, providing a direct route for volatile molecules towards sensillar cells, including neurons, is a likely target, perhaps through disruption of the normal function of the membranes of chemosensory neurons. In this hypothesis, B-TDPs and other transporter for lipophilic compounds may have been co-opted (True and Carroll, 2002) to provide a first line of defense, acting as low affinity scavengers for the great variety of harmful terpenoids or other plant volatiles encountered in the insect’s environment. The apparent hyperesthesia in flies deficient for B-TDP genes (Figure 7), and the increased spiking activity of olfactory neurons in chemosensory neurons of flies deficient for OBP28a (Larter et al., 2016), may therefore be caused by altered neuronal function, or stem from a pain (nociceptive) response, rather than result from failure of transport of a specific odorant to its cognate receptor. These observations encourage future studies to compare the resistance to the noxious effects of terpenoids in wild-type flies and flies deficient for B-TDPs and other chemosensory transporters.

Finally, in addition to the remarkably similar organization of the vertebrate and arthropod olfactory systems (Steinbrecht, 1998), human B-TDPs referred to as PLUNC proteins (Palate, Lung and Nasal epithelium Clone (Bingle et al., 2011; Bingle and Bingle, 2011)) are present in high concentrations in saliva, and the secretions of the upper airways, including the olfactory epithelium, where the B-TDP SPLUNC1 (BPIFA1) protein is one the most abundant protein (Genter et al., 2003), which suggests that B-TDPs may have ancient and conserved functions in chemical senses.

## 4. Material and methods

### Expression analyses

#### RT-QPCR

Standard procedures were used for RT-QPCR. RP49 was used as an internal standard and relative expressions were expressed with the ΔΔCt method (Livak and Schmittgen, 2001). The primers used are given in Suppl Table 1.

#### Crispr-Cas9

Gene deletion or edition were carried out with the Crispr-Cas9 by homology-directed repair (HDR) as described ((Gratz et al., 2015,), Suppl Figure 2), after evaluating gRNA target sites to minimize off-target cleavage (https://flycrispr.org/target-finder/). Cutting sites were chosen to avoid canonical nucleosomes positions likely to impede Cas9 activity (Horlbeck et al., 2016; Mavrich et al., 2008). Oligonucleotides corresponding to selected gRNAs were cloned into plasmid pU6-BbsI-chiRNA (Gratz et al., 2013), and replacement sequences were cloned into donor vector pHD-DsRed. donor vector. Plasmids were injected into Cas9 expressing early embryos y[1] M{RFP[3xP3.PB] GFP[E.3xP3]=vas-Cas9}ZH-2A w[1118]/FM7c (Bloomington stock# 51323). All deletions or editions were confirmed by PCR and sequencing.

### Production of recombinant protein and Competition Fluorescence Assay

The coding sequence of CG14661 and an additional stretch of 6 Histidine codons in place of the stop codon (CG14661-6xHis), was cloned downstream of the lac operator of the pET-24b vector. Three hours after induction with 0,5 mM IPTG, *E coli* cells were resuspended in a buffer (PBS 50 mM; imidazole 50 mM; PMSF 1mM; pH 7,4), and sonicated (3 x 20 seconds cycles at 65W). After centrifugation, the supernatant was loaded on an IMAC affinity column (HisTrap, Dutscher), and the recombinant protein eluted with an imidazole gradient on an AKTA pure FPLC system (GE Healthcare) was nearly pure (Suppl Figure 3A) and used in ligand binding assays after dialysis against appropriate buffers. Fluorescence analyses were performed on a Cary Eclipse Fluorescence Spectrometer (Agilent Technologies). Competition fluorescence assays were performed according to published guidelines (Pelosi et al., 2018a; Tan et al., 2020). The Kd of CG14661-6xHis for farnesol (∼3.8 µM) was estimated from the formula Kd_farnesol_ = [IC]50/(1 + [F]/Kd_Sypro®_), as in (Pelosi et al., 2018c), based on the estimation of the Kd of CG14661-6xHis for Sypro® (1,7 µM; Figure 6).

### Preparation of head chemosensory organs

For every sample, 100 to 500 heads of 5 to 7 days old flies were cut off and kept in 1,5 mL microtubes at -80°C until use. To harvest antennae and maxillary palps, microtubes were immersed in liquid nitrogen and vortexed for about 30 seconds. This procedure was repeated 3-4 times, after which antennae (mostly 2nd and 3rd segments) and maxillary palps detached from the heads were sieved through a fine nylon mesh into a clean microtube. These preparations contained mostly antennae and maxillary palps. Labellae (the most distal part of proboscis, were most gustatory sensillae are located), were hand dissected and frozen at -80°C until RNA extraction. For qPCR analysis, RNA samples extracted from these preparations were compared with "body" samples, consisting in thoraces and abdomens from which legs and wings were removed, to avoid including chemosensory organs located on these appendages.

#### Immunocytochemistry

Proboscis of individuals aged 5 to 7 days bearing maxillary palps were manually dissected and pierced with a fine needle in a solution of PBS 1X. Tissues were fixed in a 4% PFA/1X PBS solution for 45 minutes at room temperature with agitation, then rinsed for 5-10 min with 1XPBS at room temperature and transferred to blocking solution (0.5% Blocking Reagent (Roche), NaCl 0.15M, Tris HCl 0.1M pH 7.5) for 45 min at room temperature with agitation. They were then incubated with 1:500 goat anti-GFP (Rockland ref 600-101-215) for detection of GFP in MiMIC CG14661, CG2016, CG1124 gene trap lines (see paragraph below), or anti-flag M2 (F3165 Sigma-Aldrich, diluted 1:4000 in a solution of PBS 1X / 0.2% Saponin) for the detection of CG14661 3xFLAG, for 48 hours at 4°C. Proboscis were then rinsed (6 washes, 10 min each) with PBT and incubated with anti-goat alexa 488 1:400 and anti-mouse alexa 594 (ThermoFisher) diluted 1:400 in PBT in the dark under agitation overnight at 4°C. They were then rinsed (6 washes, 10 min each) with PBT and mounted between slide and coverslip in Vectashield mounting medium containing Dapi (Vector-H1200). Antennae were collected as described in the previous section, then fixed in PFA 4% / PBS for 2h at 4°C, washed in PBS, 3% Triton X-100 (2 x 15 min) and in PBS, 0.1% Triton X-100 for (3 x 20 min). They were transferred to blocking solution (0.5% Blocking Reagent (Roche), NaCl 0.15M, Tris HCl 0.1M pH 7.5) for 1-2h at RT. They were then incubated with anti-Flag M2 antibody (F3165 Sigma-Aldrich, diluted in TSA block at 1/4000 overnight at 4°C. After rinsing in PBS + 0.1% Triton (5 x 20 min at RT), they were incubated with anti-mouse alexa 488 diluted in PBT at 1/400 for 2-3h at RT in the dark. Finally, they were rinsed in PBS, 0.1% Triton X-100 (5 x 20 min) at RT, and mounted in Vectashield mounting medium containing Dapi (Vector- H1200). Fluorescence observations were conducted with a Leica SP8 Confocal Laser Scanning Microscope (Leica Microsystems, Wetzlar, Germany) using the 20x/0.75 objective (Numerical Aperture = 0.75) and the LAS X Microscope Software (v3.7.9; Leica Microsystems). Excitation wavelengths and emission filters were 552nm/band-pass 563–638nm for alexa594, 488nm/band- pass 500–584 nm for alexa488 and 405 nm/band-pass 415–482 nm for DAPI.

#### Fly stocks

Most stocks used in this study were obtained from the Bloomington Drosophila Stock Center (BDSC). Starting from the MiMIC line MI14535 *(Minos* mediated integration cassette, ((Venken et al., 2011); (BDSC # 59530)), we generated a protein trap line expressing the 20 first amino-acids of CG1124 fused to EGFP of the Double Header element, by recombination mediated cassette exchange (RMCE), using the dedicated BDSC fly stocks, and following the crossing scheme described in (Li-Kroeger et al., 2018). Cells expressing CG2016 were visualized in an RMCE line expressing GAL4 under the CG2016 regulatory sequences and carrying UAS-2xEYFP on the second chromosome (MI04995, (Bloomington stock # 77844)). The expression of CG14661 was documented using the MiMIC insertion line MI09778 (Bloomington stock # 55459). This line works as a gene trap, since it carries a MiMIC element inserted in the 5’UTRs of CG14661 in the orientation allowing expression of EGFP under the regulatory sequences of CG14661. Flybase (release FB2026_01) was the main resource for gene sequences and structures, bioinformatic tools and some expression data cited in the text (Öztürk-Çolak et al., 2024).

### DART assay

A modified Direct Airborne Repellent Test (DART, (Kwon et al., 2010)) was performed to quantify repulsion of flies towards terpenoids. In a typical assay, flies were exposed to vapors of pure molecules deposited on a piece of filter paper (1 µL), at one side of tube assembly as described in Suppl Figure 4. In the case of ocimene and essential oils, odorants were diluted in DMSO, which was loaded as a control (1µL) at the opposite side of the tube assembly.

### Climbing assay

Immediately after each DART repulsion assay, the locomotory performance of flies was evaluated with an assay taking advantage of the negative geotaxis reflex ("climbing assay", or RING assay (Ganetzky and Flanagan, 1978; Gargano et al., 2005)). Flies were tapped to the bottom of the tube, and a video of the flies walking back upwards was recorded. In some cases, a single picture was taken approximately 6 seconds after the flies were tapped down. Every experiment with 12 tubes typically included 6-11 replicates of a tested genotype, and 1-6 replicates with control (Canton S flies).

### Statistical analyzes

All analyses were performed with Graphpad Prism version 5.0d for Mac OS X (GraphPad Software, La Jolla, California USA).

## 6. Legends of Supplementary Figures, Tables and Movies.

**Supplementary Figure 1:**
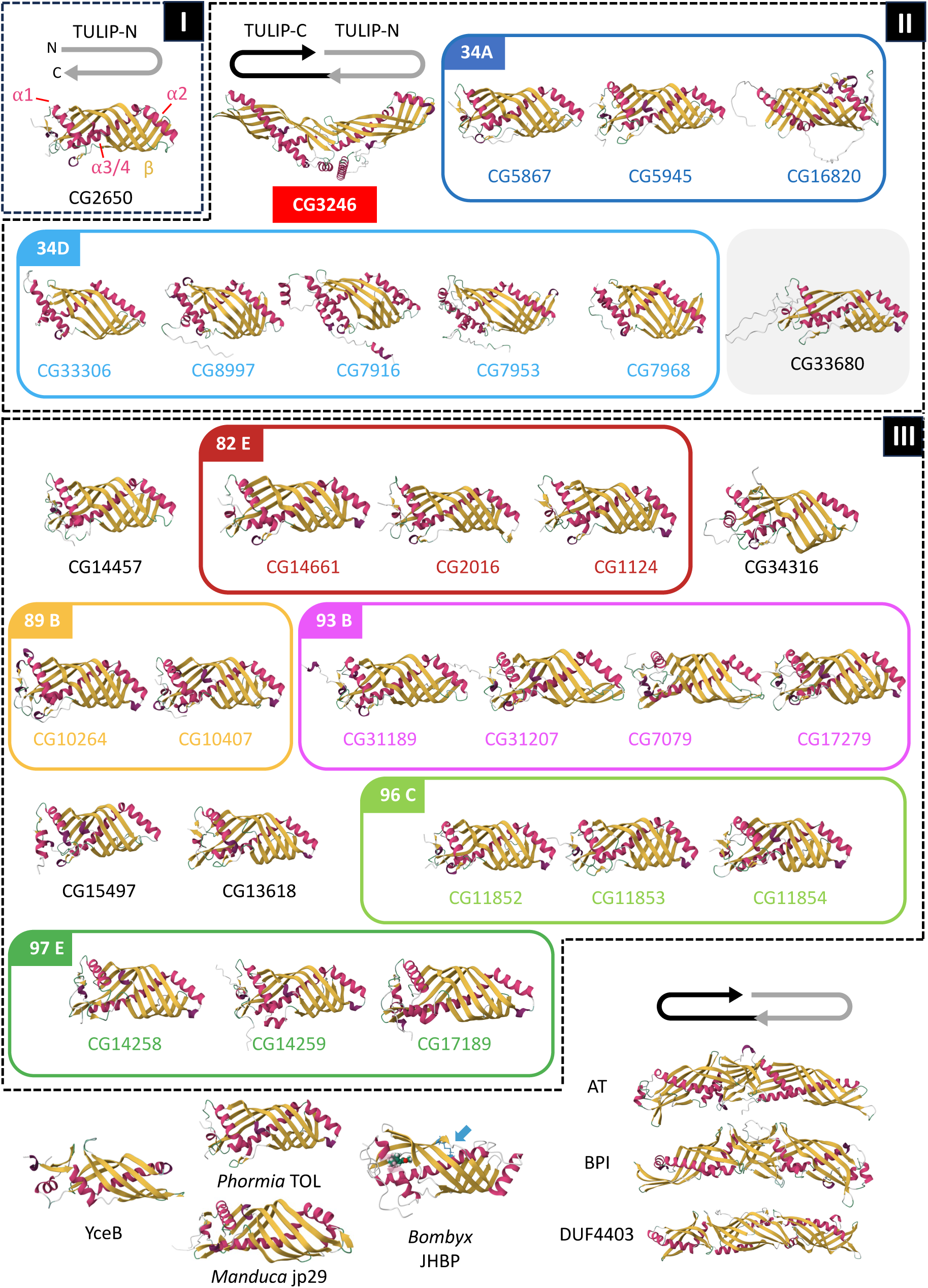
Structure predictions for BPI-like TDPs of *Drosophila* (Top), and other organisms (Bottom). **Top**: The Alphafold structure predictions (Jumper et al., 2021) of the 30 *Drosophila* BPI-like TDPs are shown in the orientation depicted by gray and black arrows. Dotted boxes represent chromosomes I-III, and colored boxes correspond to the seven genomic clusters (see text), with the same color code and cytological order as in Figure 1. The main common structural elements of the TULIP domain (α helices 1-3/4 and ß meander) are highlighted for CG2650. The protein on a grey background (CG33680) corresponds to a pseudogene (see text and Suppl Figure 6). Note that The CG3246 protein is the single pseudodimeric *Drosophila* BPI-like TULIP, composed of two TULIP domains folded in a head-to-head orientation, similarly to some related members found in other organisms (bottom right). **Bottom**: Representative BPI-like TDPs of Bacteria (YceB, DUF4403), insects (*Phormia* TOL, *Bombyx* JHBP, *Manduca* JP29), the plant *Arabidopsis thaliana* (AT: at1g04970) and the founding member, human Bactericidal, Permeability Increasing (BPI, (Beamer et al., 1997)). BPI and JHBP are X-ray structures (PDB 1EWF and 2RQF, respectively). JHBP is shown in complex with juvenile hormone III (space-filled), with an arrow pointing to a disulfide bridge (blue arrow) separating the hydrophobic pocket into two separate cavities (Kolodziejczyk et al., 2008; Suzuki et al., 2011).

**Supplementary Figure 2:**
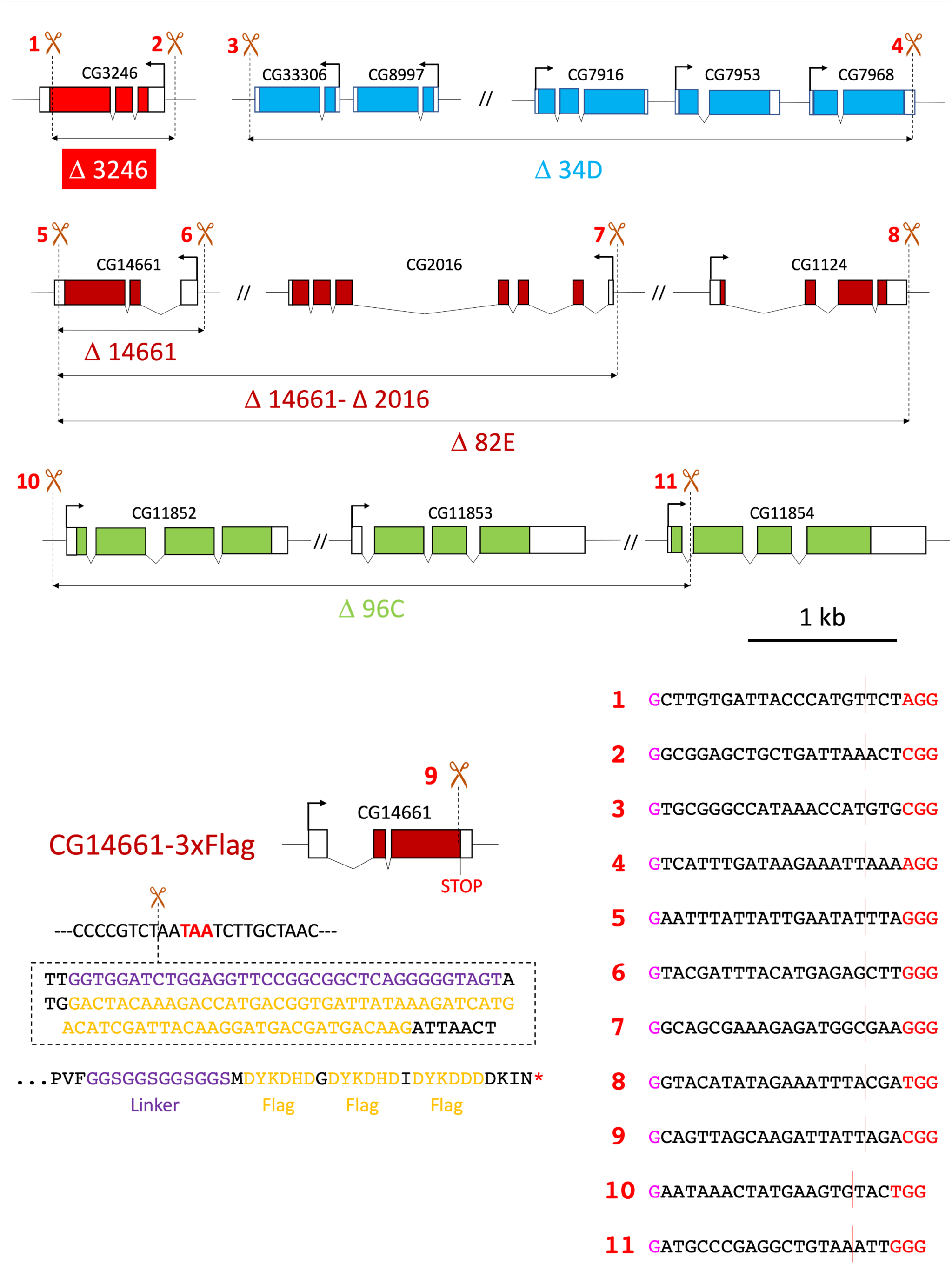
Deletions and editing of B-TDP genes and gene clusters. Approximate locations of Cas9 cleavage sites are indicated with dotted lines, and sequences of gRNA targets used are given at the bottom right. For each target sequence, the protospacer adjacent motif (PAM) is show in red, at the 3’ side. The 5’G in purple was added to improve transcription initiation from the U6 promoter of pU6-BbsI-chiRNA (Gratz et al., 2015). The extent of each deletion is shown as a with a line and arrowheads. For the CG14661-3xFLAG line, the location of the inserted sequence 5’ of the Stop codon of CG14661 in shown in dashed box. The resulting C-terminal amino acid sequence of the edited CG14661 protein is shown at bottom. The 3xFLAG epitope is recognized by the M2 antibody (F3165 Sigma-Aldrich).

**Supplementary Figure 3:**
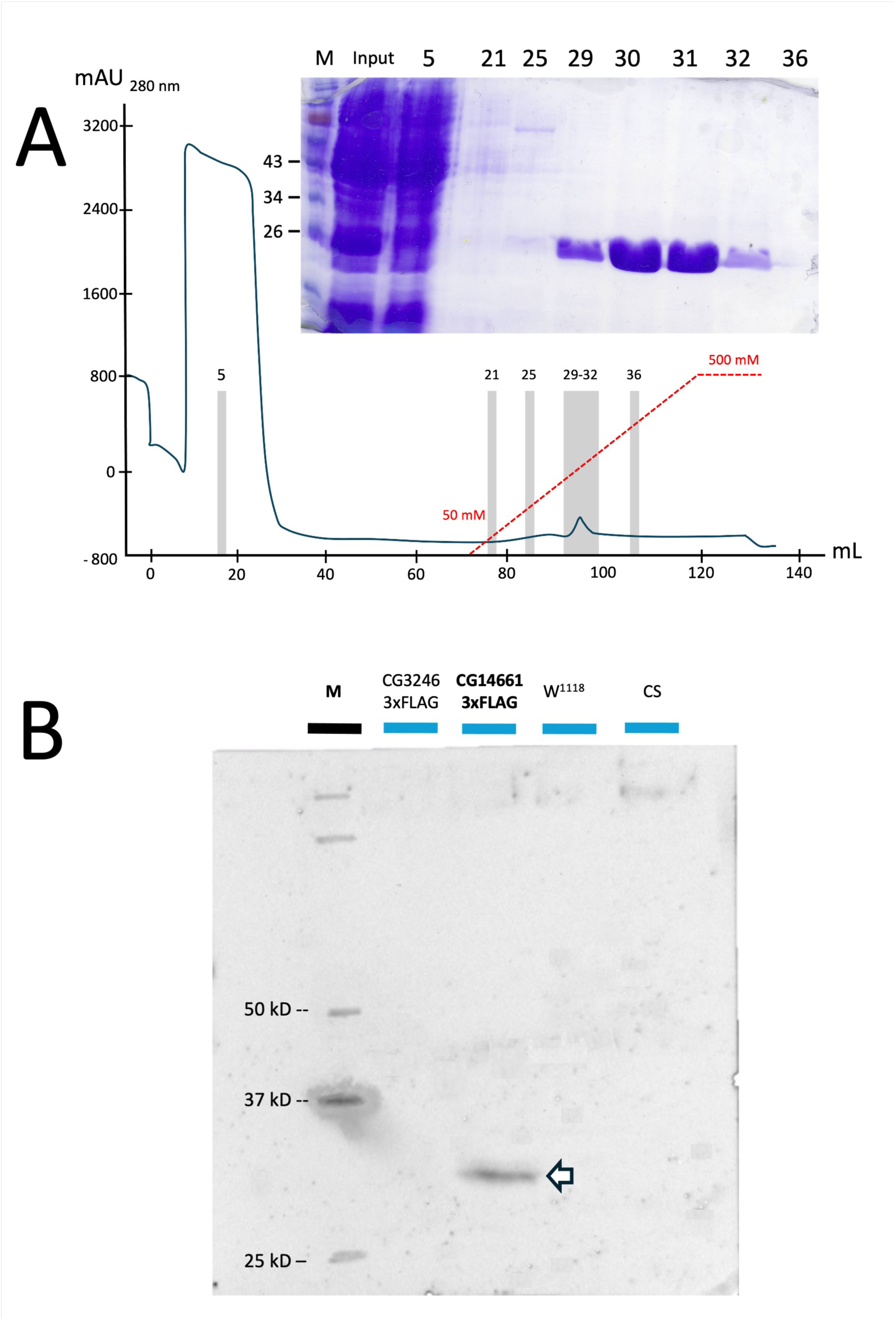
Expression and purification of CG14661-6xHis in E coli (A) and Western-blot of fly extracts with anti-FLAG antibody (B). **A:** The coding sequence of CG14661 (codon-optimized for *E coli*) and an additional stretch of 6 Histidine codons in place of the stop codon, was cloned downstream of the lac operator of the pET-24b vector. Three hours after induction with 0,5 mM IPTG, E coli cells were resuspended in a buffer (PBS 50 mM; imidazole 50 mM; PMSF 1mM; pH 7,4), and sonicated (3 x 20 seconds cycles at 65W, on ice). After centrifugation, the supernatant was loaded on an IMAC affinity column (HisTrap, Dutscher), and the recombinant protein eluted with an imidazole gradient on an AKTA FPLC system (GE Healthcare) was nearly pure and used in ligand binding assays after dialysis. **B:** Western blot analysis recombinant of wild type (CS, W^1118^) and transgenic (CG3246-3xFLAG, CG14661-3xFLAG) fly head extracts (10 heads per sample), with an anti-FLAG antibody (anti-Flag M2 antibody (F3165 Sigma-Aldrich)). The arrow points to a band with a size consistent with the CG14661-FLAG protein (30,5 kDa).

**Supplementary Figure 4:**
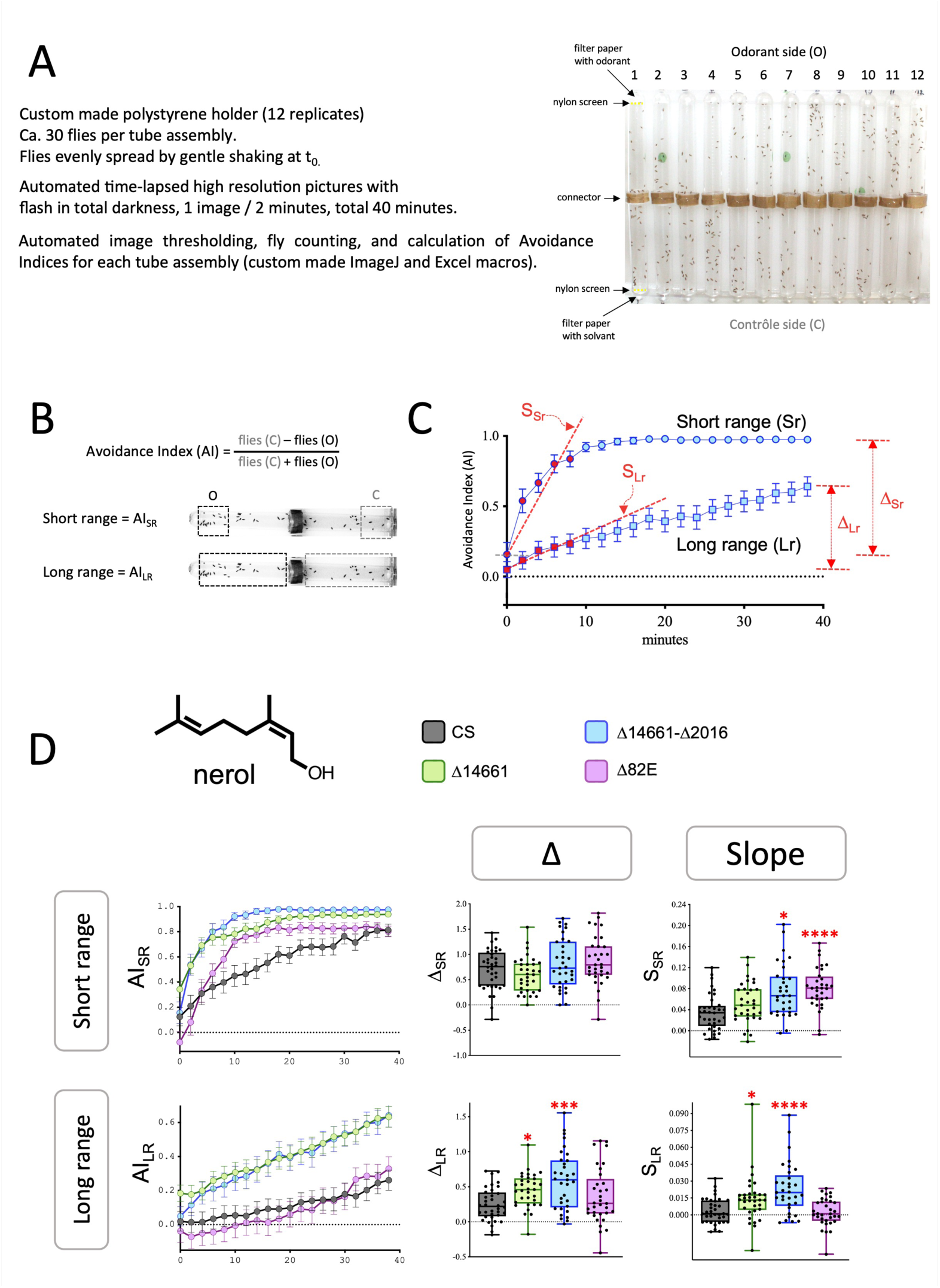
Olfactory assay. A DART assay (<u>D</u>irect <u>A</u>irborne <u>R</u>epellent <u>T</u>est) modified from (Kwon et al., 2010), was performed to assess the ability of adult flies to perceive repulsive odorants (**A**). Odorants (1 µL of pure substances or dilutions in DMSO) were loaded on a piece of filter paper placed on the odorant side of two tubes assembled with a connector, and 1 µL of DMSO on the other side when required. All compounds were used undiluted, except benzaldehyde (5%), trans-2-hexenal (2,5%) and ocimene (30%). Access to the odorant / solvent to the flies was prevented with a nylon screen. About 30 flies were loaded in the tube assembly, and 20 high resolutions pictures were automatically taken over the course of the 40 minutes experiment, in complete darkness. Flies were automatically counted in two small areas near the tube ends to calculate an Avoidance Index (AI) reflecting the behavior of the flies in the vicinity of the odorant source ((**B**), Short range, AI_SR_). A second AI was determined by counting flies in two large areas, to reflect flies’ movements at a larger distance from the odor source ((**B**), AI (Long Range, AI_LR_)). The changes of AI_SR_ and AI_LR_ were graphed over time (**C**), allowing to record the steepness of the response of the flies (increase of AI upon exposure to odorant) over the first 8 minutes of the experiment (red data points, dotted red lines, average slope: S_SR_, S_LR_). The difference of AI between the beginning and the end of the experiment was also recorded as Δ (Δ_SR_ and Δ_LR_). (**D**): Example of data obtained with nerol. Each point represents a tube with ca.∼30 flies. In Figure 7, only the fold change relative to the control wild type strain Canton S (CS) is reported along with the significance value, but the original data is gathered in Suppl Fig 9.

**Supplementary Figure 5:**
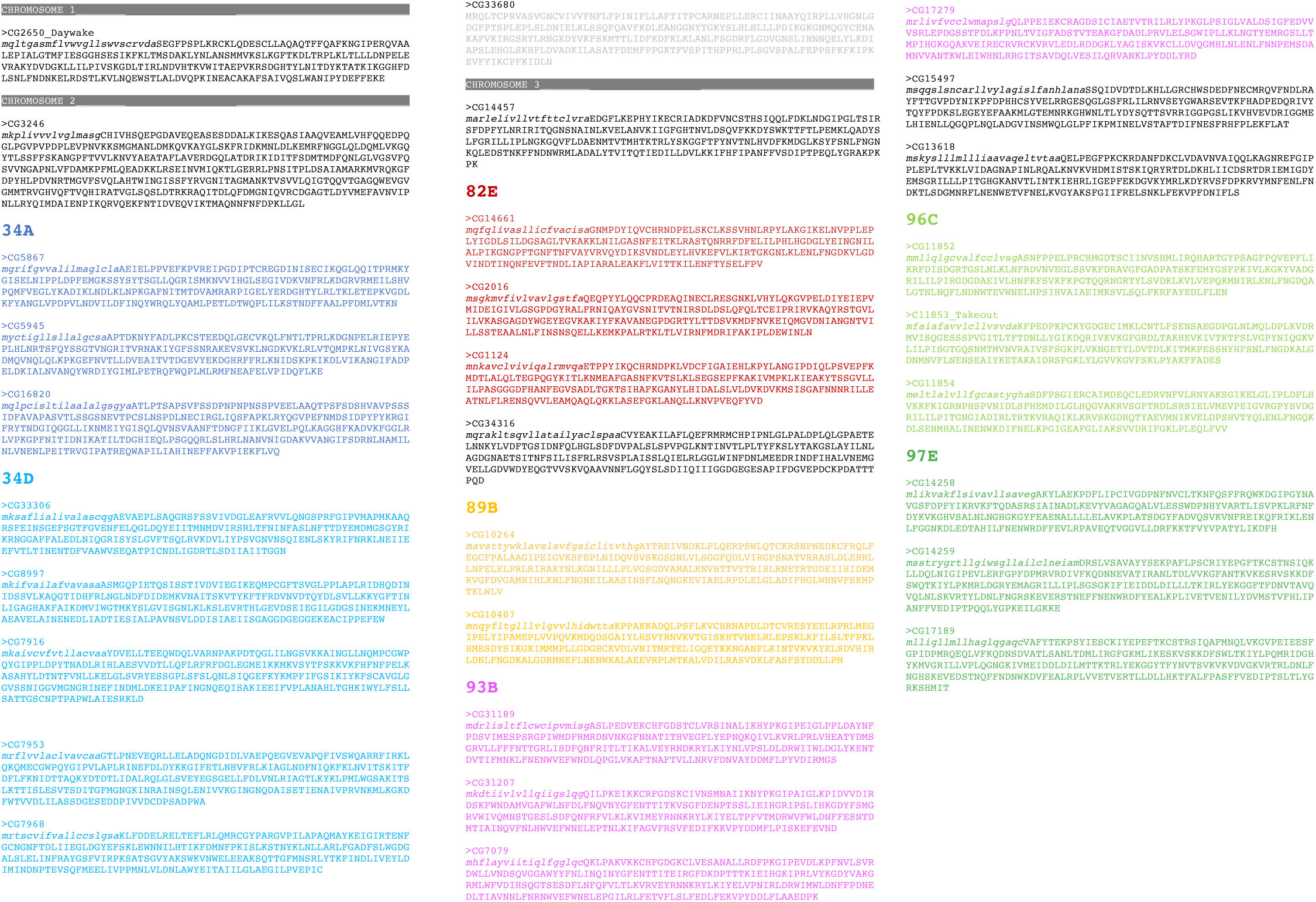
Sequences of *Drosophila melanogaster* B-TDPs. Signal peptides (standard Sec/SP1 secretory signals, lowercase italics) were predicted with SignalP 6.0 (Teufel et al., 2022), except for CG14259 (SignalP 5.0, (Almagro Armenteros et al., 2019b) and TargetP-2.0 (Almagro Armenteros et al., 2019a)). The sequence in pale grey (CG33680) correspond to a pseudogene (see text and Suppl Figure 6). Chromosomes I-III are separated by a grey line, colors correspond to the seven genomic clusters (see text), with the same color code and cytological order as in Figure 1.

**Supplementary Figure 6.**
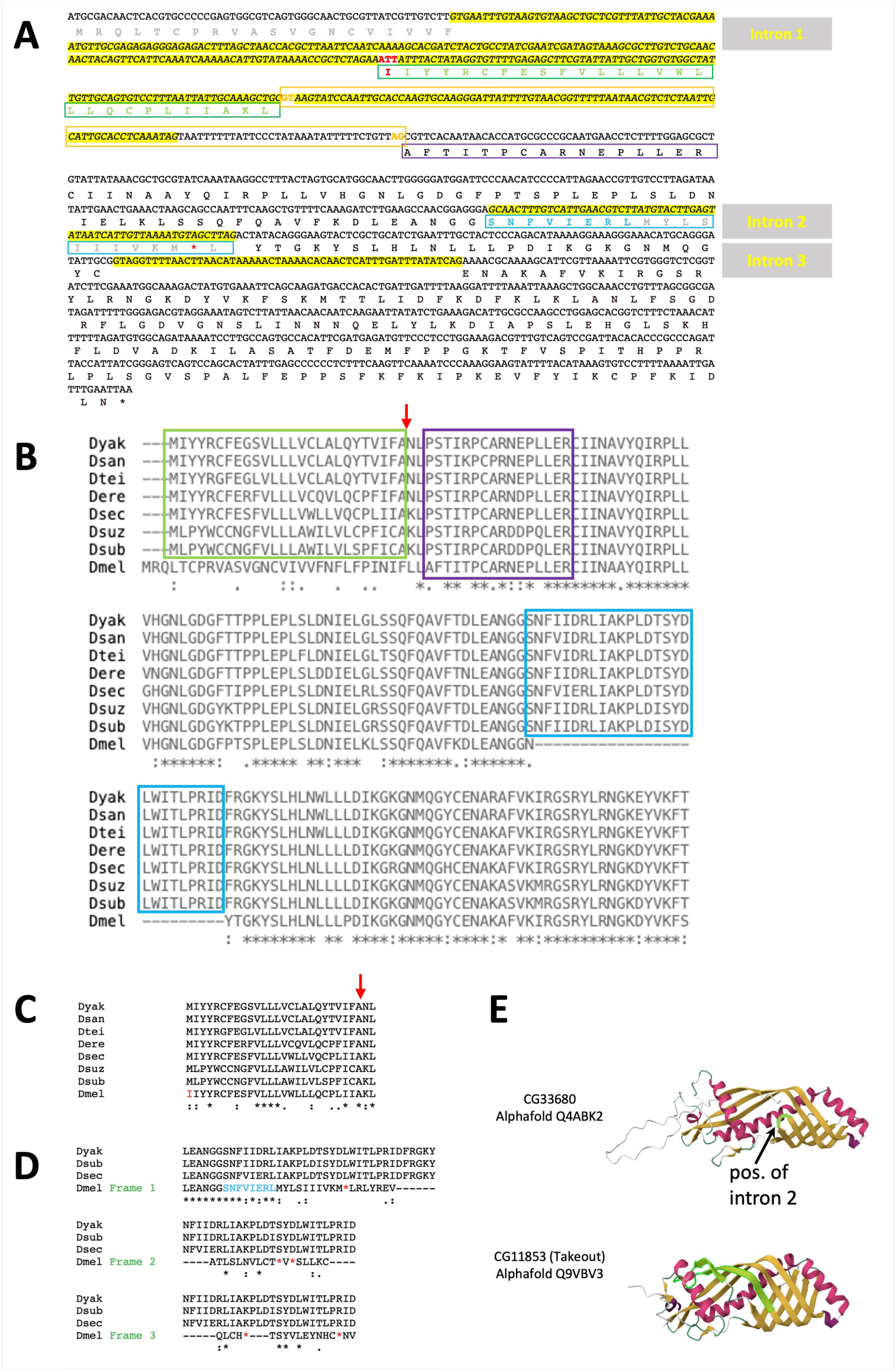
CG33680 is a pseudogene. **A:** Predicted transcript for CG33680 (NM_001032277.2 / FBtr0091660). **B:** Alignment of the predicted CG33680 protein and putative *Drosophila* orthologs with ClustalW (Thompson et al., 1994). **C:** Alignment of the putative signal peptide of D melanogaster CG33680 protein with the homologous region in putative *Drosophila* orthologs. **D:** Alignment of peptides obtained by translations of intron 2 and corresponding regions of putative *Drosophila* orthologs. **E:** Comparison of the structures of CG33680 and Takeout (CG11853) proteins predicted by Alphafold (Jumper et al., 2021). **A:** The N terminal sequence of the predicted CG33680 protein in the present gene model (MRQLT…PINIFLLAF, shown in grey letters in (**A**)) does not contain a standard secretory (Sec/SPI) signal peptide, based on SIgnalP-6.0 (Teufel et al., 2022), SignalP-5.0 nor TargetP-2.0 (Almagro Armenteros et al., 2019; Almagro Armenteros et al., 2019). This would make CG33680 the only *Drosophila melanogaster* TULIP domain protein with no secretion signal peptide, among the 30 proteins *D. melanogaster* B-TDPs. Running Flybase Blastx with the first 400bp of the CG33680 gene region (FBgn0053680, ATGCGACAACTCACG… TAACGTCTCTAATTG) as a query against GenBank protein sequences yields a *D. erecta* protein (XP_026835908.1, protein takeout), harboring a different N-terminal sequence (MIYYRCFE…LPS…), that constitutes a highly likely Sec/SPI signal peptide according to SignalP-6.0, SignalP-5.0 and TargetP-2.0 (likelihood = 0,9991 with SignalP-6.0, with a predicted cut between residues 26 and 27 (red arrow in (**B**) and (**C**)). Using this *D. erecta* protein as a query, NCBI BlastP yields many hits in the *Drosophila* genus, often fully aligning with the *D. erecta* protein, including the putative signal peptide, and likely corresponding to true orthologs (see alignment of seven among the best hits with CG33680, after removal of 26 first amino acids in (**B**)). Entries in (B) are XP_039228012 (*D. yakuba*), XP_032571828 (*D. sechellia*), XP_026835908 (*D. erecta*), XP_016927869 (D. suzukii), XP_039481884 (*D. santomea*), XP_037715276 (*D. subpulchrella*). Remarkably, a very similar peptide is encoded in the first intron of the current gene model (green boxes in (**A**) and (**B**), see also the alignment in (**C**)), and could be translated in frame with the conserved sequences located further downstream (purple boxes in (**A**) and (**B**)) after excision of an intron (orange box, 5’ and 3’ splice sites in orange letters) predicted by the Genie program (Reese et al., 1997). However, the initiator Methionine in *D. melanogaster* is replaced by an isoleucine, owing to a g->t transversion (codon highlighted in red in (**A**)). We re-sequenced this area in our Canton S stock and confirmed the presence of an ATT codon at this position. Intron 2 in the current model has a non-canonical donor GC site (GA-<u>GC</u>AACT) that is a poor match to the *Drosophila* consensus (AG-<u>GC</u>AAGT, (Kitamura–Abe et al., 2004). Translating this putative intron (Blue boxes in (**A**) and (**B**)) yields a peptide (SNFVIERL, blue letters in (**A and** (**D**)) that is conserved in CG33680 putative orthologs, followed by non-conserved residues (grey letters), and a STOP codon (red asterisk). The two other peptides potentially encoded in this putative intronic region (blue box in (**A**)) show little similarity to CG33680 orthologs and are also interrupted by STOP codons ((**D**), Frames 2 and 3). In addition, intron 2 region is too short to encode the whole peptide found in CG33680 orthologs, suggesting that *D melanogaster* CG33680 presents a deletion of about 17 bp. This questionable intronic sequence is however expected to encode important amino acids for the structure of the protein as shown in (**E**). The arrow in (**E,** top) points to the region (highlighted in green), where the peptide encoded by intron 3 should reside. By comparison, the corresponding region of the Takeout protein is shown in green (bottom).

**Supplementary Figure 7:**
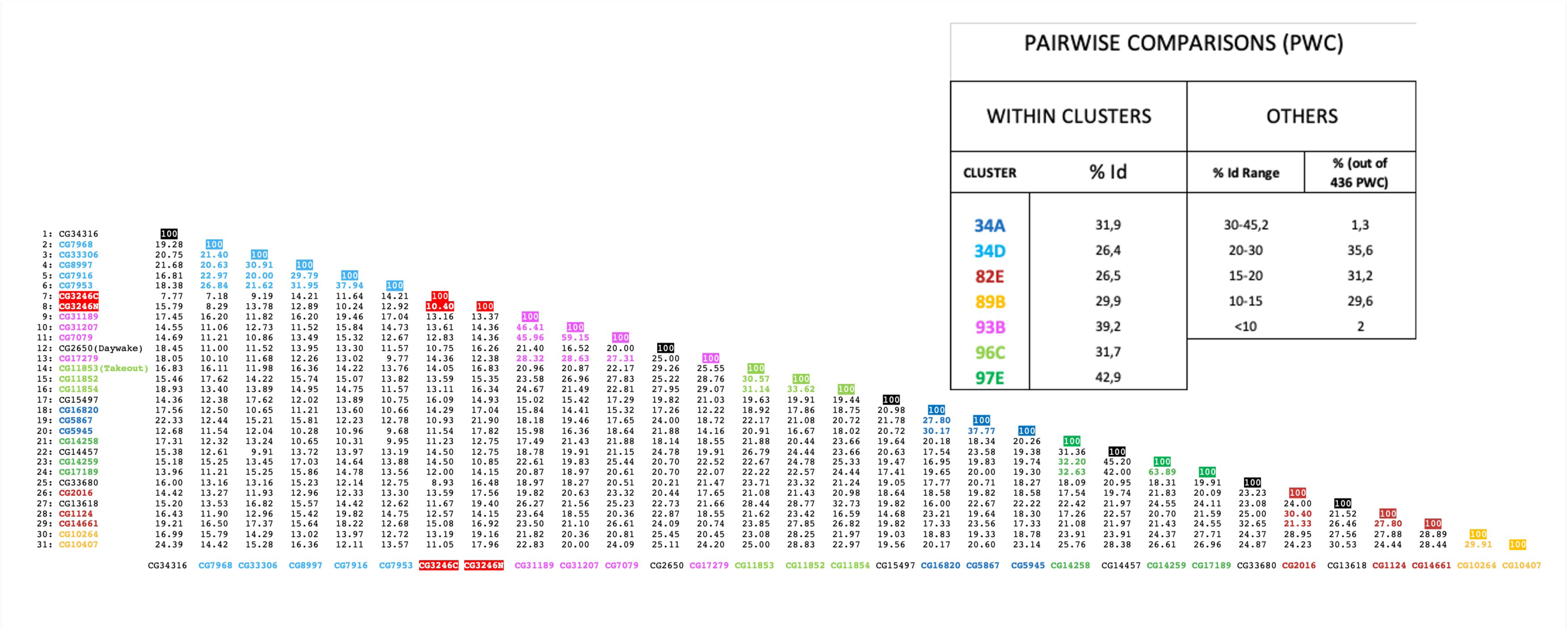
Pairwise comparisons (PWC) of *Drosophila melanogaster* B-TDP proteins. The thirty B-TDP proteins without their signal peptide were aligned with Clustal omega (Madeira et al., 2024) and the resulting table of pairwise comparison is shown, with numbers corresponding to % residues identity (% Id), and the same color code for clusters as in Figure 1. The table in inset shows average % Id values of PWC within clusters (left), and the distribution of % Id values of all (436) remaining PWC (excluding intra-cluster comparisons, right).

**Supplementary Figure 8:**
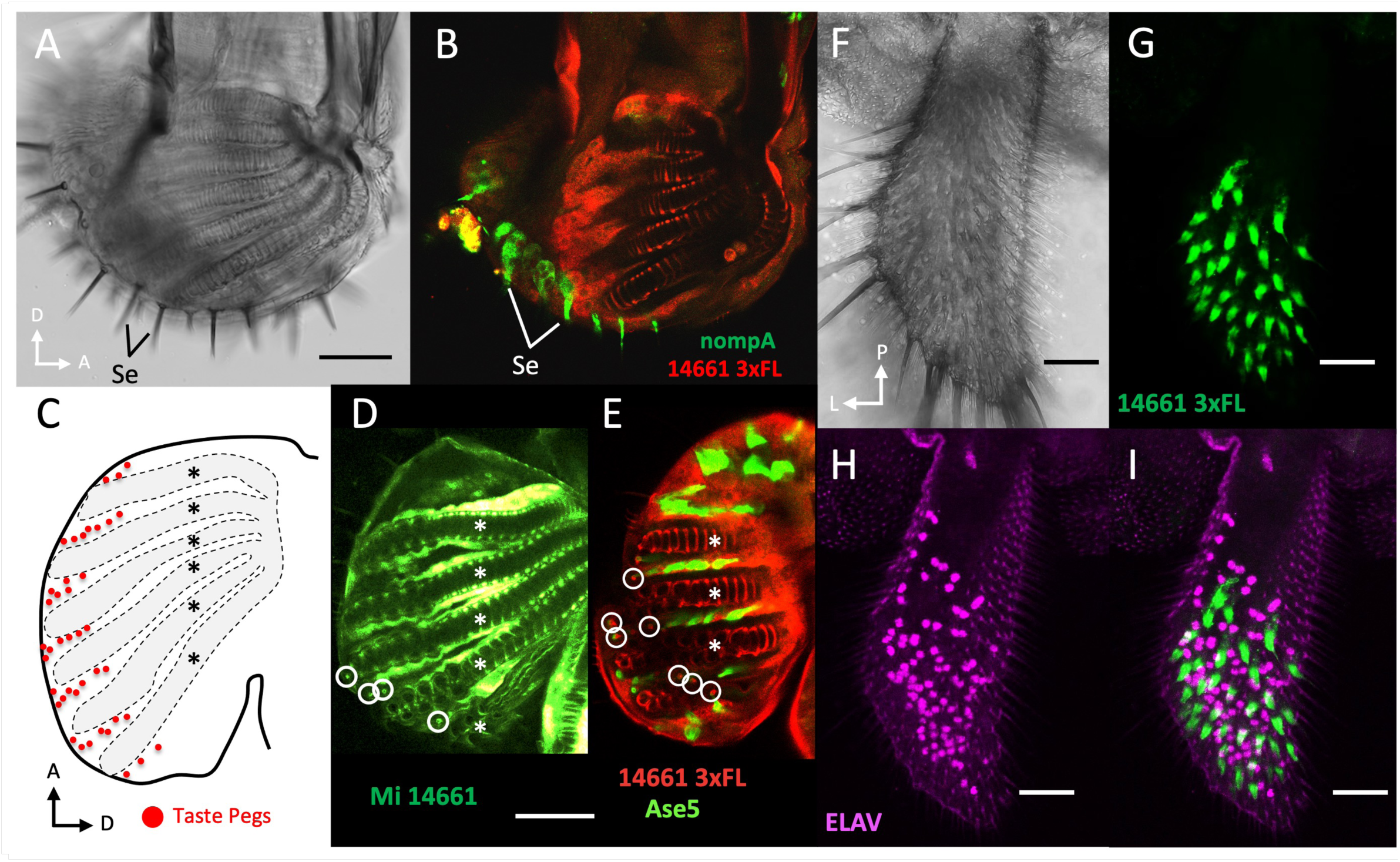
**A-B**: The expression domain of CG14661 does not overlap with gustatory sensillae on the surface of the labellum (as seen with the nompA marker, specific of the thecogen cell of the sensillae (Chung et al., 2001)). **C-E**: CG14661 is also expressed in small spots (white circles) located beetween the pseudotracheae (asterisks), that may correspond to small gustatory sensillae on the inner surface of the labellum referred to as taste pegs (red dots in **C**, redrawn from (S. R. Shanbhag et al., 2001b)). **F-I**: CG14661 is not expressed in neurons of the maxillary palp. The expression of CG14661 is analyzed with an anti-FLAG antibody in the 14661 3xFL fly line edited by Crispr-Cas9 to add a 3xFLAG epitope at the C-terminus of CG14661 (**B,E,G**) or with an anti-GFP antibody in the gene trap line Mi 14661 (**D**) (which reveals the cells transcribing the CG14661 gene). The neurons are vizualized with an antibody against the neuronal marker elav (**H,I**). **F**: Bright field. **G**: Expression of CG14661 3xFL in the same maxillary palp. **H**: Neuron nuclei. **I**: Merge of **G** and **H**. Note that the two patterns are not correlated. Thin black or white arrows indicate orientation (**A-E**: A: anterior, D: dorsal; **F-I**: L: lateral, P: proximal). Scale bars: A-E = 50 µM, F-I = 25µM.

**Supplementary Figure 9:**
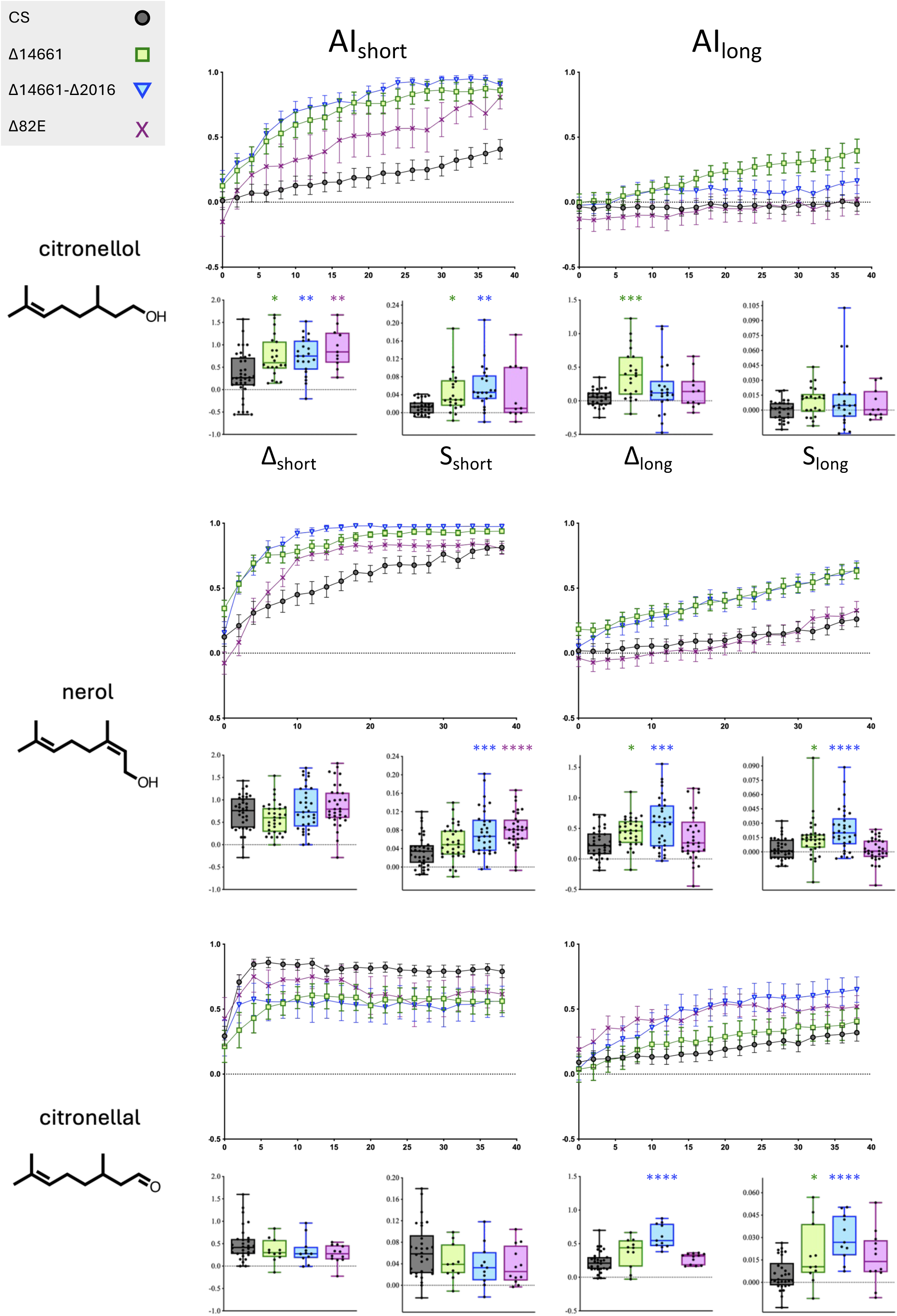
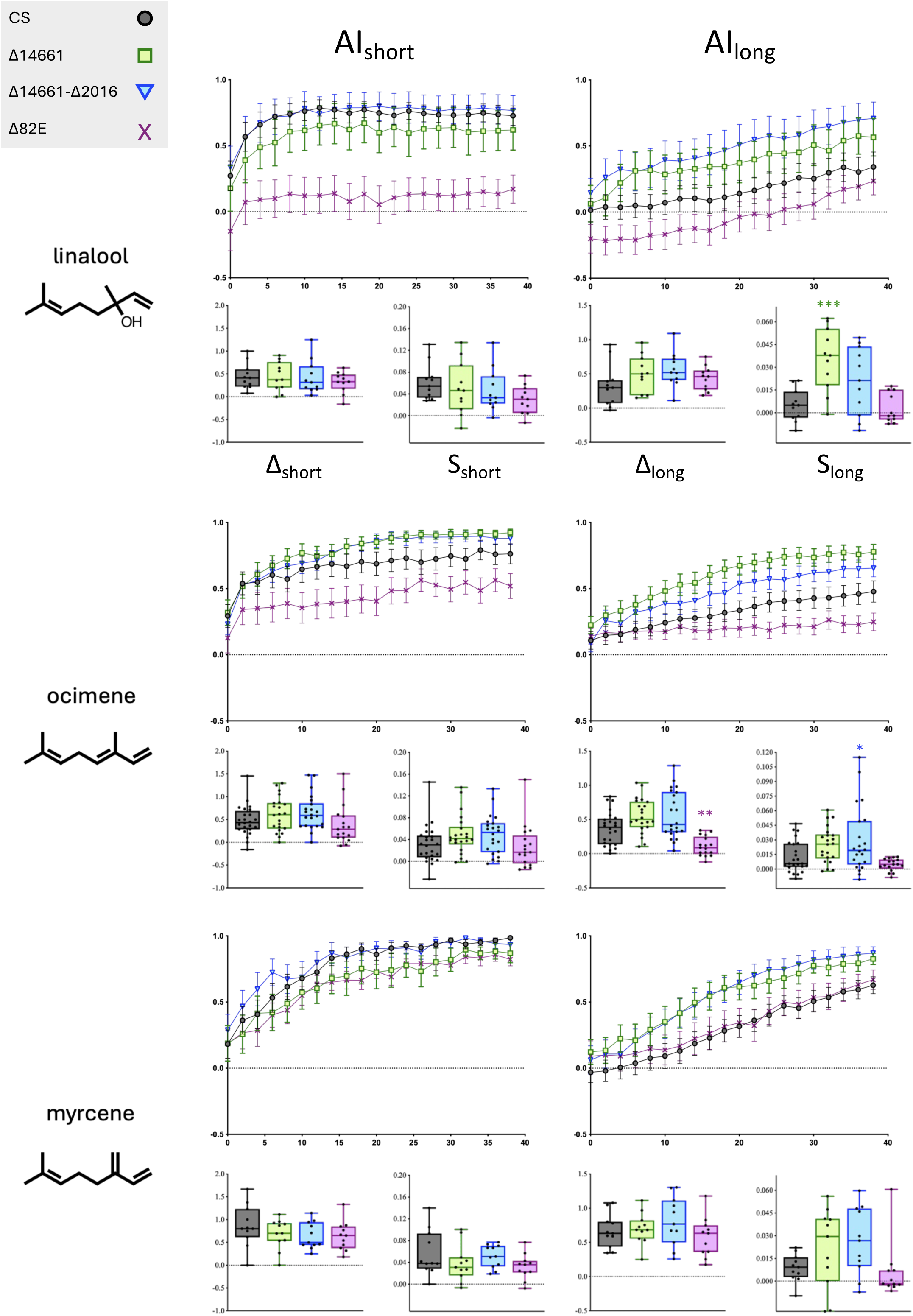
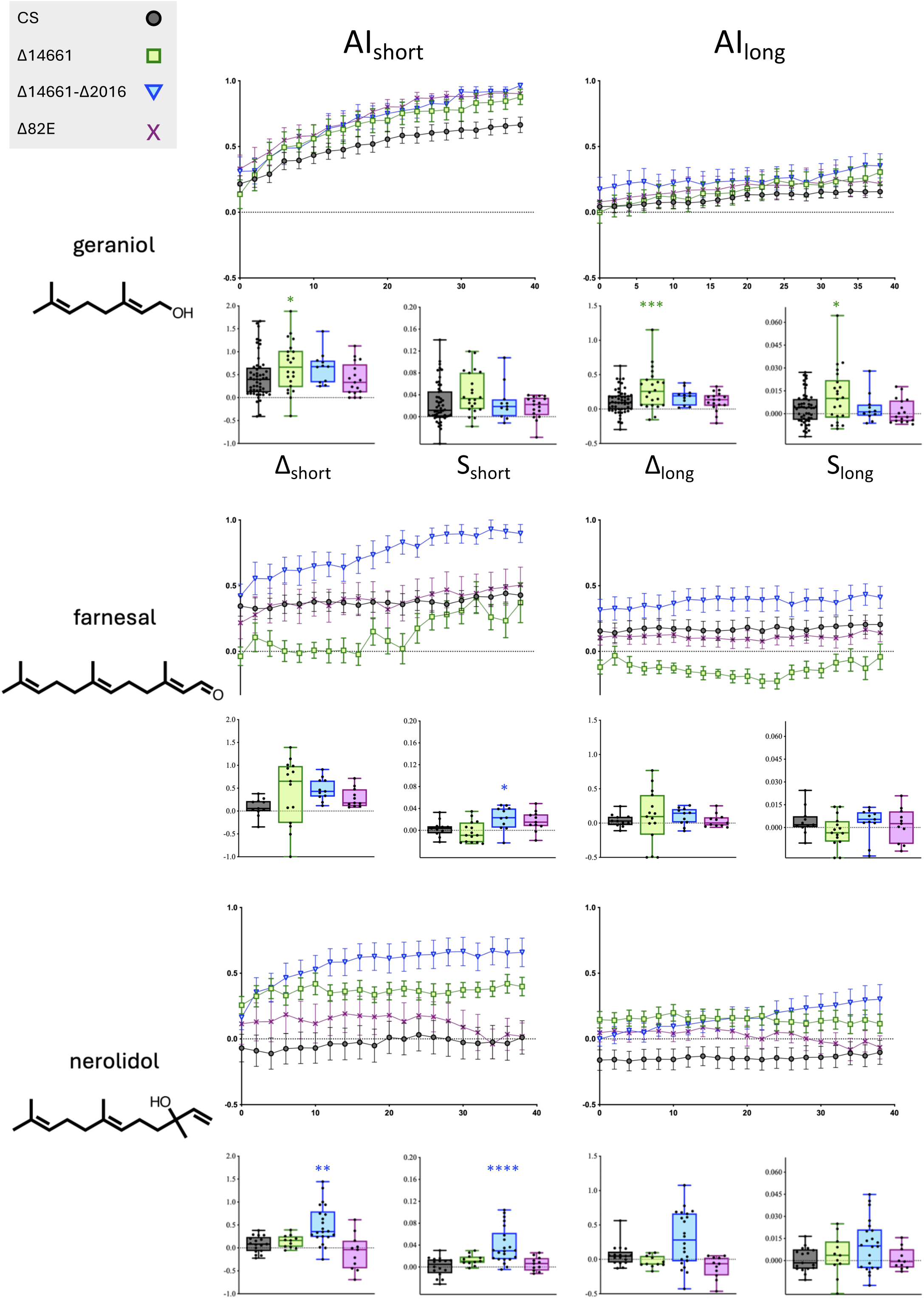
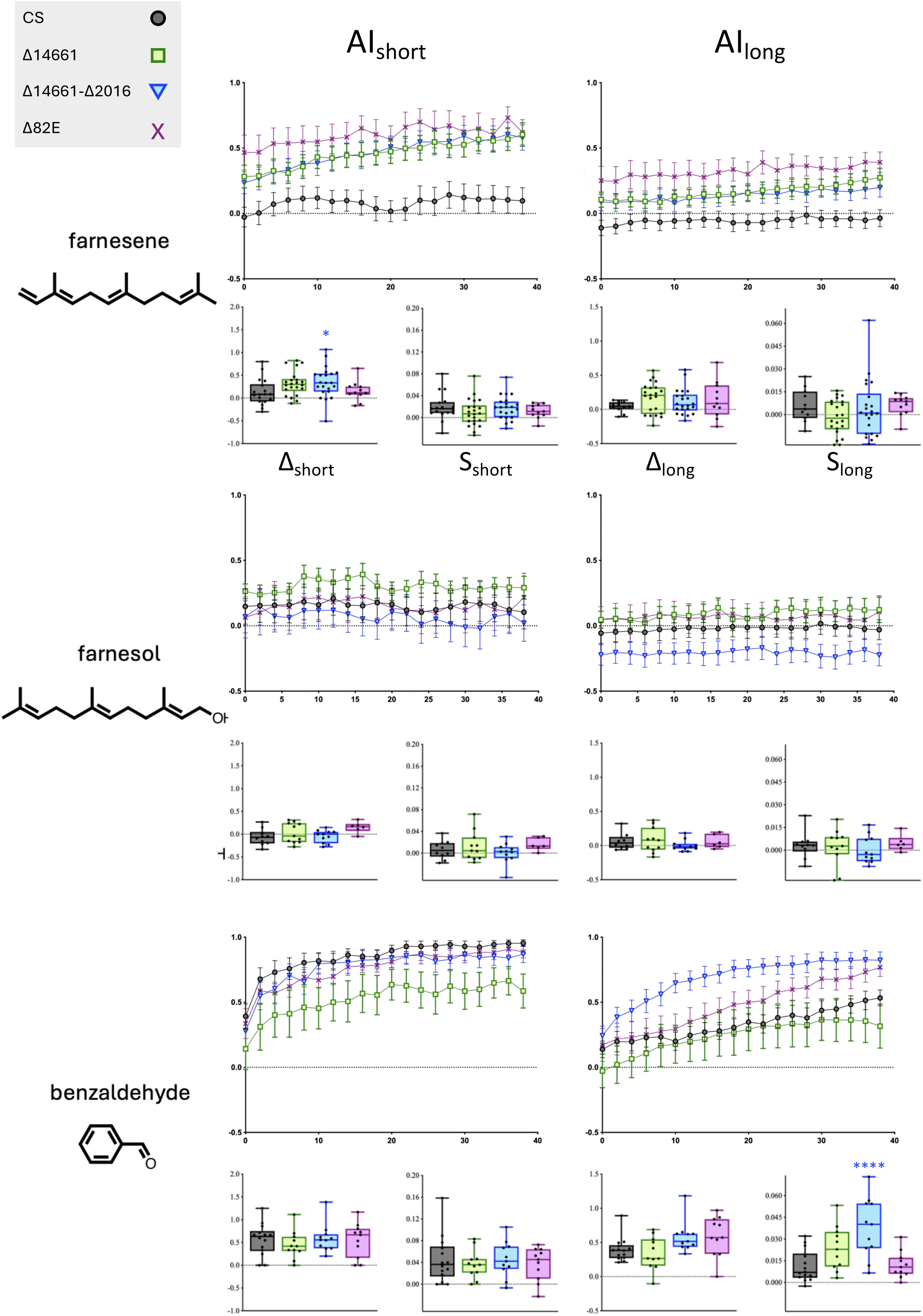
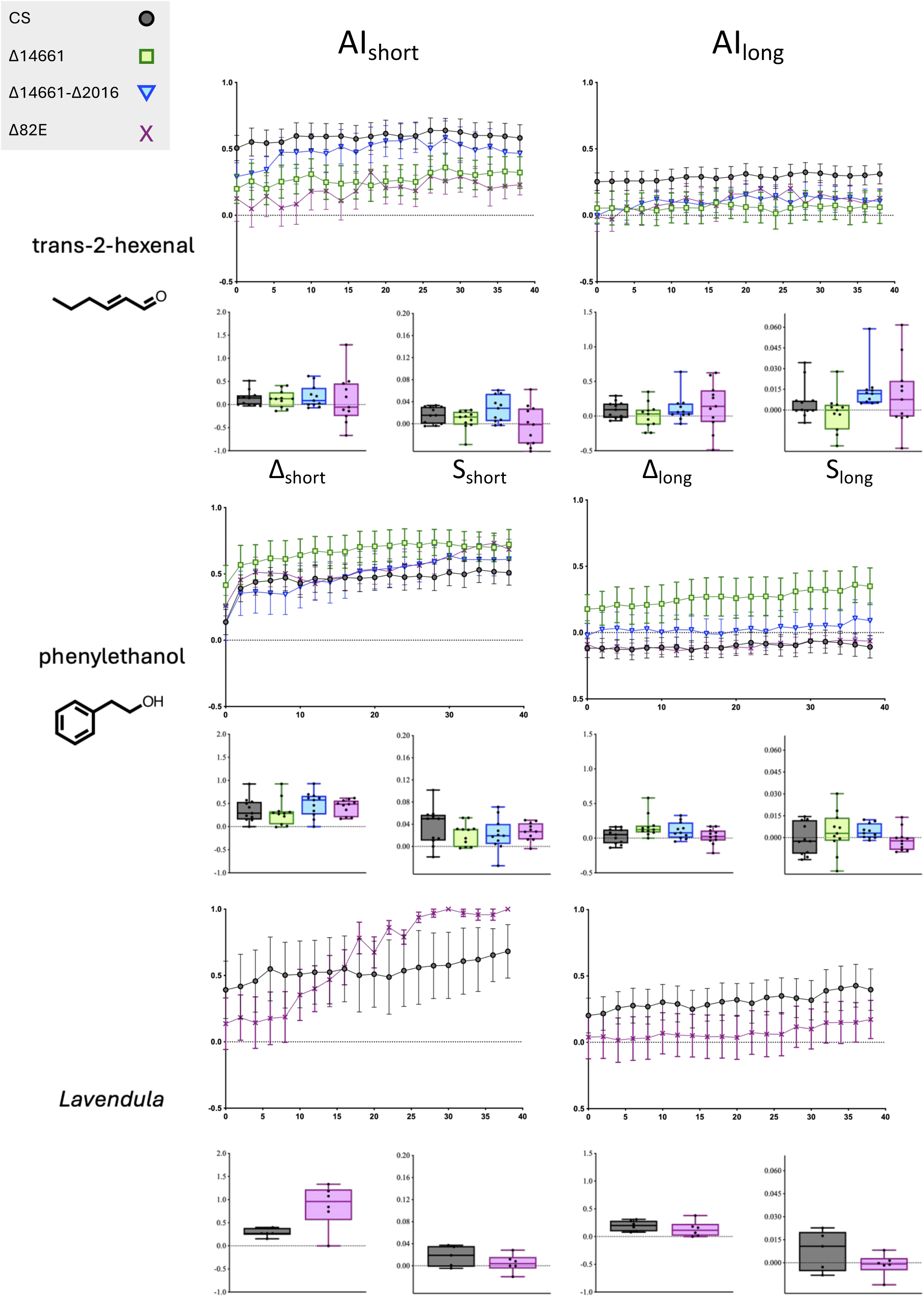
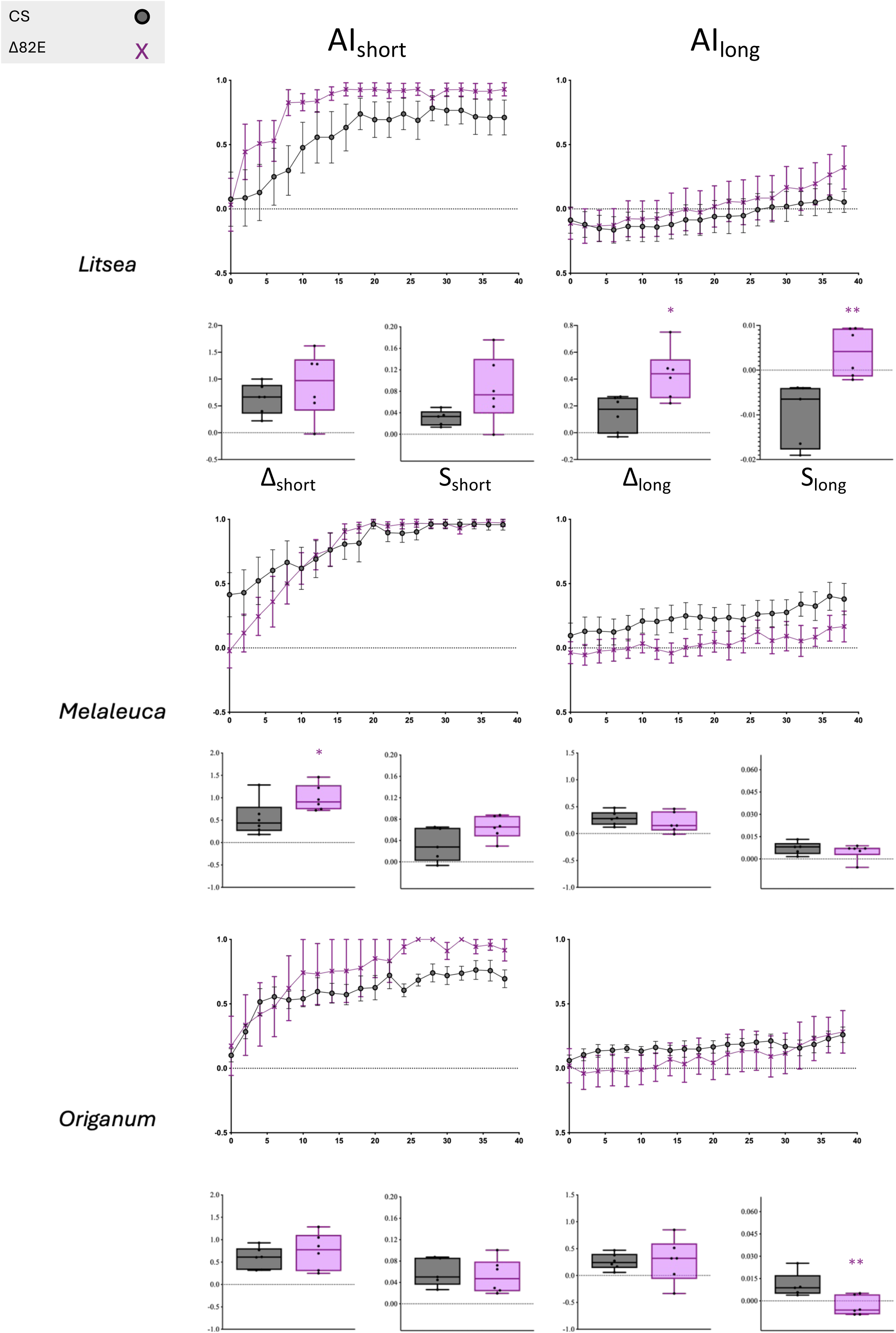
DART assay (<u>D</u>irect <u>A</u>irborne <u>R</u>epellent <u>T</u>est) data corresponding to Figure 7 and Suppl Figure 10. For each compound tested, top panels show the time course (in minutes on horizontal axis) of Avoidance Index at “short range” and “long range” (see Suppl Figure 4A,B). Errors bars represent the standard error of the mean (SEM), and the symbol correspond to the mean. The bottom panel report the span of the response (Δ: AI value at the end of the experiment, minus AI value at the onset of the experiment), and the slope of the time course change in AI value during the 8 first minutes of the experiment (S, Suppl Figure 4). Whiskers represent the span of data, shown as points. The rectangle extends from the 25th to 75th percentiles, and the horizontal line corresponds to the median. Means were compared as explained in Figure 7 (* (P<0.05), ** (P<0.01), *** (P<0.0005) or **** (P<0.0001)).

**Supplementary Figure 10:**
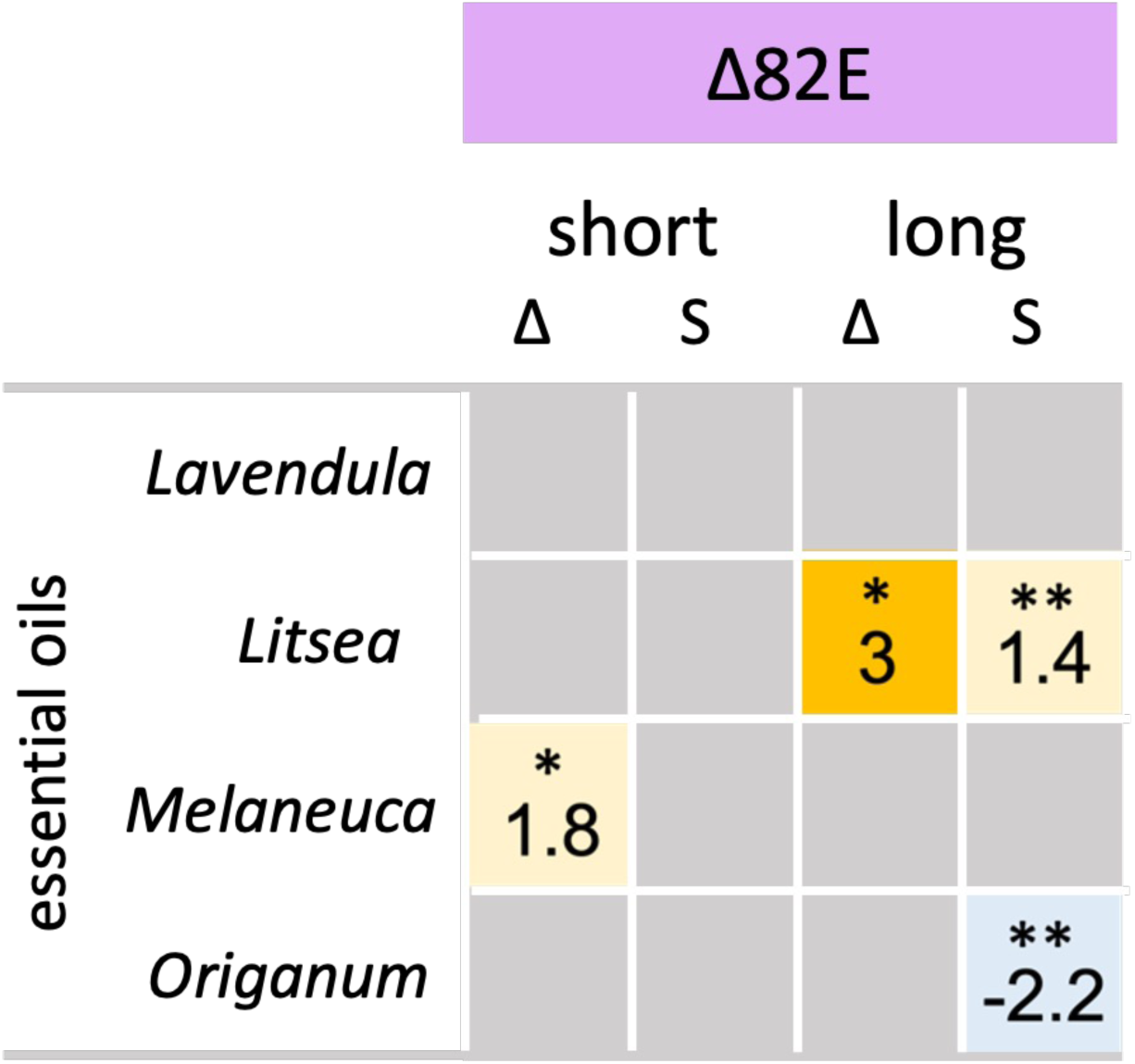
Repulsion response of flies lacking all three B-TDP genes clustered at 82E (Δ82E), towards terpene-rich plants extracts (essential oils). Conditions and parameters are described in the captions of Figure 7 and Suppl Figure 4. Original data are reported in Suppl Figure 9. The dilutions in DMSO were 10% (*Lavendula*, *Melaleuca*), and 5% (*Litsea*, *Origanum*).

**Supplementary Figure 11:**
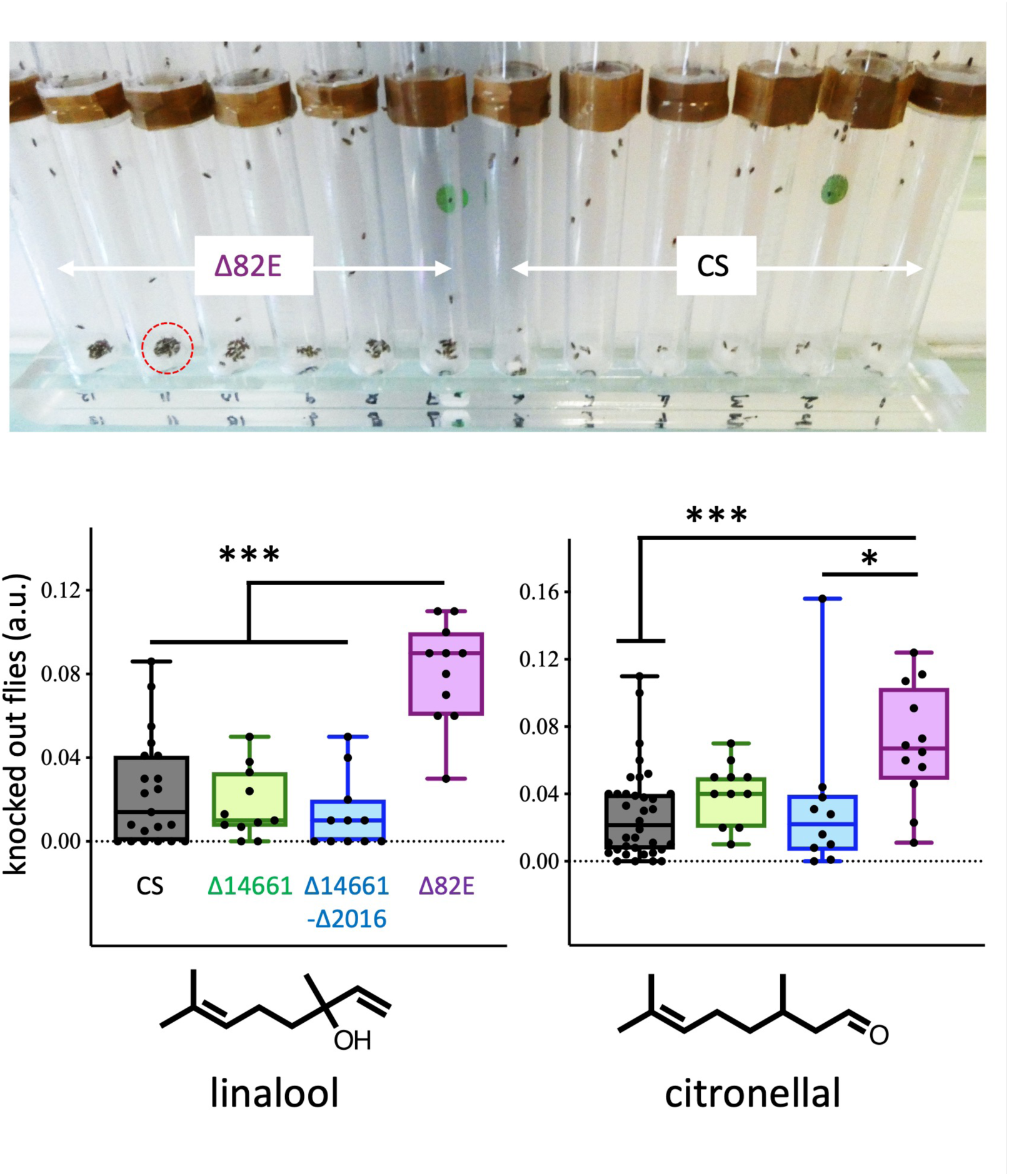
Δ82 mutants are highly sensitive to linalool and citronellal exposure. The image shows the bottom of tubes at the end of a DART experiment with citronellal. Note the greater number of knocked out or dead Δ82E flies (circle with red dashed line), as compared with control (Canton S = CS) flies. In the histograms below, the values for the Y axis reflect the surface of an image occupied by knocked out flies, measured at high magnification after calibration with an internal size reference, using imageJ (Schindelin et al., 2012). A significant difference (P<0.0005 (***), P<0.05 (*), n=9-36, ANOVA, Kruskal-Wallis test with Dunn’s post-hoc correction) is observed between Δ82E mutants relative to CS and other genotypes. Δ14661 and Δ14661-2016 flies are not significantly more sensitive than CS flies. a.u: arbitrary surface units.

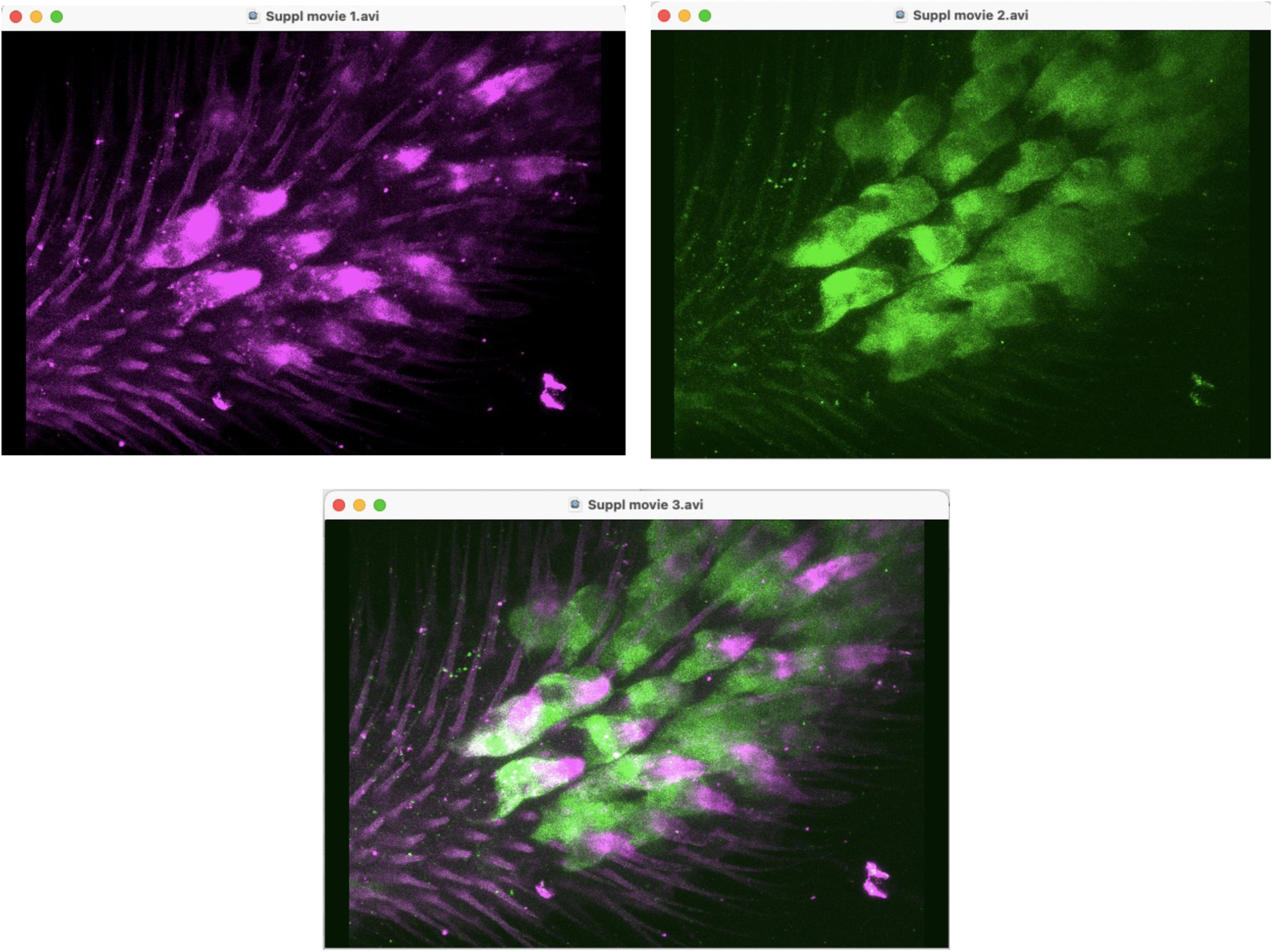

**Supplementary movies 1-3:** These movies were made with the ImageJ build-in 3D project tool (Schindelin et al., 2012) from a stack of 30 confocal slices 1 µM apart. They show a few basiconic sensillae of the dorsal surface of the maxillary palp, from which Figure 5D was drawn.

Suppl movie 1: Localization of the 14661 3XFLAG protein.

Suppl movie 2: Localization of the Mi14661 gene trap GFP reporter.

Suppl movie 3: Merge of movies 1 and 2.

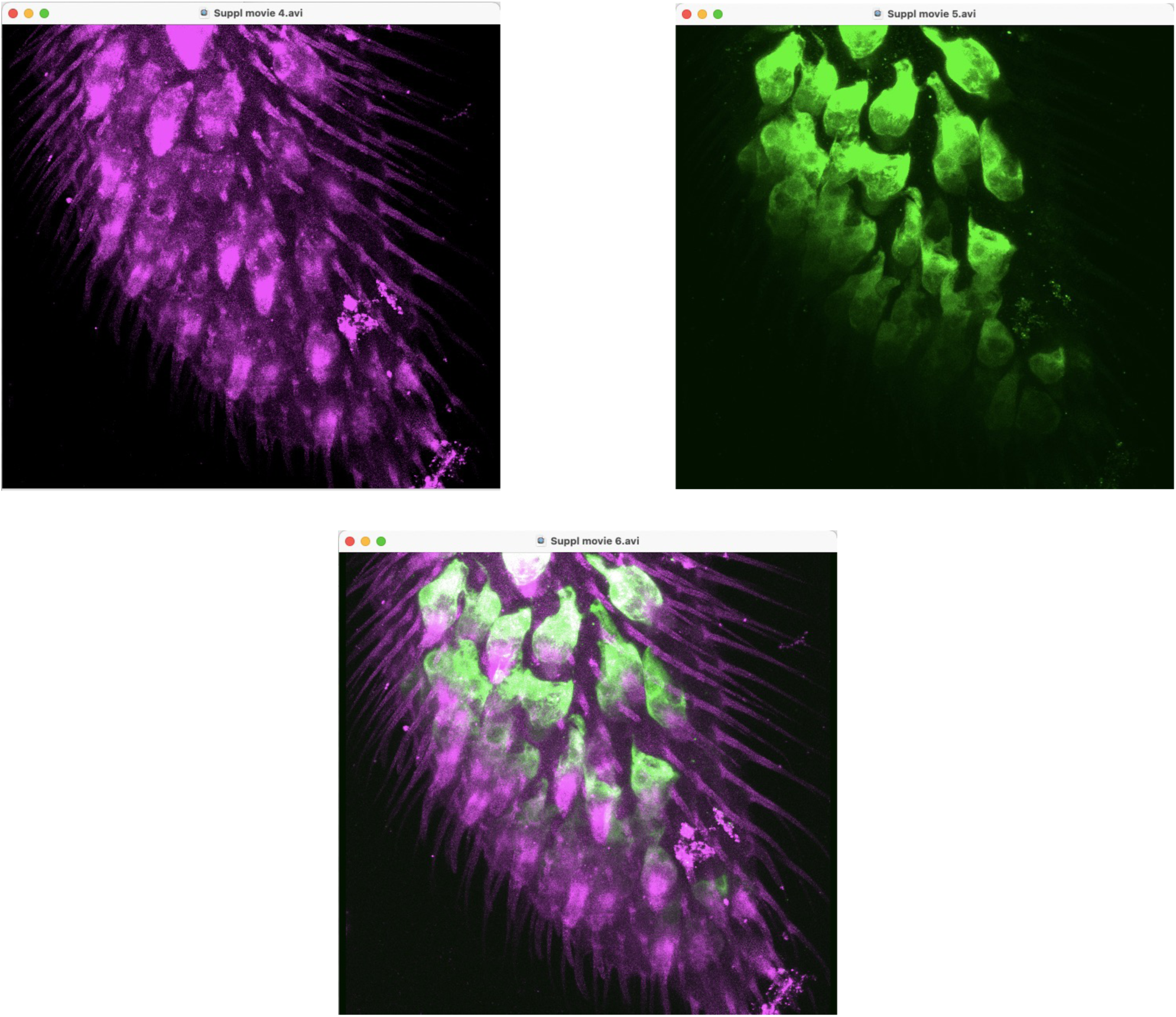

**Supplementary movies 4-6:** These movies were constructed from a stack of 16 confocal slices 1.5 µM apart, showing a few basiconic sensillae of the dorsal surface of the maxillary palp, from which Figure 5F was drawn.

Suppl movie 4: Localization of the 14661 3XFLAG protein.

Suppl movie 5: Localization of the Mi2016 gene trap GFP reporter.

Suppl movie 6: Merge of movies 1 and 2.

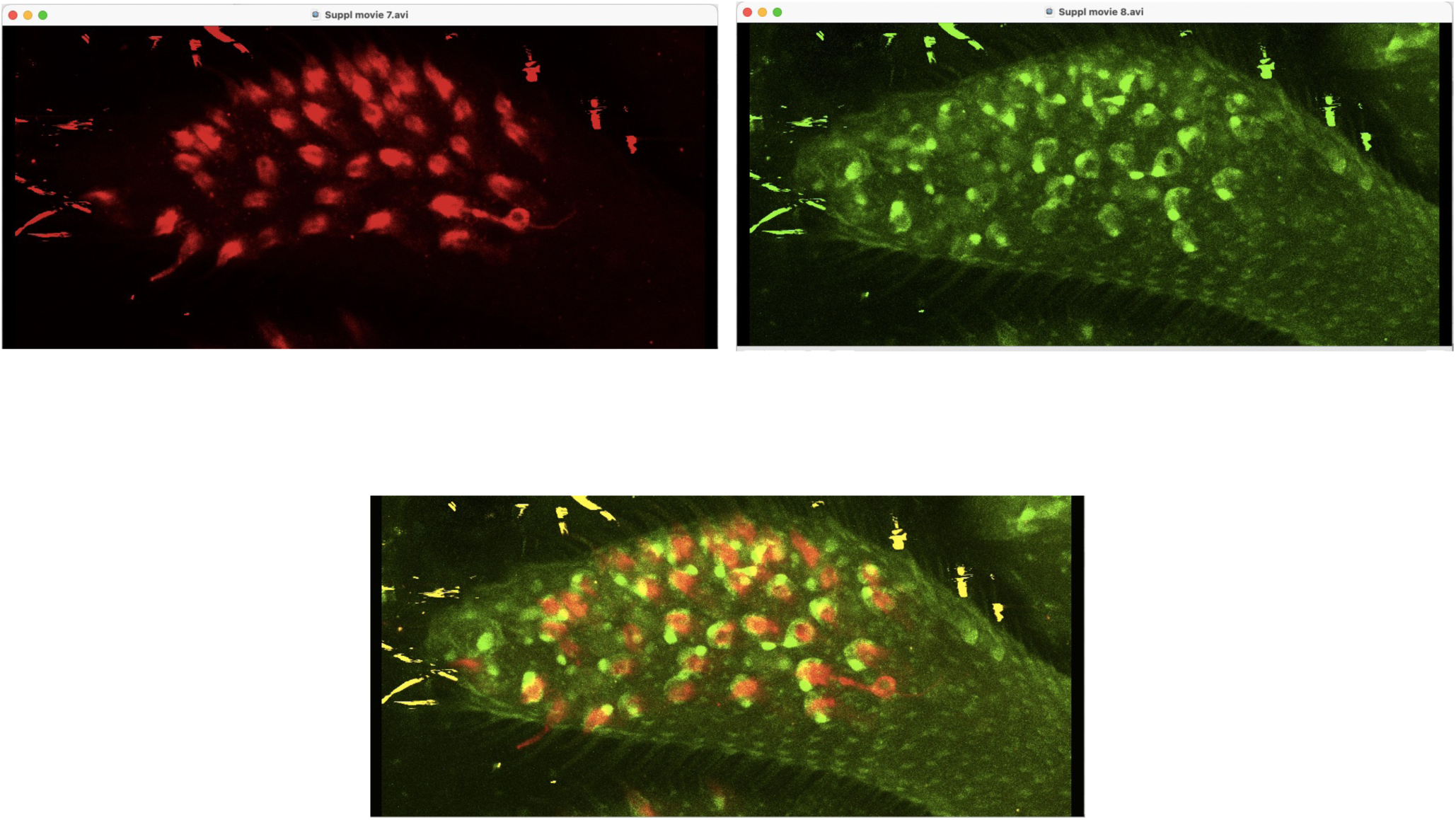

**Supplementary movies 7-9:** These movies were constructed from a stack of 32 confocal slices 1.5 µM slices apart. They show an entire maxillary palp and correspond to Figure 5H.

Suppl movie 7: Localization of the 14661 3XFLAG protein. Note the presence of a neuron expressing CG14661 at the beginning of the movie, at the bottom right-hand side. A dendrite emanating from this neuron apparently enters inside a basiconic sensilla.

Suppl movie 8: Localization of GFP driven by the Ase5 element, specifically active in the tormogen cell.

Suppl movie 9: Merge of movies 1 and 2.

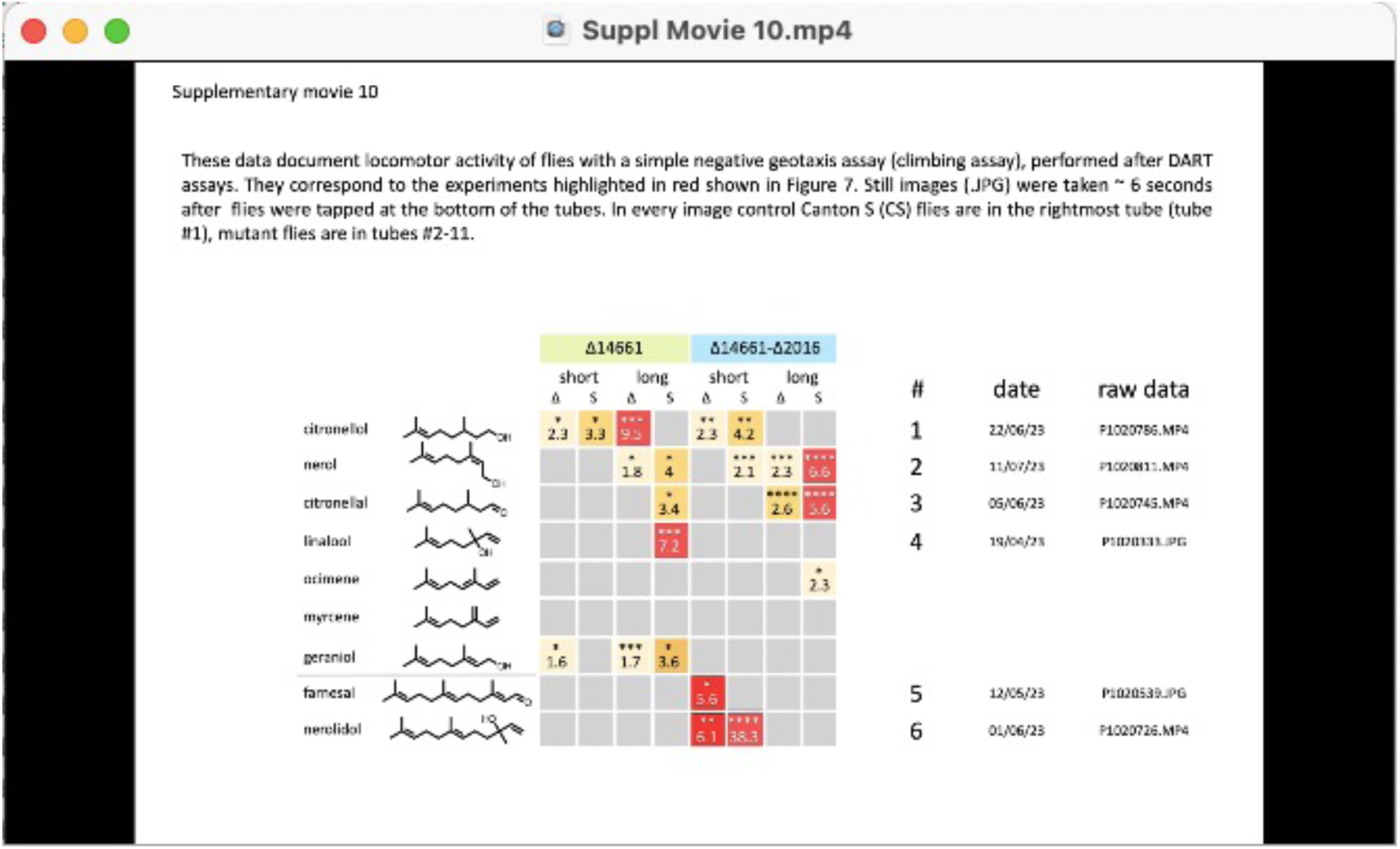

**Suppl movie 10:** Movie clips and still images of climbing assays for molecules eliciting the most acute responses (highlighted in red in Figure 7, and corresponding to fold changes >5). In all experiments the rightmost tube contains control Canton S (CS) flies. Note that the data essentially shows that mutant flies do not appear to display significantly enhanced locomotory behavior relative to CS flies.

**Supplementary Table 1:**
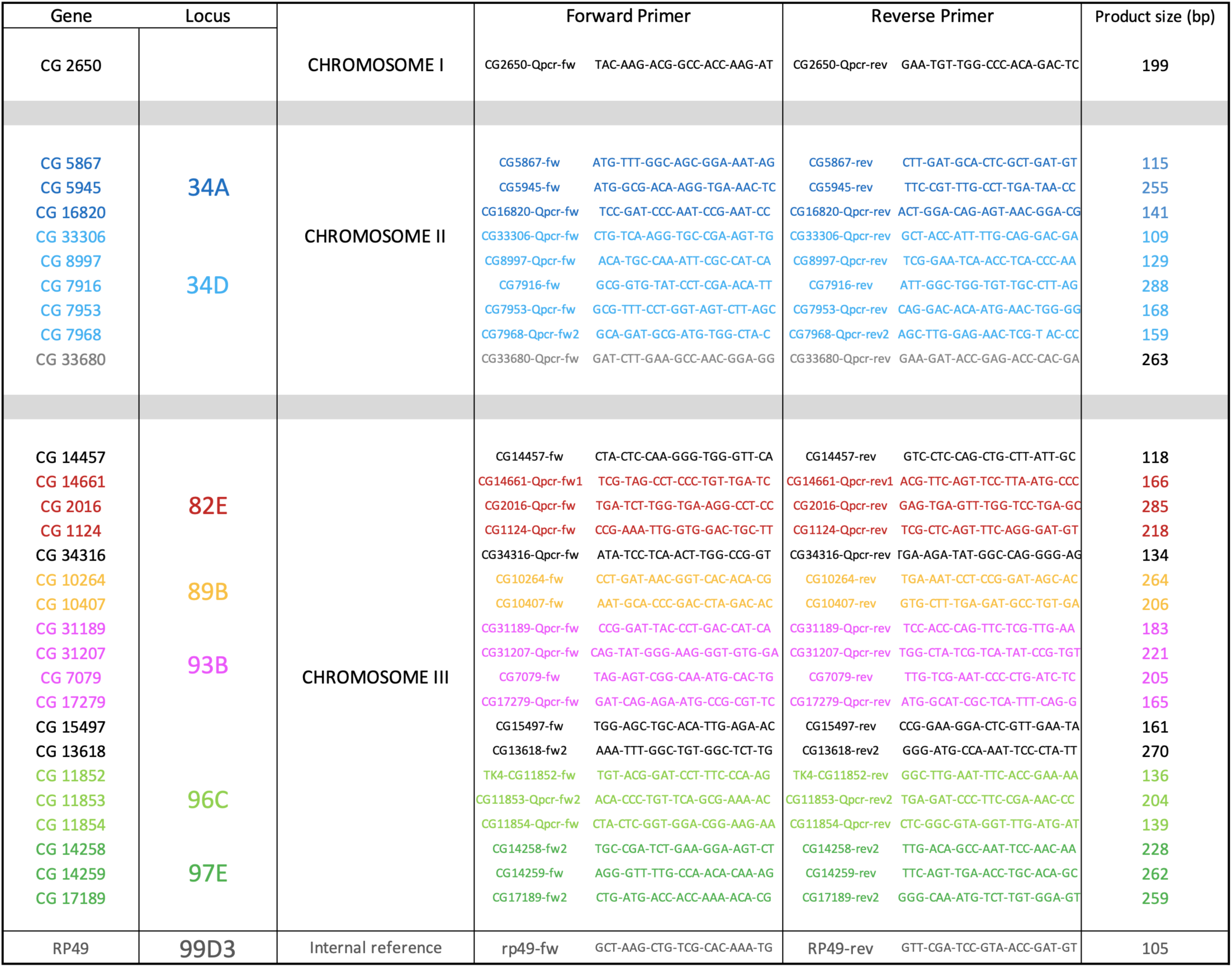
Q-PCR primers for Drosophila B-TDPs.

**Supplementary Table 2:**
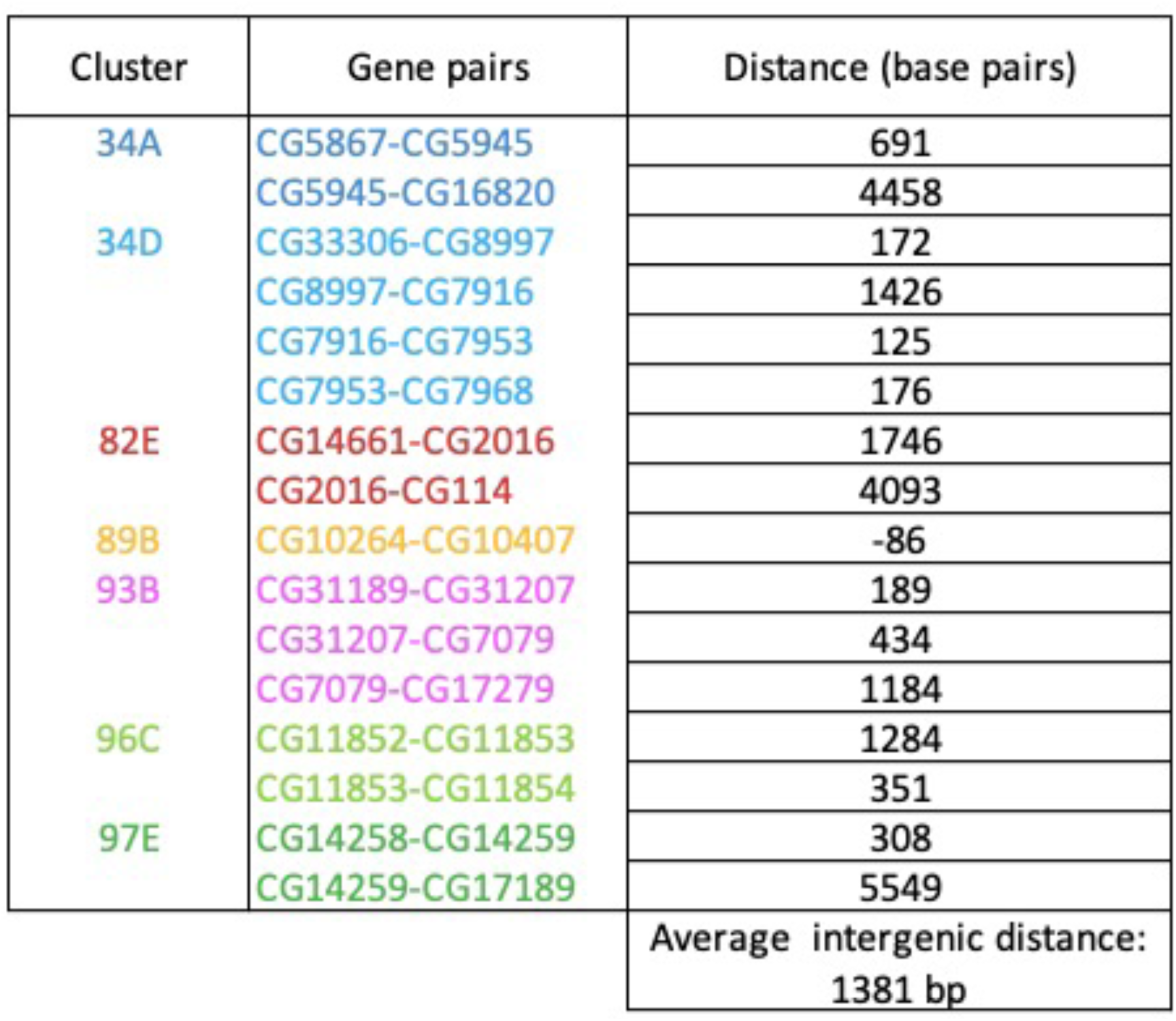
Intergenic distances in *Drosophila* B-TDP clusters.

**Supplementary Table 3 :**
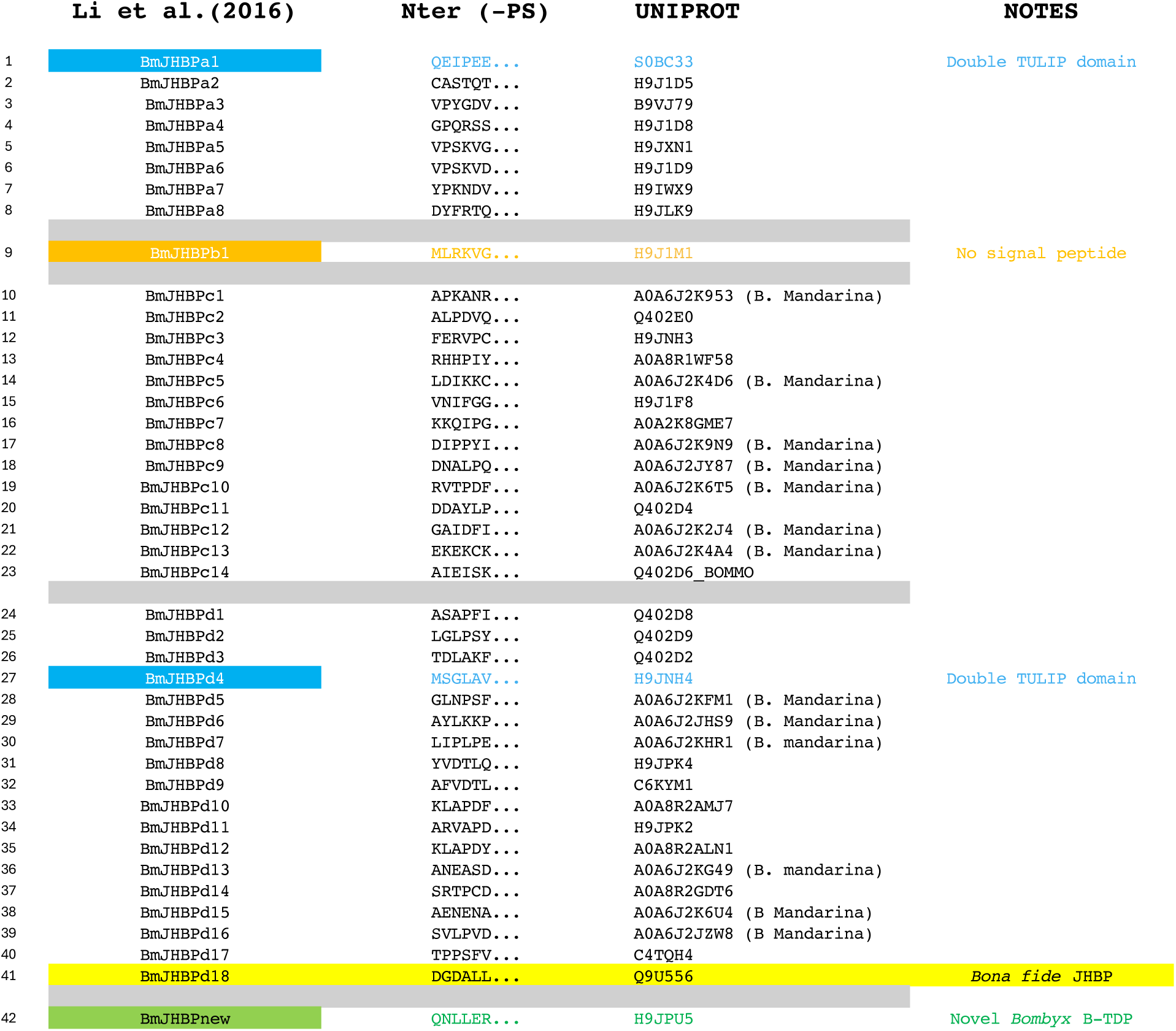
List of 42 *Bombyx mori* B-TDPs (Li et al., 2016) and UNIPROT accession numbers. Note that only one of these proteins has been demonstrated to bind Juvenile Hormones with high affinity (BmJHBPd18, (Kurata et al., 1994; Vermunt et al., 2001)) and is referred to as *bona fide* JHBP. Two pseudodimeric *Bombyx* proteins with two TULIP domains (BmJHBPd4 and BmJHBPa1) were not included in the alignment used for generating the phylogenetic tree of Figure 2.

**Supplementary Table 4:** List of odorants used in the study.

| Compound | Reference | Purity | Source |
| --- | --- | --- | --- |
| Benzaldehyde | B1334-250ML | ≥99% | Sigma |
| trans-2-Hexen-1-al | W256005-SAMPLE-K | ≥95% | Sigma |
| Phenylethanol | not recorded |  |  |
| β-myrcene | M100005-5ML | >75% | Sigma |
| Ocimene, mixture of isomers | W353901-SAMPLE | >90% | Sigma |
| Linalool | L2602-5G | > 96.5 % | Sigma |
| Geraniol | 163333-25G | > 97.5 % | Sigma |
| Citronellol | W230915 | > 95.5 % | Sigma |
| Citronellal | 27470 | ≥95.0% | Sigma |
| Farnesal, mixture of isomers | 46188-1ML-F | >= 85 % | Sigma |
| Trans,trans-Farnesol | 277541-1G | > 95.5 % | Sigma |
| Nerol | A19808 | 97% | Thermoscientific |
| Nerolidol | 20661 | 97% | Pherobank |
| (E)-beta Farnesene | 20029 | 89,50% | Pherobank |
| Farnesene, mixture of isomers | W383902-SAMPLE-K | NA | Sigma |
| Alpha farnesene (mixture of isomers) | 20027 | 90 | Pherobank |
| DMSO | 472301-500ML | ≥99.9% | Sigma |

## 6. Acknowledgments

This work was supported by INRAE, CNRS, Université Bourgogne Europe, CSGA, and in its early stage by a grant from Région Bourgogne-Franche-Comté to JPC.

## Notes

### Competing Interest Statement

The authors have declared no competing interest.

### Summary of Updates

- A few typos were corrected - Some rewording was done in several places. - Funding information was updated

## References

Almagro Armenteros, Salvatore M, Emanuelsson O, Winther O, Von Heijne G, Elofsson A, Nielsen H. 2019a. Detecting sequence signals in targeting peptides using deep learning. Life Science Alliance 2:e201900429. DOI: 10.26508/lsa.201900429

Almagro Armenteros, Tsirigos KD, Sønderby CK, Petersen TN, Winther O, Brunak S, Von Heijne G, Nielsen H. 2019b. SignalP 5.0 improves signal peptide predictions using deep neural networks. Nature Biotechnology 37:420–423. DOI: 10.1038/s41587-019-0036-z

Alva V, Lupas AN. 2016. The TULIP superfamily of eukaryotic lipid-binding proteins as a mediator of lipid sensing and transport. Biochim Biophys Acta 1861:913–23. DOI: 10.1016/j.bbalip.2016.01.016

Angeli S, Ceron F, Scaloni A, Monti M, Monteforti G, Minnocci A, Petacchi R, Pelosi P. 1999. Purification, structural characterization, cloning and immunocytochemical localization of chemoreception proteins from *Schistocerca gregaria*. European Journal of Biochemistry 262:745–754. DOI: 10.1046/j.1432-1327.1999.00438.x

Bakkali F, Averbeck S, Averbeck D, Idaomar M. 2008. Biological effects of essential oils – A review. Food and Chemical Toxicology 46:446–475. DOI: 10.1016/j.fct.2007.09.106

Baldwin IT. 2010. Plant volatiles. Curr Biol 20:R392–7. DOI: 10.1016/j.cub.2010.02.052

Barolo S, Walker RG, Polyanovsky AD, Freschi G, Keil T, Posakony JW. 2000. A Notch-Independent Activity of Suppressor of Hairless Is Required for Normal Mechanoreceptor Physiology. Cell 103:957–970. DOI: 10.1016/S0092-8674(00)00198-7

Bartlett J, Gakhar L, Penterman J, Singh P, Mallampalli RK, Porter E, McCray PB. 2011. PLUNC: a multifunctional surfactant of the airways. Biochemical Society Transactions 39:1012–1016. DOI: 10.1042/BST0391012

Beamer LJ, Carroll SF, Eisenberg D. 1997. Crystal Structure of Human BPI and Two Bound Phospholipids at 2.4 Angstrom Resolution. Science 276:1861–1864. DOI: 10.1126/science.276.5320.1861

Bhattacharya AA, Grüne T, Curry S. 2000. Crystallographic analysis reveals common modes of binding of medium and long-chain fatty acids to human serum albumin 1 1Edited by R. Huber. Journal of Molecular Biology 303:721–732. DOI: 10.1006/jmbi.2000.4158

Bingle CD, Bingle L, Craven CJ. 2011. Distant cousins: genomic and sequence diversity within the BPI fold-containing (BPIF)/PLUNC protein family. Biochem Soc Trans 39:961–5. DOI: 10.1042/BST0390961

Bingle L, Bingle CD. 2011. Distribution of human PLUNC/BPI fold-containing (BPIF) proteins. Biochem Soc Trans 39:1023–7. DOI: 10.1042/BST0391023

Bingle, Seal RL, Craven CJ. 2011. Systematic nomenclature for the PLUNC/PSP/BSP30/SMGB proteins as a subfamily of the BPI fold-containing superfamily. Biochemical Society Transactions 39:977–983. DOI: 10.1042/BST0390977

Bleeker PM, Mirabella R, Diergaarde PJ, VanDoorn A, Tissier A, Kant MR, Prins M, De Vos M, Haring MA, Schuurink RC. 2012. Improved herbivore resistance in cultivated tomato with the sesquiterpene biosynthetic pathway from a wild relative. Proceedings of the National Academy of Sciences 109:20124–20129. DOI: 10.1073/pnas.1208756109

Blum M, Andreeva A, Florentino LC, Chuguransky SR, Grego T, Hobbs E, Pinto BL, Orr A, Paysan-Lafosse T, Ponamareva I, Salazar GA, Bordin N, Bork P, Bridge A, Colwell L, Gough J, Haft DH, Letunic I, Llinares-López F, Marchler-Bauer A, Meng-Papaxanthos L, Mi H, Natale DA, Orengo CA, Pandurangan AP, Piovesan D, Rivoire C, Sigrist CJA, Thanki N, Thibaud-Nissen F, Thomas PD, Tosatto SCE, Wu CH, Bateman A. 2025. InterPro: the protein sequence classification resource in 2025. Nucleic Acids Research 53:D444–D456. DOI: 10.1093/nar/gkae1082

Bohbot J, Vogt RG. 2005. Antennal expressed genes of the yellow fever mosquito (Aedes aegypti L.); characterization of odorant-binding protein 10 and takeout. Insect Biochem Mol Biol **35**:961–79. DOI: 10.1016/j.ibmb.2005.03.010

Boncan DAT, Tsang SSK, Li C, Lee IHT, Lam H-M, Chan T-F, Hui JHL. 2020. Terpenes and Terpenoids in Plants: Interactions with Environment and Insects. International Journal of Molecular Sciences 21:7382. DOI: 10.3390/ijms21197382

Brito NF, Moreira MF, Melo ACA. 2016. A look inside odorant-binding proteins in insect chemoreception. Journal of Insect Physiology 95:51–65. DOI: 10.1016/j.jinsphys.2016.09.008

Bruce TJA. 2015. Interplay between insects and plants: dynamic and complex interactions that have coevolved over millions of years but act in milliseconds. Journal of Experimental Botany 66:455–465. DOI: 10.1093/jxb/eru391

Bruce TJA, Pickett JA. 2011. Perception of plant volatile blends by herbivorous insects – Finding the right mix. Phytochemistry 72:1605–1611. DOI: 10.1016/j.phytochem.2011.04.011

Bruce TJA, Wadhams LJ, Woodcock CM. 2005. Insect host location: a volatile situation. Trends in Plant Science 10:269–274. DOI: 10.1016/j.tplants.2005.04.003

Charbonneau DM, Tajmir-Riahi H-A. 2010. Study on the Interaction of Cationic Lipids with Bovine Serum Albumin. The Journal of Physical Chemistry B 114:1148–1155. DOI: 10.1021/jp910077h

Charles J-P, Wojtasek H, Lentz AJ, Thomas BA, Bonning BC, Palli SR, Parker AG, Dorman G, Hammock BD, Prestwich GD, Riddiford LM. 1996. Purification and reassessment of ligand binding by the recombinant, putative juvenile hormone receptor of the tobacco hornworm, Manduca sexta. Archives of Insect Biochemistry and Physiology 31:371–393. DOI: 10.1002/(SICI)1520-6327(1996)31:4<371::AID-ARCH2>3.0.CO;2-Z

Chiapparino A, Maeda K, Turei D, Saez-Rodriguez J, Gavin A-C. 2016. The orchestra of lipid-transfer proteins at the crossroads between metabolism and signaling. Progress in Lipid Research 61:30–39. DOI: 10.1016/j.plipres.2015.10.004

Chung YD, Zhu J, Han Y-G, Kernan MJ. 2001. nompA Encodes a PNS-Specific, ZP Domain Protein Required to Connect Mechanosensory Dendrites to Sensory Structures. Neuron 29:415–428. DOI: 10.1016/S0896-6273(01)00215-X

Corcoran JA, Jordan MD, Thrimawithana AH, Crowhurst RN, Newcomb RD. 2015. The Peripheral Olfactory Repertoire of the Lightbrown Apple Moth, Epiphyas postvittana. PLoS One 10:e0128596. DOI: 10.1371/journal.pone.0128596

Cordeiro CMM, Esmaili H, Ansah G, Hincke MT. 2013. Ovocalyxin-36 Is a Pattern Recognition Protein in Chicken Eggshell Membranes. PLoS ONE 8:e84112. DOI: 10.1371/journal.pone.0084112

Curry S, Mandelkow H, Brick P, Franks N. 1998. Crystal structure of human serum albumin complexed with fatty acid reveals an asymmetric distribution of binding sites. Nature Structural Biology 5:827–835. DOI: 10.1038/1869

Dauwalder B, Tsujimoto S, Moss J, Mattox W. 2002. The Drosophila takeout gene is regulated by the somatic sex-determination pathway and affects male courtship behavior. Genes & Development 16:2879–2892. DOI: 10.1101/gad.1010302, PMID: 12435630

De Groot AC, Schmidt E. 2016. Essential Oils, Part III: Chemical Composition. Dermatitis 27:161–169. DOI: 10.1097/DER.0000000000000193

Dudareva N, Negre F, Nagegowda DA, Orlova I. 2006. Plant Volatiles: Recent Advances and Future Perspectives. Critical Reviews in Plant Sciences 25:417–440. DOI: 10.1080/07352680600899973

Dupas S, Neiers F, Granon E, Rougeux E, Dupont S, Beney L, Bousquet F, Shaik HA, Briand L, Wojtasek H, Charles JP. 2020. Collisional mechanism of ligand release by Bombyxmori JHBP, a member of the TULIP / Takeout family of lipid transporters. Insect Biochem Mol Biol 117:103293. DOI: 10.1016/j.ibmb.2019.103293

Egea PF. 2021. Mechanisms of Non-Vesicular Exchange of Lipids at Membrane Contact Sites: Of Shuttles, Tunnels and, Funnels. Frontiers in Cell and Developmental Biology 9:784367. DOI: 10.3389/fcell.2021.784367

Eisenberg D, Schwarz E, Komaromy M, Wall R. 1984. Analysis of membrane and surface protein sequences with the hydrophobic moment plot. Journal of Molecular Biology 179:125–142. DOI: 10.1016/0022-2836(84)90309-7, PMID: 6502707

Finetti L, Tiedemann L, Zhang X, Civolani S, Bernacchia G, Roeder T. 2020. Monoterpenes alter TAR1-driven physiology in *Drosophila* species. Journal of Experimental Biology jeb.232116. DOI: 10.1242/jeb.232116

Fujikawa K, Seno K, Ozaki M. 2006. A novel Takeout-like protein expressed in the taste and olfactory organs of the blowfly, Phormia regina. FEBS J 273:4311–21. DOI: 10.1111/j.1742-4658.2006.05422.x

Ganetzky B, Flanagan JR. 1978. On the relationship between senescence and age-related changes in two wild-type strains of Drosophila melanogaster. Experimental Gerontology 13:189–196. DOI: 10.1016/0531-5565(78)90012-8

Gargano J, Martin I, Bhandari P, Grotewiel M. 2005. Rapid iterative negative geotaxis (RING): a new method for assessing age-related locomotor decline in. Experimental Gerontology 40:386–395. DOI: 10.1016/j.exger.2005.02.005

Gautron J, Réhault-Godbert S, Pascal G, Nys Y, Hincke MT. 2011. Ovocalyxin-36 and other LBP/BPI/PLUNC-like proteins as molecular actors of the mechanisms of the avian egg natural defences. Biochemical Society Transactions 39:971–976. DOI: 10.1042/BST0390971

Genter MB, Van Veldhoven PP, Jegga AG, Sakthivel B, Kong S, Stanley K, Witte DP, Ebert CL, Aronow BJ. 2003. Microarray-based discovery of highly expressed olfactory mucosal genes: potential roles in the various functions of the olfactory system. Physiol Genomics 16:67–81. DOI: 10.1152/physiolgenomics.00117.2003

Gibson CW, Thomson NH, Abrams WR, Kirkham J. 2005. Nested genes: Biological implications and use of AFM for analysis. Gene 350:15–23. DOI: 10.1016/j.gene.2004.12.045

Goodman WG. 1990. Biosynthesis, titer regulation and transport of juvenile hormones. Morphogenetic Hormones of Arthropods. Rutgers Uniiversity Press, New Brunswick and London. p. 83–124.

Gramates LS, Agapite J, Attrill H, Calvi BR, Crosby MA, Dos Santos Gilberto, Goodman JL, Goutte-Gattat D, Jenkins VK, Kaufman T, Larkin A, Matthews BB, Millburn G, Strelets VB, the FlyBase Consortium, Perrimon N, Gelbart SR, Agapite J, Broll K, Crosby L, Dos Santos Gil, Falls K, Gramates LS, Jenkins V, Longden I, Matthews B, Seme J, Tabone CJ, Zhou P, Zytkovicz M, Brown N, Antonazzo G, Attrill H, Garapati P, Goutte-Gattat D, Larkin A, Marygold S, McLachlan A, Millburn G, Öztürk-Çolak A, Pilgrim C, Trovisco V, Calvi B, Kaufman T, Goodman J, Krishna P, Strelets V, Thurmond J, Cripps R, Lovato T. 2022. FlyBase: a guided tour of highlighted features. Genetics **220**:iyac035. DOI: 10.1093/genetics/iyac035

Gratz SJ, Cummings AM, Nguyen JN, Hamm DC, Donohue LK, Harrison MM, Wildonger J, O’Connor-Giles KM. 2013. Genome Engineering of *Drosophila* with the CRISPR RNA-Guided Cas9 Nuclease. Genetics 194:1029–1035. DOI: 10.1534/genetics.113.152710

Gratz SJ, Rubinstein CD, Harrison MM, Wildonger J, O’Connor-Giles KM. 2015. CRISPR-Cas9 Genome Editing in Drosophila. Curr Protoc Mol Biol 111:31 2 1-31 2 20. DOI: 10.1002/0471142727.mb3102s111

Gratz SJ, Ukken FP, Rubinstein CD, Thiede G, Donohue LK, Cummings AM, O’Connor-Giles KM. 2014. Highly specific and efficient CRISPR/Cas9-catalyzed homology-directed repair in Drosophila. Genetics 196:961–71. DOI: 10.1534/genetics.113.160713

Gu SH, Wang SP, Zhang XY, Wu KM, Guo YY, Zhou JJ, Zhang YJ. 2011. Identification and tissue distribution of odorant binding protein genes in the lucerne plant bug Adelphocoris lineolatus (Goeze). Insect Biochem Mol Biol 41:254–63. DOI: 10.1016/j.ibmb.2011.01.002

Hagai T, Cohen M, Bloch G. 2007. Genes encoding putative Takeout/juvenile hormone binding proteins in the honeybee (Apis mellifera) and modulation by age and juvenile hormone of the takeout-like gene. Insect Biochemistry and Molecular Biology 37:689–701. DOI: 10.1016/j.ibmb.2007.04.002

Hamiaux C, Basten L, Greenwood DR, Baker EN, Newcomb RD. 2013. Ligand promiscuity within the internal cavity of Epiphyas postvittana Takeout 1 protein. J Struct Biol 182:259–63. DOI: 10.1016/j.jsb.2013.03.013

Hamiaux C, Stanley D, Greenwood DR, Baker EN, Newcomb RD. 2009. Crystal structure of Epiphyas postvittana takeout 1 with bound ubiquinone supports a role as ligand carriers for takeout proteins in insects. J Biol Chem 284:3496–503. DOI: 10.1074/jbc.M807467200

Hekmat-Scafe DS, Scafe CR, McKinney AJ, Tanouye MA. 2002. Genome-Wide Analysis of the Odorant-Binding Protein Gene Family in *Drosophila melanogaster*. Genome Research 12:1357–1369. DOI: 10.1101/gr.239402

Hekmat-Scafe DS, Steinbrecht RA, Carlson JR. 1997. Coexpression of Two Odorant-Binding Protein Homologs in *Drosophila* : Implications for Olfactory Coding. The Journal of Neuroscience 17:1616–1624. DOI: 10.1523/JNEUROSCI.17-05-01616.1997

Himmel NJ, Moi D, Benton R. 2023. Remote homolog detection places insect chemoreceptors in a cryptic protein superfamily spanning the tree of life. Current biology: CB 33:5023–5033.e4. DOI: 10.1016/j.cub.2023.10.008, PMID: 37913770

Hojo M, Morioka M, Matsumoto T, Miura T. 2005. Identification of soldier caste-specific protein in the frontal gland of nasute termite Nasutitermes takasagoensis (Isoptera: Termitidae). Insect Biochem Mol Biol 35:347–54. DOI: 10.1016/j.ibmb.2005.01.007

Horlbeck MA, Witkowsky LB, Guglielmi B, Replogle JM, Gilbert LA, Villalta JE, Torigoe SE, Tjian R, Weissman JS. 2016. Nucleosomes impede Cas9 access to DNA in vivo and in vitro. Elife 5. DOI: 10.7554/eLife.12677

Illergård K, Ardell DH, Elofsson A. 2009. Structure is three to ten times more conserved than sequence—A study of structural response in protein cores. Proteins: Structure, Function, and Bioinformatics 77:499–508. DOI: 10.1002/prot.22458

Isman MB. 2006. BOTANICAL INSECTICIDES, DETERRENTS, AND REPELLENTS IN MODERN AGRICULTURE AND AN INCREASINGLY REGULATED WORLD. Annual Review of Entomology 51:45–66. DOI: 10.1146/annurev.ento.51.110104.151146

Joseph RM, Carlson JR. 2015. Drosophila Chemoreceptors: A Molecular Interface Between the Chemical World and the Brain. Trends in genetics: TIG 31:683–695. DOI: 10.1016/j.tig.2015.09.005, PMID: 26477743

Jumper J, Evans R, Pritzel A, Green T, Figurnov M, Ronneberger O, Tunyasuvunakool K, Bates R, Žídek A, Potapenko A, Bridgland A, Meyer C, Kohl SAA, Ballard AJ, Cowie A, Romera-Paredes B, Nikolov S, Jain R, Adler J, Back T, Petersen S, Reiman D, Clancy E, Zielinski M, Steinegger M, Pacholska M, Berghammer T, Bodenstein S, Silver D, Vinyals O, Senior AW, Kavukcuoglu K, Kohli P, Hassabis D. 2021. Highly accurate protein structure prediction with AlphaFold. Nature 596:583–589. DOI: 10.1038/s41586-021-03819-2

Justice RW, Dimitratos S, Walter MF, Woods DF, Biessmann H. 2003. Sexual dimorphic expression of putative antennal carrier protein genes in the malaria vector Anopheles gambiae. Insect Mol Biol 12:581–94. DOI: 10.1046/j.1365-2583.2003.00443.x

Katoh K, Rozewicki J, Yamada KD. 2019. MAFFT online service: multiple sequence alignment, interactive sequence choice and visualization. Briefings in Bioinformatics 20:1160–1166. DOI: 10.1093/bib/bbx108

Kawano S, Tamura Y, Kojima R, Bala S, Asai E, Michel AH, Kornmann B, Riezman I, Riezman H, Sakae Y, Okamoto Y, Endo T. 2018. Structure–function insights into direct lipid transfer between membranes by Mmm1–Mdm12 of ERMES. Journal of Cell Biology 217:959–974. DOI: 10.1083/jcb.201704119

Kendroud S, Bohra AA, Kuert PA, Nguyen B, Guillermin O, Sprecher SG, Reichert H, VijayRaghavan K, Hartenstein V. 2018. Structure and development of the subesophageal zone of the *Drosophila* brain. II. Sensory compartments. Journal of Comparative Neurology 526:33–58. DOI: 10.1002/cne.24316

Kitamura–Abe S, Itoh H, Washio T, Tsutsumi A, Tomita M. 2004. CHARACTERIZATION OF THE SPLICE SITES IN GT–AG AND GC–AG INTRONS IN HIGHER EUKARYOTES USING FULL-LENGTH cDNAs. Journal of Bioinformatics and Computational Biology **02**:309–331. DOI: 10.1142/S0219720004000570

Kolodziejczyk R, Bujacz G, Jakób M, Ozyhar A, Jaskolski M, Kochman M. 2008. Insect juvenile hormone binding protein shows ancestral fold present in human lipid-binding proteins. Journal of Molecular Biology 377:870–881. DOI: 10.1016/j.jmb.2008.01.026, PMID: 18291417

Kopec KO, Alva V, Lupas AN. 2011. Bioinformatics of the TULIP domain superfamily. Biochem Soc Trans 39:1033–8. DOI: 10.1042/BST0391033

Kopec KO, Alva V, Lupas AN. 2010. Homology of SMP domains to the TULIP superfamily of lipid-binding proteins provides a structural basis for lipid exchange between ER and mitochondria. Bioinformatics 26:1927–31. DOI: 10.1093/bioinformatics/btq326

Kornmann B, Currie E, Collins SR, Schuldiner M, Nunnari J, Weissman JS, Walter P. 2009. An ER-mitochondria tethering complex revealed by a synthetic biology screen. Science 325:477–81. DOI: 10.1126/science.1175088

Krasity BC, Troll JV, Weiss JP, McFall-Ngai MJ. 2011. LBP/BPI proteins and their relatives: conservation over evolution and roles in mutualism. Biochemical Society Transactions 39:1039–1044. DOI: 10.1042/BST0391039

Kroeger T, Frieg B, Zhang T, Hansen FK, Marmann A, Proksch P, Nagel-Steger L, Groth G, Smits SHJ, Gohlke H. 2017. EDTA aggregates induce SYPRO orange-based fluorescence in thermal shift assay. PLOS ONE 12:e0177024. DOI: 10.1371/journal.pone.0177024

Kurata K, Nakamura M, Okuda T, Hirano H, Shinbo H. 1994. Purification and characterization of a juvenile hormone binding protein from hemolymph of the silkworm, Bombyx mori. Comparative Biochemistry and Physiology Part B: Comparative Biochemistry 109:105–114. DOI: 10.1016/0305-0491(94)90147-3

Kwon Y, Kim SH, Ronderos DS, Lee Y, Akitake B, Woodward OM, Guggino WB, Smith DP, Montell C. 2010. Drosophila TRPA1 channel is required to avoid the naturally occurring insect repellent citronellal. Curr Biol 20:1672–8. DOI: 10.1016/j.cub.2010.08.016

Labandeira CC. 2013. A paleobiologic perspective on plant–insect interactions. Current Opinion in Plant Biology 16:414–421. DOI: 10.1016/j.pbi.2013.06.003

Larter NK, Sun JS, Carlson JR. 2016. Organization and function of Drosophila odorant binding proteins. eLife 5:e20242. DOI: 10.7554/eLife.20242

Lee I, Hong W. 2006. Diverse membrane-associated proteins contain a novel SMP domain. The FASEB Journal 20:202–206. DOI: 10.1096/fj.05-4581hyp

Lee MY. 2018. Essential Oils as Repellents against Arthropods. BioMed Research International 2018:1–9. DOI: 10.1155/2018/6860271

Levine TP. 2019. Remote homology searches identify bacterial homologues of eukaryotic lipid transfer proteins, including Chorein-N domains in TamB and AsmA and Mdm31p. BMC Molecular and Cell Biology 20:43. DOI: 10.1186/s12860-019-0226-z

Li W, Cheng T, Hu W, Peng Z, Liu C, Xia Q. 2016. Genome-wide identification and analysis of JHBP-domain family members in the silkworm Bombyx mori. Molecular Genetics and Genomics 291:2159–2171. DOI: 10.1007/s00438-016-1245-5

Li-Kroeger D, Kanca O, Lee PT, Cowan S, Lee MT, Jaiswal M, Salazar JL, He Y, Zuo Z, Bellen HJ. 2018. An expanded toolkit for gene tagging based on MiMIC and scarless CRISPR tagging in Drosophila. Elife 7. DOI: 10.7554/eLife.38709

Liu Y-L, Guo H, Huang L-Q, Pelosi P, Wang C-Z. 2014. Unique function of a chemosensory protein in the proboscis of two *Helicoverpa* species. Journal of Experimental Biology jeb.102020. DOI: 10.1242/jeb.102020

Livak KJ, Schmittgen TD. 2001. Analysis of Relative Gene Expression Data Using Real-Time Quantitative PCR and the 2−ΔΔCT Method. Methods 25:402–408. DOI: 10.1006/meth.2001.1262

Madeira F, Madhusoodanan N, Lee J, Eusebi A, Niewielska A, Tivey ARN, Lopez R, Butcher S. 2024. The EMBL-EBI Job Dispatcher sequence analysis tools framework in 2024. Nucleic acids research gkae241. DOI: 10.1093/nar/gkae241, PMID: 38597606

Maffei ME, Gertsch J, Appendino G. 2011. Plant volatiles: production, function and pharmacology. Nat Prod Rep 28:1359–80. DOI: 10.1039/c1np00021g

Malnic B, Hirono J, Sato T, Buck LB. 1999. Combinatorial Receptor Codes for Odors. Cell 96:713–723. DOI: 10.1016/S0092-8674(00)80581-4

Martin F, Boto T, Gomez-Diaz C, Alcorta E. 2013. Elements of Olfactory Reception in Adult *Drosophila melanogaster*. The Anatomical Record 296:1477–1488. DOI: 10.1002/ar.22747

Masson D, Jiang X-C, Lagrost L, Tall AR. 2009. The role of plasma lipid transfer proteins in lipoprotein metabolism and atherogenesis. Journal of Lipid Research 50 **Suppl**:S201-206. DOI: 10.1194/jlr.R800061-JLR200, PMID: 19023137

Mavrich TN, Jiang C, Ioshikhes IP, Li X, Venters BJ, Zanton SJ, Tomsho LP, Qi J, Glaser RL, Schuster SC, Gilmour DS, Albert I, Pugh BF. 2008. Nucleosome organization in the Drosophila genome. Nature 453:358–62. DOI: 10.1038/nature06929

Meunier N, Belgacem YH, Martin JR. 2007. Regulation of feeding behaviour and locomotor activity by takeout in Drosophila. J Exp Biol 210:1424–34. DOI: 10.1242/jeb.02755

Moi D, Bernard C, Steinegger M, Nevers Y, Langleib M, Dessimoz C. 2023. Structural phylogenetics unravels the evolutionary diversification of communication systems in gram-positive bacteria and their viruses. DOI: 10.1101/2023.09.19.558401

Mueller GA, Edwards LL, Aloor JJ, Fessler MB, Glesner J, Pomés A, Chapman MD, London RE, Pedersen LC. 2010. The structure of the dust mite allergen Der p 7 reveals similarities to innate immune proteins. Journal of Allergy and Clinical Immunology 125:909–917.e4. DOI: 10.1016/j.jaci.2009.12.016

Nagnan-Le Meillour P. 2000. Chemosensory Proteins from the Proboscis of Mamestra brassicae. Chemical Senses 25:541–553. DOI: 10.1093/chemse/25.5.541

Naresh Singh R, Nayak SV. 1985. Fine structure and primary sensory projections of sensilla on the maxillary palp of Drosophila melanogaster Meigen (Diptera : Drosophilidae). International Journal of Insect Morphology and Embryology 14:291–306. DOI: 10.1016/0020-7322(85)90044-3

Nayak SV, Naresh Singh R. 1983. Sensilla on the tarsal segments and mouthparts of adult Drosophila melanogaster meigen (Diptera : Drosophilidae). International Journal of Insect Morphology and Embryology 12:273–291. DOI: 10.1016/0020-7322(83)90023-5

Öztürk-Çolak A, Marygold SJ, Antonazzo G, Attrill H, Goutte-Gattat D, Jenkins VK, Matthews BB, Millburn G, Dos Santos G, Tabone CJ, FlyBase Consortium, Perrimon N, Gelbart SR, Broll K, Crosby M, Dos Santos G, Falls K, Gramates LS, Jenkins VK, Longden I, Matthews BB, Seme J, Tabone CJ, Zhou P, Zytkovicz M, Brown N, Antonazzo G, Attrill H, Goutte-Gattat D, Larkin A, Marygold S, McLachlan A, Millburn G, Pilgrim C, Öztürk-Çolak A, Kaufman T, Calvi B, Campbell S, Goodman J, Strelets V, Thurmond J, Cripps R, Lovato T. 2024. FlyBase: updates to the *Drosophila* genes and genomes database. GENETICS 227:iyad211. DOI: 10.1093/genetics/iyad211

Palli SR, Osir EO, Eng W, Boehm MF, Edwards M, Kulcsar P, Ujvary I, Hiruma K, Prestwich GD, Riddiford LM. 1990. Juvenile hormone receptors in insect larval epidermis: identification by photoaffinity labeling. Proceedings of the National Academy of Sciences of the United States of America 87:796–800. PMID: 11607060

Palli SR, Touhara K, Charles JP, Bonning BC, Atkinson JK, Trowell SC, Hiruma K, Goodman WG, Kyriakides T, Prestwich GD. 1994. A nuclear juvenile hormone-binding protein from larvae of Manduca sexta: a putative receptor for the metamorphic action of juvenile hormone. Proceedings of the National Academy of Sciences 91:6191–6195. DOI: 10.1073/pnas.91.13.6191

Park S-K, Shanbhag SR, Wang Q, Hasan G, Steinbrecht RA, Pikielny CW. 2000. Expression patterns of two putative odorant-binding proteins in the olfactory organs of Drosophila melanogaster have different implications for their functions. Cell and Tissue Research 300:181–192. DOI: 10.1007/s004410050059

Pelosi P, Iovinella I, Felicioli A, Dani FR. 2014. Soluble proteins of chemical communication: an overview across arthropods. Frontiers in Physiology 5. DOI: 10.3389/fphys.2014.00320

Pelosi P, Iovinella I, Zhu J, Wang G, Dani FR. 2018a. Beyond chemoreception: diverse tasks of soluble olfactory proteins in insects. Biological Reviews 93:184–200. DOI: 10.1111/brv.12339

Pelosi P, Zhu J, Knoll W. 2018b. From radioactive ligands to biosensors: binding methods with olfactory proteins. Applied Microbiology and Biotechnology 102:8213–8227. DOI: 10.1007/s00253-018-9253-5

Pelosi P, Zhu J, Knoll W. 2018c. From radioactive ligands to biosensors: binding methods with olfactory proteins. Appl Microbiol Biotechnol 102:8213–8227. DOI: 10.1007/s00253-018-9253-5

Prestwich GD, Wojtasek H, Lentz AJ, Rabinovich JM. 1996. Biochemistry of proteins that bind and metabolize juvenile hormones. Archives of Insect Biochemistry and Physiology 32:407–419. DOI: 10.1002/(SICI)1520-6327(1996)32:3/4%3C407::AID-ARCH13%3E3.0.CO;2-G

Qiu X, Mistry A, Ammirati MJ, Chrunyk BA, Clark RW, Cong Y, Culp JS, Danley DE, Freeman TB, Geoghegan KF, Griffor MC, Hawrylik SJ, Hayward CM, Hensley P, Hoth LR, Karam GA, Lira ME, Lloyd DB, McGrath KM, Stutzman-Engwall KJ, Subashi AK, Subashi TA, Thompson JF, Wang IK, Zhao H, Seddon AP. 2007. Crystal structure of cholesteryl ester transfer protein reveals a long tunnel and four bound lipid molecules. Nat Struct Mol Biol 14:106–13. DOI: 10.1038/nsmb1197

Reese MG, Eeckman FH, Kulp D, Haussler D. 1997. Improved Splice Site Detection in Genie. Journal of Computational Biology 4:311–323. DOI: 10.1089/cmb.1997.4.311

Regnault-Roger C, Vincent C, Arnason JT. 2012. Essential Oils in Insect Control: Low-Risk Products in a High-Stakes World. Annual Review of Entomology 57:405–424. DOI: 10.1146/annurev-ento-120710-100554

Rezával C, Werbajh S, Ceriani MF. 2007. Neuronal death in *Drosophila* triggered by GAL4 accumulation. European Journal of Neuroscience 25:683–694. DOI: 10.1111/j.1460-9568.2007.05317.x

Rihani K, Ferveur J-F, Briand L. 2021. The 40-Year Mystery of Insect Odorant-Binding Proteins. Biomolecules 11:509. DOI: 10.3390/biom11040509

Robertson HM. 2019. Molecular Evolution of the Major Arthropod Chemoreceptor Gene Families. Annual Review of Entomology 64:227–242. DOI: 10.1146/annurev-ento-020117-043322

Rosenkranz M, Chen Y, Zhu P, Vlot AC. 2021. Volatile terpenes – mediators of plant-to-plant communication. The Plant Journal 108:617–631. DOI: 10.1111/tpj.15453

Rost B. 1997. Protein structures sustain evolutionary drift. Folding and Design 2:S19–S24. DOI: 10.1016/S1359-0278(97)00059-X

Rund SS, Bonar NA, Champion MM, Ghazi JP, Houk CM, Leming MT, Syed Z, Duffield GE. 2013. Daily rhythms in antennal protein and olfactory sensitivity in the malaria mosquito Anopheles gambiae. Sci Rep 3:2494. DOI: 10.1038/srep02494

Saito K, Su ZH, Emi A, Mita K, Takeda M, Fujiwara Y. 2006. Cloning and expression analysis of takeout/JHBP family genes of silkworm, Bombyx mori. Insect Mol Biol 15:245–51. DOI: 10.1111/j.1365-2583.2006.00612.x

Saitou N, Nei M. 1987. The neighbor-joining method: a new method for reconstructing phylogenetic trees. Molecular Biology and Evolution 4:406–425. DOI: 10.1093/oxfordjournals.molbev.a040454, PMID: 3447015

Sarov-Blat L, So WV, Liu L, Rosbash M. 2000. The Drosophila takeout gene is a novel molecular link between circadian rhythms and feeding behavior. Cell 101:647–56.

Saurabh S, Vanaphan N, Wen W, Dauwalder B. 2018. High functional conservation of takeout family members in a courtship model system. PLOS ONE 13:e0204615. DOI: 10.1371/journal.pone.0204615

Scheuermann EA, Smith DP. 2019. Odor-Specific Deactivation Defects in a *Drosophila* Odorant-Binding Protein Mutant. Genetics 213:897–909. DOI: 10.1534/genetics.119.302629

Schindelin J, Arganda-Carreras I, Frise E, Kaynig V, Longair M, Pietzsch T, Preibisch S, Rueden C, Saalfeld S, Schmid B, Tinevez J-Y, White DJ, Hartenstein V, Eliceiri K, Tomancak P, Cardona A. 2012. Fiji: an open-source platform for biological-image analysis. Nature Methods 9:676–682. DOI: 10.1038/nmeth.2019

Schmidt HR, Benton R. 2020. Molecular mechanisms of olfactory detection in insects: beyond receptors. Open Biology 10:200252. DOI: 10.1098/rsob.200252

Shanbhag, S.-K. P, C. P, R. S. 2001. Gustatory organs of Drosophila melanogaster : fine structure and expression of the putative odorant-binding protein PBPRP2. Cell and Tissue Research 304:423–437. DOI: 10.1007/s004410100388

Shanbhag SR. 1999. Atlas of olfactory organs of Drosophila melanogaster 1. Types, external organization, innervation and distribution of olfactory sensilla. International Journal of Insect Morphology and Embryology.

Shanbhag SR, Hekmat-Scafe D, Kim M -S., Park S -K., Carlson JR, Pikielny C, Smith DP, Steinbrecht RA. 2001a. Expression mosaic of odorant-binding proteins in *Drosophila* olfactory organs. Microscopy Research and Technique 55:297–306. DOI: 10.1002/jemt.1179

Shanbhag SR, MuÈller B, Steinbrecht RA. 2000. Atlas of olfactory organs of Drosophila melanogaster 2. Internal organization and cellular architecture of olfactory sensilla. Arthropod Structure.

Shanbhag SR, Singh K, Singh RN. 1995. Fine structure and primary sensory projections of sensilla located in the sacculus of the antenna of Drosophilamelanogaster. Cell Tissue Res 237–249.

Shanbhag SR, S.-K. P, C. P, R. S. 2001b. Gustatory organs of Drosophila melanogaster : fine structure and expression of the putative odorant-binding protein PBPRP2. Cell and Tissue Research 304:423–437. DOI: 10.1007/s004410100388

Shin S-W, Jeon J-H, Kim J-A, Park D-S, Shin Y-J, Oh H-W. 2022. Inducible Expression of Several Drosophila melanogaster Genes Encoding Juvenile Hormone Binding Proteins by a Plant Diterpene Secondary Metabolite, Methyl Lucidone. Insects 13:420. DOI: 10.3390/insects13050420

Silver L. 2001. Evolution of Gene Families. Encyclopedia of Genetics. Elsevier. p. 666–669. DOI: 10.1006/rwgn.2001.0433

So WV, Sarov-Blat L, Kotarski CK, McDonald MJ, Allada R, Rosbash M. 2000. *takeout*, a Novel *Drosophila* Gene under Circadian Clock Transcriptional Regulation. Molecular and Cellular Biology 20:6935–6944. DOI: 10.1128/MCB.20.18.6935-6944.2000

Stahl E, Hilfiker O, Reymond P. 2018. Plant–arthropod interactions: who is the winner? The Plant Journal 93:703–728. DOI: 10.1111/tpj.13773

Steinberg TH. 2009. Chapter 31 Protein Gel Staining Methods. Methods in Enzymology. Elsevier. p. 541–563. DOI: 10.1016/S0076-6879(09)63031-7

Steinberg TH, Jones LJ, Haugland RP, Singer VL. 1996. SYPRO Orange and SYPRO Red Protein Gel Stains: One-Step Fluorescent Staining of Denaturing Gels for Detection of Nanogram Levels of Protein. Analytical Biochemistry 239:223–237. DOI: 10.1006/abio.1996.0319

Steinbrecht RA. 1998. Odorant-Binding Proteins: Expression and Function. Annals of the New York Academy of Sciences 855:323–332. DOI: 10.1111/j.1749-6632.1998.tb10591.x

Steinbrecht RA. 1997. Pore structures in insect olfactory sensilla: A review of data and concepts. International Journal of Insect Morphology and Embryology 26:229–245. DOI: 10.1016/S0020-7322(97)00024-X

Steinbrecht RA, Mueller B. 1971. On the stimulus conducting structures in insect olfactory receptors. Zeitschrift für Zellforschung und Mikroskopische Anatomie 117:570–575. DOI: 10.1007/BF00330716

Steinbrecht RA, Ozaki M, Ziegelberger G. 1992. Immunocytochemical localization of pheromone-binding protein in moth antennae. Cell & Tissue Research 270:287–302. DOI: 10.1007/BF00328015

Stocker RF. 1994. The organization of the chemosensory system in Drosophila melanogaster: a rewiew. Cell and Tissue Research 275:3–26. DOI: 10.1007/BF00305372

Sugahara R, Tsuchiya W, Yamazaki T, Tanaka S, Shiotsuki T. 2020. Recombinant yellow protein of the takeout family and albino-related takeout protein specifically bind to lutein in the desert locust. Biochemical and Biophysical Research Communications 522:876–880. DOI: 10.1016/j.bbrc.2019.11.113

Sun JS, Xiao S, Carlson JR. 2018. The diverse small proteins called odorant-binding proteins. Open Biology 8:180208. DOI: 10.1098/rsob.180208, PMID: 30977439

Suzuki R, Fujimoto Z, Shiotsuki T, Tsuchiya W, Momma M, Tase A, Miyazawa M, Yamazaki T. 2011. Structural mechanism of JH delivery in hemolymph by JHBP of silkworm, Bombyx mori. Sci Rep 1:133. DOI: 10.1038/srep00133

Tan J, Zaremska V, Lim S, Knoll W, Pelosi P. 2020. Probe-dependence of competitive fluorescent ligand binding assays to odorant-binding proteins. Anal Bioanal Chem 412:547–554. DOI: 10.1007/s00216-019-02309-9

Tavares M, da Silva MRM, de Oliveira de Siqueira LB, Rodrigues RAS, Bodjolle-d’Almeida L, Dos Santos EP, Ricci-Júnior E. 2018. Trends in insect repellent formulations: A review. International Journal of Pharmaceutics **539**:190–209. DOI: 10.1016/j.ijpharm.2018.01.046, PMID: 29410208

Teufel F, Almagro Armenteros JJ, Johansen AR, Gíslason MH, Pihl SI, Tsirigos KD, Winther O, Brunak S, von Heijne G, Nielsen H. 2022. SignalP 6.0 predicts all five types of signal peptides using protein language models. Nature Biotechnology 40:1023–1025. DOI: 10.1038/s41587-021-01156-3

Thompson JD, Higgins DG, Gibson TJ. 1994. CLUSTAL W: improving the sensitivity of progressive multiple sequence alignment through sequence weighting, position-specific gap penalties and weight matrix choice.

Toulmay A, Prinz WA. 2011. A conserved membrane-binding domain targets proteins to organelle contact sites. Journal of Cell Science.

True JR, Carroll SB. 2002. Gene Co-Option in Physiological and Morphological Evolution. Annual Review of Cell and Developmental Biology 18:53–80. DOI: 10.1146/annurev.cellbio.18.020402.140619

Tsuchihara K, Fujikawa K, Ishiguro M, Yamada T, Tada C, Ozaki K, Ozaki M. 2005. An odorant-binding protein facilitates odorant transfer from air to hydrophilic surroundings in the blowfly. Chem Senses 30:559–64. DOI: 10.1093/chemse/bji049

Turina A del V, Nolan MV, Zygadlo JA, Perillo MA. 2006. Natural terpenes: Self-assembly and membrane partitioning. Biophysical Chemistry 122:101–113. DOI: 10.1016/j.bpc.2006.02.007

Unsicker SB, Kunert G, Gershenzon J. 2009. Protective perfumes: the role of vegetative volatiles in plant defense against herbivores. Current Opinion in Plant Biology 12:479–485. DOI: 10.1016/j.pbi.2009.04.001

van Kempen M, Kim SS, Tumescheit C, Mirdita M, Lee J, Gilchrist CLM, Söding J, Steinegger M. 2024. Fast and accurate protein structure search with Foldseek. Nature Biotechnology 42:243–246. DOI: 10.1038/s41587-023-01773-0, PMID: 37156916

Vanaphan N, Dauwalder B, Zufall RA. 2012. Diversification of takeout, a male-biased gene family in Drosophila. Gene 491:142–8. DOI: 10.1016/j.gene.2011.10.003

Venken KJT, Schulze KL, Haelterman NA, Pan H, He Y, Evans-Holm M, Carlson JW, Levis RW, Spradling AC, Hoskins RA, Bellen HJ. 2011. MiMIC: a highly versatile transposon insertion resource for engineering Drosophila melanogaster genes. Nature Methods 8:737–743. DOI: 10.1038/nmeth.1662, PMID: 21985007

Vermunt AMW, Kamimura M, Hirai M, Kiuchi M, Shiotsuki T. 2001. The juvenile hormone binding protein of silkworm haemolymph: gene and functional analysis. Insect Molecular Biology 10:147–154. DOI: 10.1046/j.1365-2583.2001.00249.x

Villegas G, Pereira MT, Love CR, Edery I. 2024. DAYWAKE implicates novel roles for circulating lipid-binding proteins as extracerebral regulators of daytime wake–sleep behavior. FEBS Letters 598:321–330. DOI: 10.1002/1873-3468.14789

Visser JH. 1986. Host Odor Perception in Phytophagous Insects. 121–44.

Vogt RG, Riddiford LM. 1981. Pheromone binding and inactivation by moth antennae. Nature 293:161–163. DOI: 10.1038/293161a0, PMID: 18074618

Wei X, Henke VG, Strübing C, Brown EB, Clapham DE. 2003. Real-Time Imaging of Nuclear Permeation by EGFP in Single Intact Cells. Biophysical Journal 84:1317–1327. DOI: 10.1016/S0006-3495(03)74947-9

Weiss J. 2003. Bactericidal/permeability-increasing protein (BPI) and lipopolysaccharide-binding protein (LBP): structure, function and regulation in host defence against Gram-negative bacteria. Biochemical Society Transactions 31:785–790. DOI: 10.1042/bst0310785

Wong LH, Gatta AT, Levine TP. 2019. Lipid transfer proteins: the lipid commute via shuttles, bridges and tubes. Nature Reviews Molecular Cell Biology 20:85–101. DOI: 10.1038/s41580-018-0071-5

Wong LH, Levine TP. 2017. Tubular lipid binding proteins (TULIPs) growing everywhere. Biochim Biophys Acta Mol Cell Res 1864:1439–1449. DOI: 10.1016/j.bbamcr.2017.05.019

Wu T, Hornsby M, Zhu L, Yu JC, Shokat KM, Gestwicki JE. 2023. Protocol for performing and optimizing differential scanning fluorimetry experiments. STAR Protocols 4:102688. DOI: 10.1016/j.xpro.2023.102688

Wybrandt GB, Andersen SO. 2001. Purification and sequence determination of a yellow protein from sexually mature males of the desert locust, Schistocerca gregaria. Insect Biochemistry and Molecular Biology 31:1183–1189. DOI: 10.1016/S0965-1748(01)00064-9

Xiao S, Sun JS, Carlson JR. 2019. Robust olfactory responses in the absence of odorant binding proteins. eLife 8:e51040. DOI: 10.7554/eLife.51040

Xu H, Turlings TCJ. 2018. Plant Volatiles as Mate-Finding Cues for Insects. Trends in Plant Science 23:100–111. DOI: 10.1016/j.tplants.2017.11.004

Yoshizawa Y, Sato R, Tsuchihara K, Ozaki K, Mita K, Asaoka K, Taniai K. 2011. Ligand carrier protein genes expressed in larval chemosensory organs of Bombyx mori. Insect Biochem Mol Biol 41:545–62. DOI: 10.1016/j.ibmb.2011.03.006

Zhu J, Guo M, Ban L, Song L-M, Liu Y, Pelosi P, Wang G. 2018. Niemann-Pick C2 Proteins: A New Function for an Old Family. Frontiers in Physiology 9:52. DOI: 10.3389/fphys.2018.00052

